# Leveraging Fungal Calcineurin-Inhibitor Structures, Biophysics and Dynamics to Design Selective and Non-Immunosuppressive FK506 Analogs

**DOI:** 10.1101/2020.04.14.039800

**Authors:** Sophie M.-C. Gobeil, Benjamin G. Bobay, Praveen R. Juvvadi, D. Christopher Cole, Joseph Heitman, William J. Steinbach, Ronald A. Venters, Leonard D. Spicer

## Abstract

Calcineurin is a critical enzyme in fungal pathogenesis and antifungal drug tolerance and, therefore, an attractive antifungal target. Current clinically-accessible calcineurin inhibitors, such as FK506, are immunosuppressive to humans, so exploiting calcineurin inhibition as an antifungal strategy necessitates fungal-specificity in order to avoid inhibiting the human pathway. Harnessing fungal calcineurin-inhibitor crystal structures, we recently developed a less immunosuppressive FK506 analog, APX879, with broad-spectrum antifungal activity and demonstrable efficacy in a murine model of invasive fungal infection. Our overarching goal is to better understand, at a molecular level, the interaction determinants of the human and fungal FK506-binding proteins (FKBP12) required for calcineurin inhibition in order to guide the design of fungal-selective, non-immunosuppressive FK506 analogs. To this end, we characterized high-resolution structures of the *M. circinelloides* FKBP12 bound to FK506, and of the *A. fumigatus, M. circinelloides* and human FKBP12 proteins bound to the FK506 analog, APX879, which exhibits enhanced selectivity for fungal pathogens. Combining structural, genetic and biophysical methodologies with molecular dynamics simulations, we identify critical variations in these structurally similar FKBP12-ligand complexes that will guide the rational design of inhibitors with enhanced fungal-selectivity.

**Significance statement:** Invasive fungal infections are a leading cause of death in the immunocompromised patient population. The rise in drug resistance to current antifungals highlights the urgent need to develop more efficacious and highly selective agents. Numerous investigations of major fungal pathogens have confirmed the critical role of the calcineurin pathway for fungal virulence, making it an attractive target for antifungal development. Although FK506 inhibits calcineurin, it is immunosuppressive in humans and cannot be used as an antifungal. By combining structural, genetic, biophysical, and *in silico* methodologies, we pinpoint regions of FK506 and a less immunosuppressive analog, APX879, that could be altered to enhance fungal selectivity. This work represents a significant advancement toward realizing calcineurin as a viable target for antifungal drug discovery.

---

Invasive fungal infections are a leading cause of death in immunocompromised patients. More than 1.6 million people die annually of infections caused by the major fungal pathogenic species of *Aspergillus, Candida, Cryptococcus*, and *Mucorales*(1). Due to rapidly emerging drug resistance to existing antifungals targeting the fungal cell wall and membrane, there is an urgent need to design more efficacious and highly selective antifungals targeting other critical fungal cellular pathways. However, this poses a fundamental challenge as both fungi and humans are eukaryotes and share many orthologous proteins and pathways(2). Recent structure-based inhibitor binding studies on the fungal heat shock protein 90 (Hsp90) have demonstrated the feasibility of increasing fungal-selective targeting of Hsp90(3, 4).

Calcineurin, the target of the immunosuppressive macrocyclic drug FK506 (tacrolimus) and the cyclic peptide cyclosporin A (CsA), is a promising target for the development of effective antifungal drugs(5). Calcineurin plays central roles in fungal growth, pathogenesis, cellular stress responses, and drug tolerance/resistance(6). The calcineurin protein complex consists of a catalytic subunit, calcineurin A (CnA), and a regulatory subunit, calcineurin B (CnB)(7). The immunosuppressants first bind to their respective immunophilins, FKBP12 (12-kDa FK506 binding protein) and CypA (cyclophilin A), which subsequently bind to calcineurin in a groove between the CnA and CnB subunits. The immunophilin-immunosuppressant complexes inhibit calcineurin serine-threonine phosphatase activity blocking the dephosphorylation of downstream targets, such as the human nuclear transcription factor NFAT involved in T-cell activation and interleukin-2 transcription, and the fungal transcription factor Crz1 (NFAT homolog), implicated in virulence, stress response, and thermotolerance(8-12). In humans, this leads to potent immunosuppression and is critical in preventing graft rejection, but also precludes FK506 and CsA usage as antifungals in immunocompromised patients.

FKBP12 proteins are members of the FKBP PPIase (peptidyl-prolyl isomerase) superfamily and catalyze the *cis*-*trans* isomerization of proline imidic peptide bonds(13-20). Based on our recent characterization of the FKBP12-FK506 and calcineurin-FK506-FKBP12 crystal structures(21, 22), we synthesized an FK506 analog, APX879, modified with an acetohydrazine moiety on FK506-C22 (**Fig. 1A**). Based on our crystal structures, we proposed that APX879 would interact less favorably with the human FKBP12 (*h*FKBP12) His88 residue compared with the corresponding *A. fumigatus* FKBP12 (*Af*FKBP12) Phe88 residue thus, potentially, reducing immunosuppression(22). In fact, APX879 displayed a 71-fold reduced immunosuppressive activity compared to FK506 while maintaining broad-spectrum *in vitro* antifungal activity against a wide range of human pathogenic fungi(22). *In vivo* testing in a murine model confirmed reduced immunosuppressive activity, and efficacy in a Cryptococcal model of invasive fungal infection(22).

**Figure 1.**
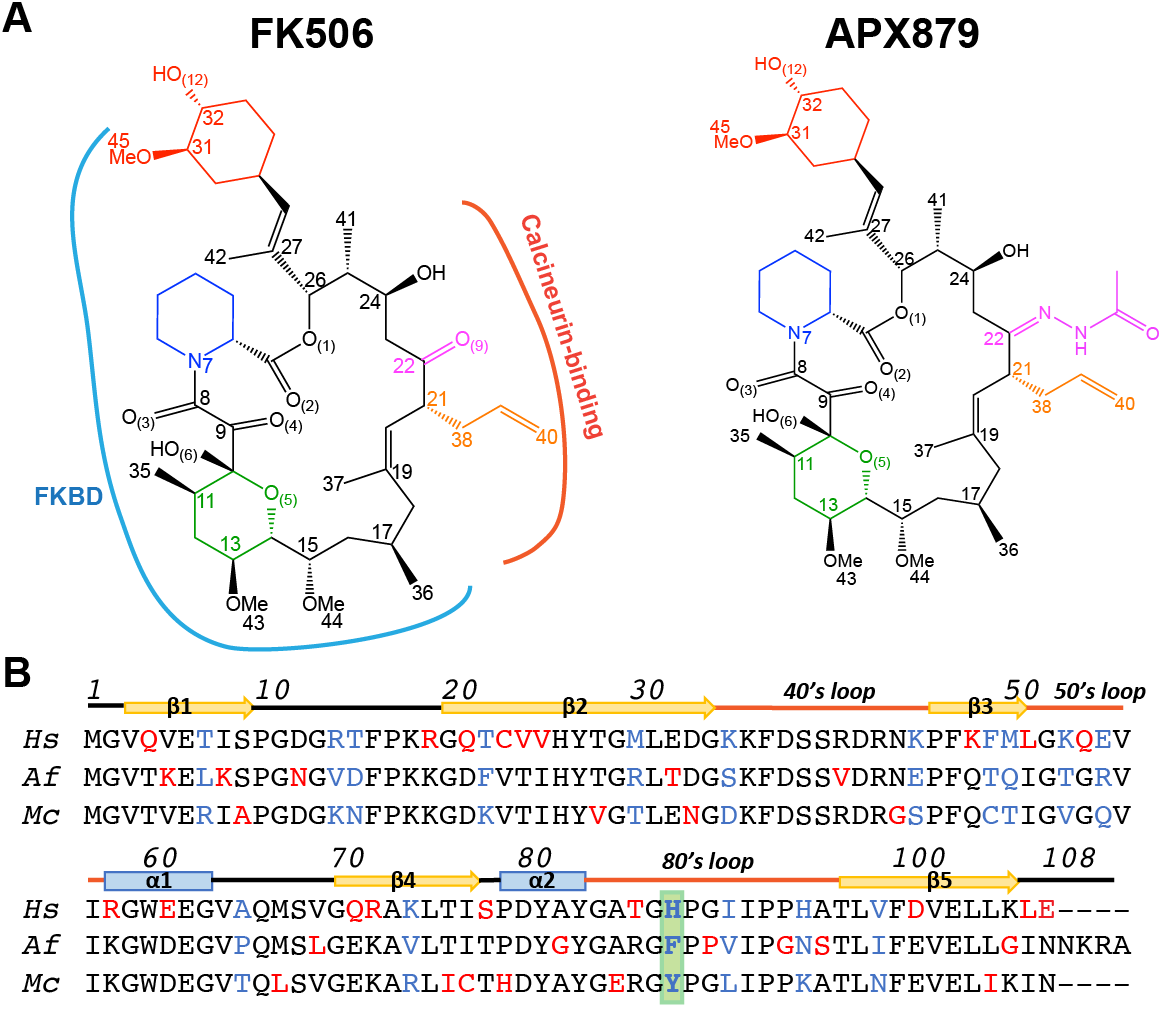
Chemical structure of FK506 and APX879 and sequence alignment of the human, *A. fumigatus* and *M. circinelloides* FKBP12 proteins. (A) APX879 is built on FK506 scaffold and incorporates an acetohydrazine moiety on FK506-C22. The cyclohexylidene (red), pipecolate (blue), pyranose rings (green), and the C21 allyl (orange) and C22 ketone/APX879 acetohydrazine moieties (magenta) are referenced according to FK506 atom numbering. The FKBP12 binding domain (FKBD; blue) and calcineurin binding interface (red) are indicated on FK506 (PDB 6TZ7). (B) Alignment of the human (*Hs*), *A. fumigatus (Af)*, and *M. circinelloides (Mc)* FKBP12 proteins. Residues are colored in red when varying in one of the sequences and in blue when different in all three sequences. Residue 88 is highlighted in green. Numbering and secondary structural elements are identified.

To guide the design of non-immunosuppressive FK506 analogs selectively targeting fungal calcineurin, here we quantitatively compared FK506 and APX879 binding to the human and fungal FKBP12 proteins from *A. fumigatus* and *M. circinelloides* through a combination of genetic, structural, and biophysical approaches. As *M. circinelloides* is an emerging human pathogen recalcitrant to many current antifungals, efficient targeting of calcineurin aided by the molecular characterization of the FKBP12 protein (*Mc*FKBP12) is of utmost importance(23, 24). Here, we report the first high-resolution crystal structures of *Mc*FKBP12 bound to FK506 in addition to the *h*FKBP12, *Af*FKBP12, and *Mc*FKBP12 proteins bound to APX879(22). Through genetic studies, we demonstrate that *Mc*FKBP12 does not functionally complement *Af*FKBP12 and reveal key requirements of FKBP12 residue 88 for “productive” binding and inhibition of calcineurin. While FK506 binds to both human and fungal FKBP12 proteins with high affinity (2 to 5 nM), APX879 binds 40- to 100-fold less tightly (120 to 450 nM). Strikingly, the human and fungal FKBP12 proteins show different responses to APX879 binding as observed by NMR, isothermal calorimetric titration (ITC) experiments, and molecular dynamic (MD) simulations that are not readily apparent in the static X-ray crystal structures. Furthermore, MD simulations allowed quantitative comparison of the significance of the FKBP12-ligand interactions between the human and fungal proteins. This analysis reveals regions of the ligands that could be altered to enhance selectivity toward the fungal FKBP12 proteins. Our approach highlights the potential of combining structural, genetic, biophysical, and *in silico* methodologies to fully describe and identify variation in protein-ligand interactions involving structurally similar proteins. We take a rational approach to understand the balance between the immunosuppressive and antifungal activities of FK506 and a new, less immunosuppressive FK506 analog, APX879, in an attempt to broaden the therapeutic window for the development of efficacious antifungals.

## Results

### Crystal structure of M. circinelloides FKBP12 bound to FK506

*Mc*FKBP12 shares 58% and 65% sequence identity with *h*FKBP12 and *Af*FKBP12, respectively. Sequence variations to *h*FKBP12 are located in: (i) β2, back wall of the FK506-binding pocket leading into the 40’s loop, (ii) the 50’s loop, and (iii) β4, leading into the 80’s loop (**Figs. 1B** and **2A**). Crystal structures of the apo and FK506-bound *h*FKBP12 and *Af*FKBP12 proteins have been reported(21, 25, 26). To correlate sequence variations with structure and identify potential differences between *Mc*FKBP12 and other FKBP12 proteins, we attempted to crystallize *Mc*FKBP12 in its apo form. All attempts failed to yield protein crystals hinting at potentially high conformational flexibility of *Mc*FKBP12. However, crystals were obtained with FK506 bound (2.5 Å, *P3*_*2*_*21*) suggesting rigidification of *Mc*FKBP12 by FK506 binding (**Table S1**).

**Figure 2.**
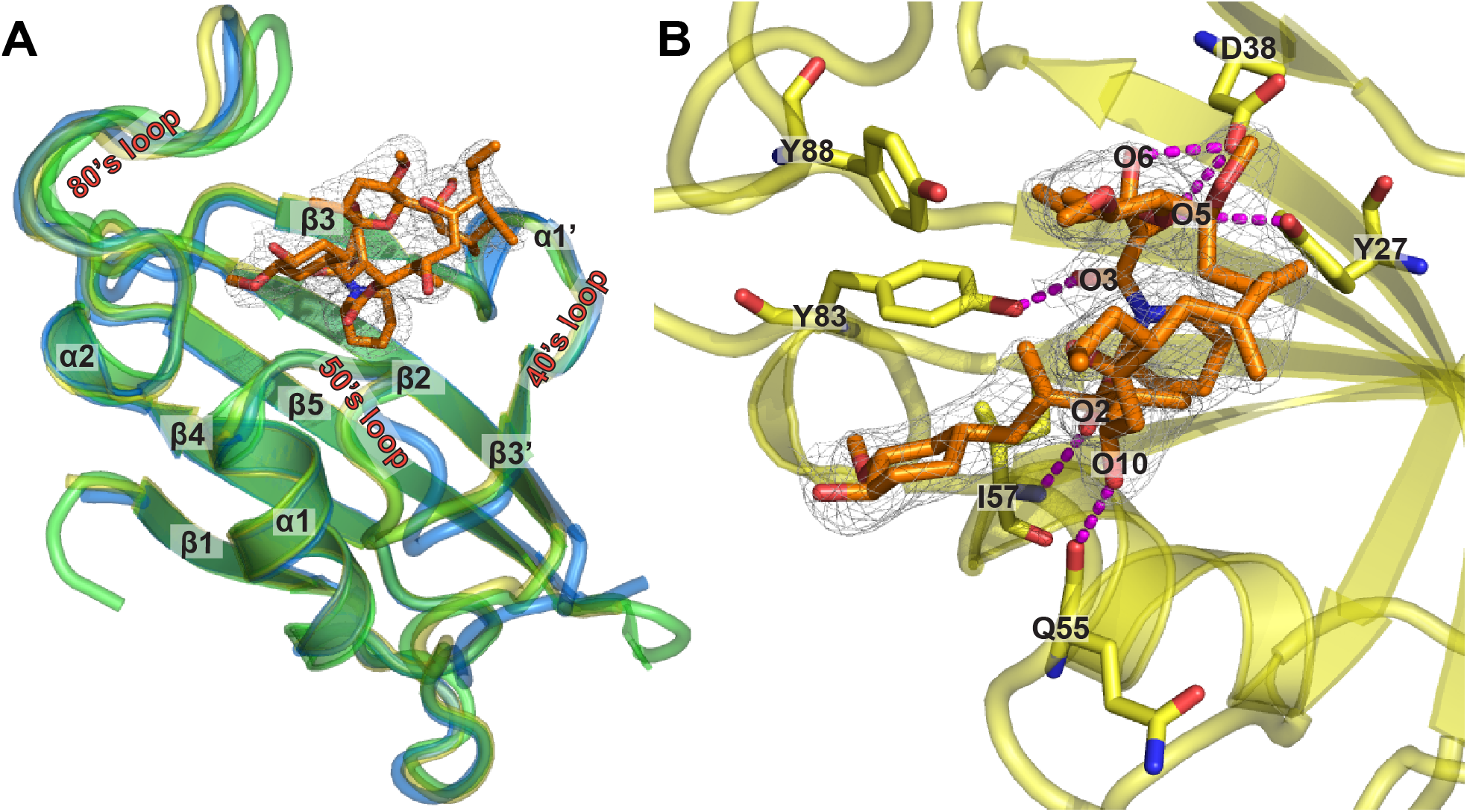
Crystal structure of *M. circinelloides* FKBP12 bound to FK506. (A) Overlay of the crystal structures of *Mc*FKBP12 (yellow; PDB ID 6VRX), *h*FKBP12 (blue; PDB ID 1FKJ) and *Af*FKBP12-P90G (green; PDB ID 5HWC) bound to FK506. Secondary structural elements are labeled and FK506 from the *Mc*FKBP12 crystal structure is shown in orange stick representation with the 2mFO-dFc density map at the 1 σ level. (B) FK506 binding pocket in *Mc*FKBP12. Five residues (Tyr27, Asp38, Gln55, Ile57, and Tyr83) are forming H-bonds to FK506 (in magenta dashed lines). See **Table S1** for Data collection and refinement statistics.

*Mc*FKBP12 shares the same fold as *h*FKBP12 and *Af*FKBP12 (Cα-RMSD ∼0.5 Å) with a 5-stranded β-sheet wrapping around an α-helix (**Fig. 2A**). The three main loops defining the FK506-binding cavity (40’s, 50’s, and 80’s loops) adopt the same conformation as observed in apo and FK506-bound forms of other FKBP12 proteins. When bound to *Mc*FKBP12, FK506 also adopts the same conformation as when bound to other mammalian and fungal FKBP12 proteins (FK506 RMSD ∼0.3 Å). Six hydrogen bonds (H-bonds) between FK506 and *Mc*FKBP12 are observed: 2 involving the backbone of residues Gln55 and Ile57, and 4 to the side chains of residues Tyr27, Asp38, and Tyr83 (**Fig. 2B**). Despite the sequence differences between *Mc*FKBP12 and *Af*FKBP12, they share high structural similarity, prompting investigation into their functional equivalence.

### M. circinelloides FKBP12 does not functionally complement A. fumigatus FKBP12 in calcineurin inhibition

The deletion of *A. fumigatus Af*FKBP12 encoding gene leads to FK506 resistance, establishing its central role for calcineurin inhibition(27). To verify if *Mc*FKBP12 can functionally complement *Af*FKBP12, an *A. fumigatus* strain expressing *Mc*FKBP12 was generated through genetic replacement of the *Affkbp12* native locus with *Mcfkbp12* (**Fig. S1**). Growth assays in the presence of increasing concentrations of FK506 revealed that *Mc*FKBP12 does not restore *A. fumigatus* FK506 sensitivity indicating that *Mc*FKBP12 may not bind or may bind but not inhibit *A. fumigatus* calcineurin *in vivo* (**Fig. 3A and S2**), consistent with previous analysis using *h*FKBP12(22). Using structure and sequence alignments, a single mutation of *h*FKBP12-His88 to Phe (*Af*FKBP12 identity), was shown to restore FK506 sensitivity(22). Interestingly, in *Mc*FKBP12 residue 88 is a Tyr, potentially sterically hindering the ternary complex formation with *A. fumigatus* calcineurin. To test this hypothesis, we generated a *Mc*FKBP12-Y88F mutant and confirmed restoration of FK506 sensitivity in *A. fumigatus* (**Fig. 3A, S1** and **S2**). We also verified the calcineurin-FK506-FKBP12 complex formation *in vivo* using GFP tagged *Mc*FKBP12 and *Mc*FKBP12-Y88F expression constructs by fluorescence microscopy (**Fig. 3B** and **S1**). In the absence of FK506, as observed with *Af*FKBP12, the *Mc*FKBP12 and *Mc*FKBP12-Y88F proteins localize in the cytoplasm and nuclei. Upon FK506 addition, both proteins localize to the septum as observed with native *Af*FKBP12, demonstrating the formation of the ternary complex with calcineurin at the hyphal septum, independent of the Y88F mutation. This highlights the central role of residue 88 in the formation of a “productive” inhibitory complex with *A. fumigatus* calcineurin. Despite the binding of *Mc*FKBP12 (and *h*FKBP12) to *A. fumigatus* calcineurin *in vivo* in the presence of FK506, subsequent inhibition is dependent on the presence of the critical Phe residue at position 88 suggesting that the side chain size (*Mc*FKBP12-Tyr88) and charge (*h*FKBP12-His88) control the formation of a “productive” inhibitory protein-ligand-protein interface.

**Figure 3.**
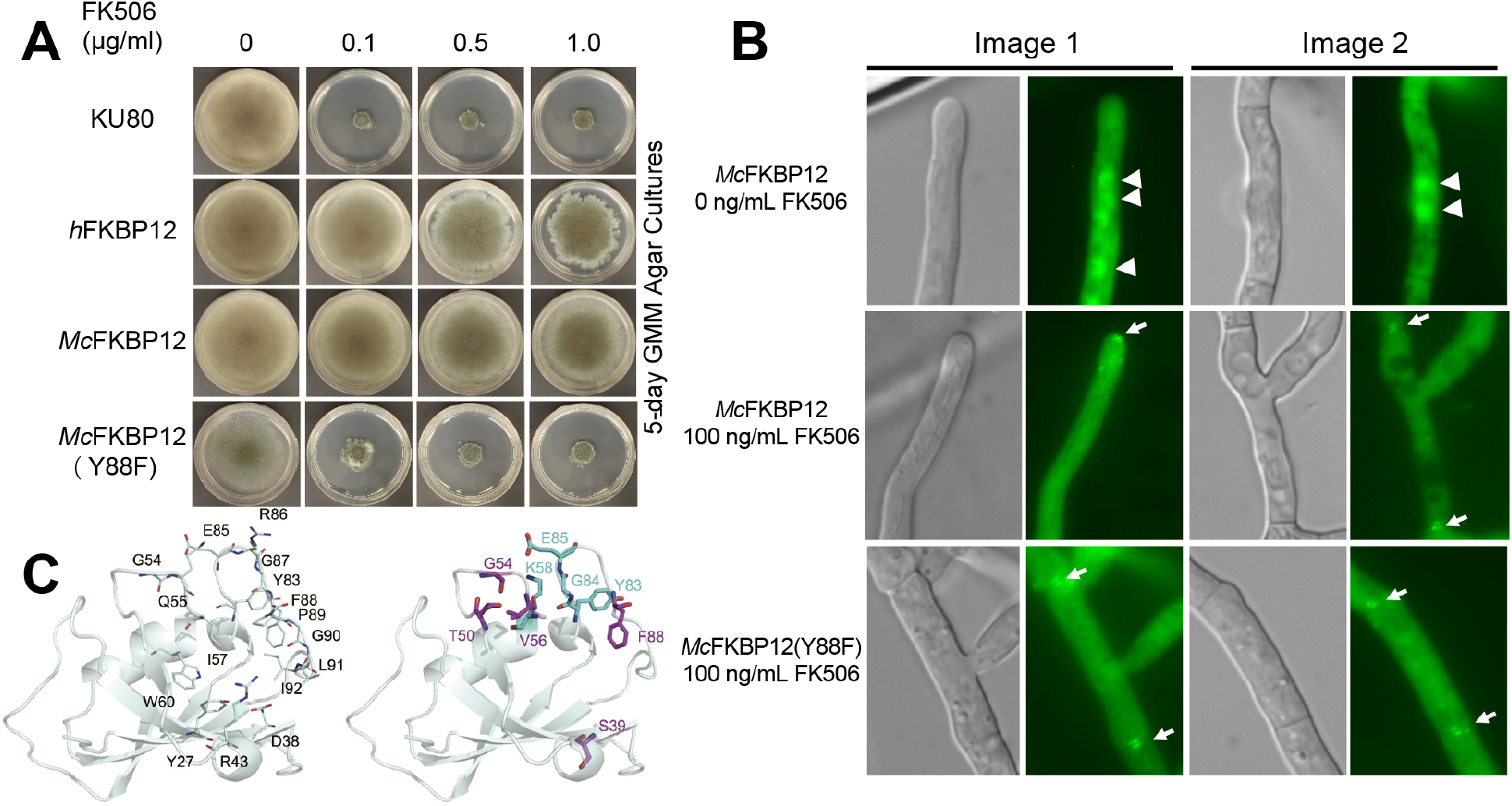
*M. circinelloides* FKBP12 does not functionally complement *A. fumigatus* FKBP12. (A) Growth of the wild-type *A. fumigatus* strain (KU80) and the strain expressing the *h*FKBP12, *Mc*FKBP12, and *Mc*FKBP12-Y88F proteins in the absence and presence of FK506 for 5-days at 37 °C. (B) Microscopic localization of the *Mc*FKBP12 and *Mc*FKBP12-Y88F proteins *in vivo* in the absence or presence of FK506. Arrowheads show nuclear localization of *Mc*FKBP12. Arrows indicate binding of *Mc*FKBP12 and *Mc*FKBP12-Y88F to *Af* calcineurin at the hyphal septum. (C) *Mc*FKBP12(Y88F)-FK506 structure (cartoon representation) with the position of common H-bonds (line representation) noted with labeled residues (left). The right image shows the H-bonds to FK506 gained (cyan) or lost (purple) due to the Tyr88Phe mutation.

To assess the structural implications of FK506 binding to the *Mc*FKBP12-Y88F protein, we performed 500 ns MD simulations on *Mc*FKBP12, *Mc*FKBP12-Y88F, and *Af*FKBP12 while bound to FK506 (**Fig. S3**). While these data suggest that the Y88F mutation in *Mc*FKBP12 does not grossly alter the conformational state, Cα−RMSF of *Mc*FKBP12, or average number of H-bonds between FKBP12 and FK506 (*Mc*FKBP12: 1.5, *Mc*FKBP12-Y88F: 1.7, *Af*FKBP12: 2.1), it does alter which residues of *Mc*FKBP12 and atoms of FK506 are forming H-bonds pairs.

H-bonds to FK506 involving residues 27, 38, 43, 54, 55, 57, 60, 83, 85, 86-88, and 91 are observed during the course of the MD simulation calculations for *Mc*FKBP12, *Mc*FKBP12-Y88F, and *Af*FKBP12 confirming the central role of the 50’s and 80’s loops in FK506 binding; correlating with observations noted in the crystal structures (**Fig. 3C**). Interestingly, the H-bonds between FK506 and *Mc*FKBP12-Y88F residues Ser39-OG, Thr50-OG1, Gly54-N, Val56-O, and Tyr83-OH are disrupted in comparison to *Mc*FKBP12 while new H-bonds involving Lys58, Tyr83, Gly84, and Glu85 backbone are observed. These latter two new H-bonds, implicating the 80’s loop, are also observed for *Af*FKBP12-FK506 during the MD simulation. This suggests that the Y88F mutation shifts the relative significance of the 50’s and 80’s loop in the interaction of *Mc*FKBP12 with FK506. These results emphasize the central role of *Af*FKBP12-Phe88 for the formation of a productive FKBP12-FK506 composite surface allowing for calcineurin inhibition.

### Crystal structures of the human and fungal FKBP12 proteins bound to APX879

The FK506 analog, APX879 substituted on the C22-ketone with an acetohydrazine moiety (**Fig. 1A**), was hypothesized, based on our structural data, to introduce a steric clash with *h*FKBP12-His88; potentially benefiting fungal-selectivity.(22) To understand the differential binding of APX879 to the fungal and human FKBP12 proteins, we obtained X-ray crystal structures of each of these complexes (**Fig. 4** and **Table S1**). Crystals of *h*FKBP12, *Af*FKBP12, and *Mc*FKBP12 bound to APX879 diffracted at resolutions of 1.7, 1.6, and 1.9 Å, respectively. The FKBP12-APX879 structures showed minimal structural variations compared to their FK506-bound counterparts (Cα-RMSDs ∼0.5 Å) maintaining similar H-bonds pattern (*i*.*e*. 4 to 5): two with the backbone of residues Glu_*h*_/Arg_*Af*_/Gln_*Mc*_55 and Ile57 and two/three with the side chain of residues Tyr27, Asp38, and Tyr83.

**Figure 4.**
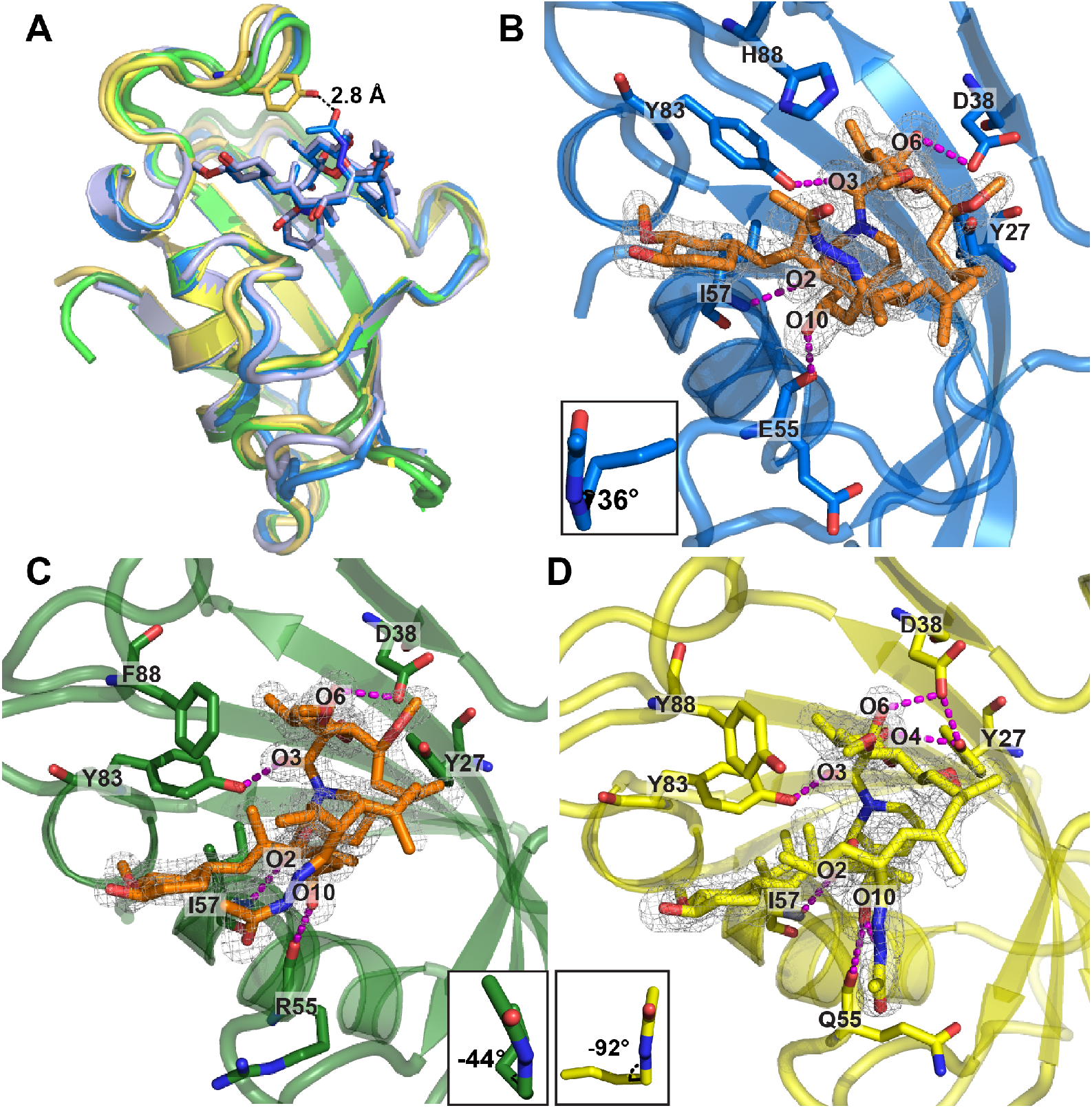
Crystal structures of the human, *A. fumigatus*, and *M. circinelloides* FKBP12 proteins bound to APX879. (A) Comparison of the overlaid crystal structures of *Mc*FKBP12-APX879 (yellow; PDB ID 6VCT), *Mc*FKBP12-FK506 (gold; PDB ID 6VRX), *h*FKBP12-APX879 (blue; PDB ID 6VCU), *h*FKBP12-FK506 (light blue; PDB ID 1FKJ), *Af*FKBP12-APX879 (dark green; PDB ID 6VCV), and *Af*FKBP12(P90G)-FK506 (light green; PDB ID 5HWC). APX879 and FK506 from the *h*FKBP12 crystal structures are represented in blue and light blue sticks. Distance to APX879-C60 and *Mc*FKBP12-Tyr88 was estimated at 2.9 Å. (B - D) Representation of APX879 (orange) in the 2mFO-dFc density map (at 1 σ level) in the binding cavity of (B) *h*FKBP12, (C) *Af*FKBP12, and (D) *Mc*FKBP12. The C21-C22 dihedral angle measured are illustrated in the lower corner. Residues (Tyr27, Asp38, E_*h*_/R_*Af*_/Q_*Mc*_55, Ile57, and Tyr83) are forming H-bonds (identified by magenta dashed lines) maintaining APX879 in the binding pockets. See **Table S1** for Data collection and refinement statistics.

Previous crystal structures of *Af*FKBP12 bound to FK506 required a Pro90Gly mutation in order to trap the ligand in the binding cavity. Without this mutation, an apo homodimer was captured that is hypothesized to result from a self-catalysis mechanism involving the Pro89-Pro90 motif(21). Here, we obtained crystals of *Af*FKBP12 bound to APX879 without the requirement for a P90G mutation by the addition of APX879 during purification. Interestingly, Pro90 adopted the *cis* conformation, as observed in the crystal structure of the calcineurin-FK506-FKBP12 complex (PDB 6TZ7)(22). In contrast, the human and *Mc*FKBP12 proteins, having a Pro89-Gly90 motif, do not show any evidence of an intermolecular self-catalysis-like binding event. The Pro90 *cis* conformer in *Af*FKBP12 allows the 80’s loop to reduce the size of the FK506-binding pocket (as estimated by 3V volume calculations - PDB 5HWB; *trans* conformer ∼390 Å^3^; *cis* conformer 240 Å^3^) to a volume similar to other FKBP12 proteins containing a Gly90 residue (*h*FKBP12 220 Å^3^, PDB 2PPN)(28). Using apo and FK506-bound *Af*FKBP12 NMR assignments, we confirmed the adoption of the Pro90 *cis* conformer in solution as suggested by the Cβ (apo: 33.2 ppm; FK506-bound: 34.2 ppm) and Cγ (apo: 24.2 ppm) resonances(29-31). Our *Af*FKBP12-APX879 crystal structure further emphasizes that the Pro90 *cis* conformation might modulate the binding cavity volume.

We next assessed APX879 acetohydrazine moiety accommodation by the different FKBP12 proteins. Similar to the *h*FKBP12-FK506 structure, in the *h*FKBP12-APX879 crystal structure a C21-C22 dihedral angle of 36° is measured positioning the acetohydrazine in the same orientation as the FK506-C22 ketone (**Fig. 4B**). In the *Af*FKBP12-APX879 crystal structure, two protein monomers bound to APX879 were resolved, one showing the acetohydrazine moiety in the same orientation as the FK506-C22 ketone (C21-C22 dihedral angle of 68°) and the other showing a 90° rotation of the acetohydrazine moiety with a C21-C22 dihedral angle of -44° (**Fig. 4C**). This suggests conformational flexibility that might be restrained upon calcineurin binding. Interestingly, when bound to *Mc*FKBP12, the APX879 acetohydrazine moiety adopts a third conformation with a C21-C22 dihedral angle of -92° leading to a ∼130° flip compared to FK506-C22 ketone (*Mc*FKBP12-FK506 C21-C22 dihedral angle ∼32°) (**Fig. 4D**). The adoption of this conformation prevents steric clashes with *Mc*FKBP12-Tyr88, positioned ∼3 Å away from APX879-C60 when overlaid with *Mc*FKBP12-FK506. Altogether, the crystal structures suggest that the flexibility of the acetohydrazine moiety compensates, at least in part, for the bulkiness of the amino acids at position 88. The *h*FKBP12-His88 does not by itself prevent APX879 binding but complex formation with calcineurin might add further conformational restraints on APX879. An overlay of the *A. fumigatus* and bovine calcineurin-FK506-FKBP12 crystal structures (PDB 6TZ7(22) and 1TCO(32)) with *h*FKBP12, *Af*FKBP12, and *Mc*FKBP12 bound to APX879 show that the conserved CnA residues Pro377 and Phe378 in the CnB binding helix (CnA-BBH) are less than 3 Å away from the APX879 acetohydrazine moiety, supporting the necessity for a rearrangement of either the ligand or the CnA-BBH in order to facilitate the formation of the calcineurin-APX879-FKBP12 complex.

### Biophysical characterization of the human versus fungal FKBP12-FK506/APX879 protein-ligand interaction

To identify biophysical variations between FK506 and APX879 interactions with the human and fungal FKBP12 proteins, we performed ITC assays to establish thermodynamic constants (**Table 1** and **S4**). As previously reported(33-37), *h*FKBP12 interacts with FK506 with high affinity (*K*_*d*_ 2.7 ± 0.5 nM). *Af*FKBP12 and *Mc*FKBP12 also interact tightly with FK506 (*K*_*d*_ 4.7 ± 0.6 and 3.0 ± 0.7 nM, respectively). The affinity for APX879 was reduced by ∼40-fold for *h*FKBP12 and *Mc*FKBP12 (*K*_*d*_ ∼120 nM) and 100-fold for *Af*FKBP12 (*K*_*d*_ = 462 ± 23 nM). The binding stoichiometry for *h*FKBP12 and *Mc*FKBP12 was 1:1 while, for *Af*FKBP12, a stoichiometry of 2:1 (ligand-FKBP12) was obtained for both FK506 and APX879 binding. This could be due to the displacement of a sparsely populated low-affinity homodimer or a shift in the Pro90 *cis*/*trans* equilibrium triggered by FK506/APX879 binding(21). All of the FKBP12 proteins showed similar free energy (*ΔG°*) for FK506 (∼ -12 kcal/mol) or APX879 (∼ -9 kcal/mol) binding. The fungal FKBP12 proteins demonstrated a 5 to 8 kcal/mol increase in enthalpy (*ΔH*) compared to *h*FKBP12 for FK506 binding while for APX879 they all shared a similar *ΔH* (−1.5 kcal/mol). This corresponds to an increase of 6 to 13 kcal/mol for APX879 binding compared to FK506. The entropic component contribution (*TΔS*) for FK506 binding is increased by 5 to 8 kcal/mol in the fungal FKBP12 proteins compared to *h*FKBP12. For APX879, all FKBP12 proteins showed a *TΔS* of ∼ 7 kcal/mol, corresponding to an 11 to 3 kcal/mol increase compared to FK506.

**Table 1.** Thermodynamic parameters for the binding of FK506 and APX879 to human, *Af* and *Mc* FKBP12 proteins determined by ITC. Average from triplicates performed at 25°C, See **Fig. S4** for heat pulse data.

| FK506 | $K_d$<br>(nM) | $\Delta H$<br>(kcal/mol) | $T\Delta S$<br>(kcal/mol) | Stoichiometry | $\Delta G^\circ$<br>(kcal/mol) |
| --- | --- | --- | --- | --- | --- |
| *h*FKBP12 | $2.7 \pm 0.5$ | $-15.0 \pm 1.1$ | $-3.3 \pm 1.2$ | $1.09 \pm 0.03$ | $-12.0 \pm 1.6$ |
| *Af*FKBP12 | $4.7 \pm 0.6$ | $-6.7 \pm 0.3$ | $4.6 \pm 0.3$ | $2.02 \pm 0.06$ | $-11.3 \pm 0.4$ |
| *Mc*FKBP12 | $3.0 \pm 0.7$ | $-9.8 \pm 0.1$ | $1.9 \pm 0.2$ | $1.06 \pm 0.06$ | $-11.7 \pm 0.2$ |
| APX879 | $K_d$<br>(nM) | $\Delta H$<br>(kcal/mol) | $T\Delta S$<br>(kcal/mol) | Stoichiometry | $\Delta G^\circ$<br>(kcal/mol) |
| *h*FKBP12 | $119.4 \pm 15.8$ | $-1.8 \pm 0.4$ | $7.4 \pm 0.8$ | $1.41 \pm 0.03$ | $-9.2 \pm 0.9$ |
| *Af*FKBP12 | $461.6 \pm 22.6$ | $-1.2 \pm 0.4$ | $7.2 \pm 1.0$ | $1.96 \pm 0.13$ | $-8.0 \pm 1.0$ |
| *Mc*FKBP12 | $126.4 \pm 30.3$ | $-2.0 \pm 0.1$ | $7.4 \pm 0.2$ | $0.93 \pm 0.17$ | $-9.4 \pm 0.2$ |

NMR titrations of FK506 and APX879 into *h*FKBP12, *Af*FKBP12, and *Mc*FKBP12, as followed by ^1^H-^15^N HSQC, allowed for the characterization, in solution and at the residue level, of the protein responses to ligand binding (**Fig. 5** and **S5**). Both FK506 and APX879 induced chemical shift perturbations at least one standard deviation larger than the average for residues Ile25, His26, Thr_*Af*_/Cys_*Mc*_49, Val56, Ile57, Val102, and Glu103 in the fungal FKBP12 proteins(31). Additionally, *Mc*FKBP12 Phe47, Gln48, and Glu101, and *Af*FKBP12 Tyr27 and Arg43 also showed significant chemical shift differences upon ligand binding. Strikingly, a different set of residues in *h*FKBP12 (Val25, Ser40, Arg43, Gly52, Ile57, Glu62, Ala65, and Phe100) are observed as having large chemical shift variations between the apo and FK506/APX879 bound forms; strongly indicating an altered binding orientation and differential binding determinants.

**Figure 5.**
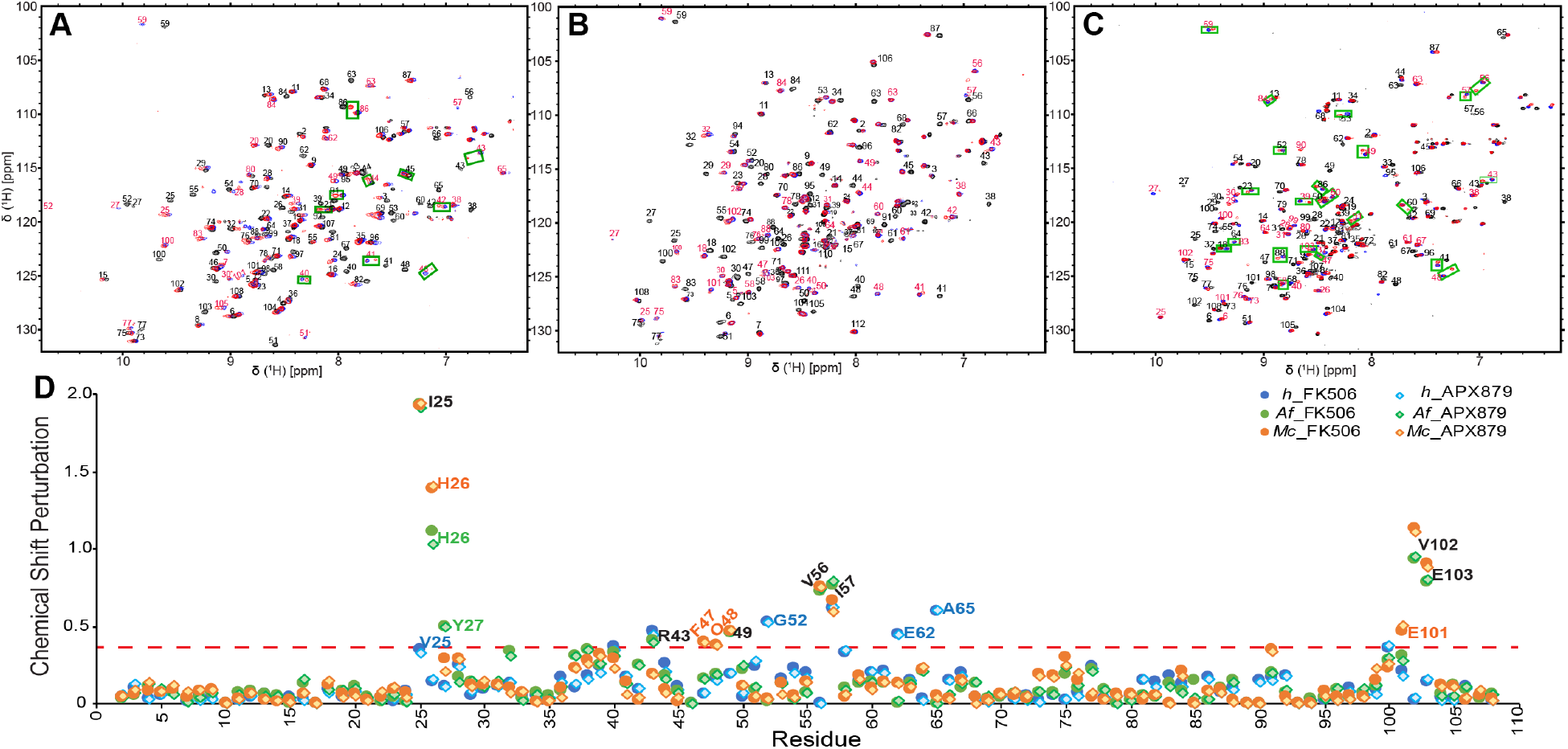
NMR binding of FK506 and APX879 to the human, *A. fumigatus*, and *M. circinelloides* FKBP12 proteins. Initial (0:1; in black) and final (2:1; in red) ^15^N-HSQC of the titration of APX879 with the (A) human, (B) *A. fumigatus* and (C) *M. circinelloides* FKBP12 proteins. The FK506 final titration point (2:1) with the respective proteins is shown in blue for reference. Peaks doubling when protein is fully bound to APX879 are indicated by green squares (See **Fig. S5** for full titration and zoomed regions where peak doubling are observed). (D) Protein chemical shift perturbation due to FK506 and APX879 binding. The red dashed line represents significant chemical shift perturbations (≥ protein mean + 1 S.D.).

The chemical exchange for both FK506 and APX879 binding occurs in the slow exchange regime (tight binding) for all three FKBP12 proteins leading, in most cases, to the observation of the chemical shifts of both the unbound and bound populations within the intermediate titration points (**Fig. S5**). Interestingly, peak doubling at the highest ligand:FKBP12 ratios (1:1 and 2:1) was observed for residues in the *h*FKBP12 40’s loop (40-45 and Lys48), 80’s loop (Thr86, Ile91, Ile92, and His95) and residues Glu62-Gly63 suggesting conformational dynamics on the NMR timescale for the bound protein. *Mc*FKBP12 also showed peak doubling for an increased number of residues located in the 40’s, 50’s and 80’s loop (Ser39, Arg41, Arg43, 48-50, 52-53, 56-60, 83-84, Arg86, Tyr88, Leu91, and Glu103) while none were observed in *Af*FKBP12. These observations suggest that APX879 binds tightly to all three FKBP12 proteins with an intermediate to slow exchange rate but that *h*FKBP12 and *Mc*FKBP12, when bound to APX879, experience more conformational flexibility than *Af*FKBP12 or when they are bound to FK506.

### MD simulations reveal structural differences relative to the X-ray crystal structures and suggest conformational flexibility

To relieve crystal contacts (**Table S2**) that might affect the protein and ligand conformation and to substantiate conformational flexibility suggested by NMR, six 500 ns MD simulations were performed for each protein-ligand complex (**Fig. S6**). Interactions as characterized by Cα-RMSD, the solvent accessible surface area, center of mass measurements, and H-bonds patterns were found to be generally similar between the different complexes (**Fig. S6** and **Table S3**). Contrastingly, the Cα-RMSF (atomic positions in the MD simulation fitted to the crystal structures) highlighted areas of the proteins and ligands exhibiting conformational differences (**Figs. S7** and **S8**). Both fungal FKBP12 proteins when bound to APX879 showed high RMSF deviations for residues 52-55 not observed for the *h*FKBP12-APX879 complex nor any of the FK506-bound counterparts suggestive of altered conformational states for these residues. The Cα-RMSF also revealed a common deviation of the 80’s loop for all FKBP12:ligand complexes. The core of the FK506 and APX879 macrocycles were found to be more similar when comparing the crystal structures and MD simulations (lower RMSF) than the solvent exposed portions of the molecules (spheres in **Fig. S7)**. For all FK506-bound complexes, the largest atomistic RMSF deviations occurred at atoms C40 (FK506-C21 allyl moiety) and C45 (cyclohexylidene ring C31-O-methyl) and to a lesser extent for atoms C33, C34, O11, and O12 of the cyclohexylidene ring that are directly involved in interactions with the 80’s loop and are also at the interface for calcineurin binding. For complexes bound to APX879, the ligand showed significant deviations at the acetohydrazine moiety. In addition, significant deviations were observed for an extended region surrounding the cyclohexylidene ring (atoms C28-C34, C42, and C45) and O8, and O10 when bound to *Mc*FKBP12 or *h*FKBP12. These data suggest that the crystallization process might have artificially favored the adoption of one conformer in otherwise flexible regions of the proteins and ligands.

### Significance of FKBP12 and FK506/APX879 contacts observed in the MD simulations

The significance of protein-ligand contacts observed throughout the MD simulations was quantified allowing for identification of variations between the human and fungal complexes potentially informing the design of fungal-selective FK506 analogs. In all FKBP12-ligand complexes a common core of conserved residues (Tyr27, Phe37, Asp38, Phe47, Val56, Ile57, and Trp60) and one non-conserved residue (E_*h*_/R_*Af*_/Q_*Mc*_55) made a similar number of contacts to FK506 or APX879 (**Figs. S9** and **S10**). In contrast, the 80’s loop residues showed a varying degree of interaction to the ligands; the fungal FKBP12 proteins had a significantly larger number of contacts involving this loop.

Z-score metrics allowed for the quantification of the relative significance of the contacts for the human and the fungal proteins when binding either FK506 or APX879 (**Fig. 6** and **S10**). Residues Tyr83 and His_*h*_/Phe_*Af*_/Tyr_*Mc*_88 shared an increased importance for FK506 and APX879 binding to both fungal FKBP12 proteins while residues Phe49 and Ile91 were more significant for *h*FKBP12. Interestingly, *h*FKBP12 also relied on Gly84 and Phe100 for FK506 binding while Arg43 was notably important for APX879 binding. Binding to FK506 also implicated residues Tyr27, Val56, and Trp60 as making differentiating contacts between *h*FKBP12 and *Af*FKBP12 and residue R_*h*_/T_*Mc*_86 as making differentiating contacts between *h*FKBP12 and *Mc*FKBP12 that are not involved in differentiating contacts when binding APX879. Z-score metrics also pointed to a common core of atoms from both ligands, approximating 65% of the molecules, demonstrate similar significance for binding to all three FKBP12 proteins (**Fig. 6** and **S10**). In the FK506-bound forms, atoms C2-C4 of the pipecolate ring and C45 (cyclohexylidene ring C31-O-methylation) are more significant for interaction with *h*FKBP12, while C35 (pyranose ring methylation) and C37 (C19-methylation) are more significant for interaction with the fungal FKBP12 proteins. Interestingly, for the APX879 bound forms, atoms of the cyclohexylidene ring (C33, C42, and O12), pipecolate ring (C3-C4), and C15-C19 region (C15-C16, C18, C36-C37, and O8) are more significant for interaction with *h*FKBP12, while atoms C11-C12 and C35 of the pyranose ring, C28, C30, and C45 of the opposite side of the cyclohexylidene ring, and C61 of the acetohydrazine moiety show more significant interactions with the fungal FKBP12 proteins. The complexes bound to APX879 show nearly identical importance of the acetohydrazine moiety and, therefore, no significant differences are noted within this region with the exception of C61 possibly making significant interactions with the fungal FKBP12 proteins (*Mc*FKBP12 Z-score right under +1, right above +1 for *Af*FKBP12).

**Figure 6.**
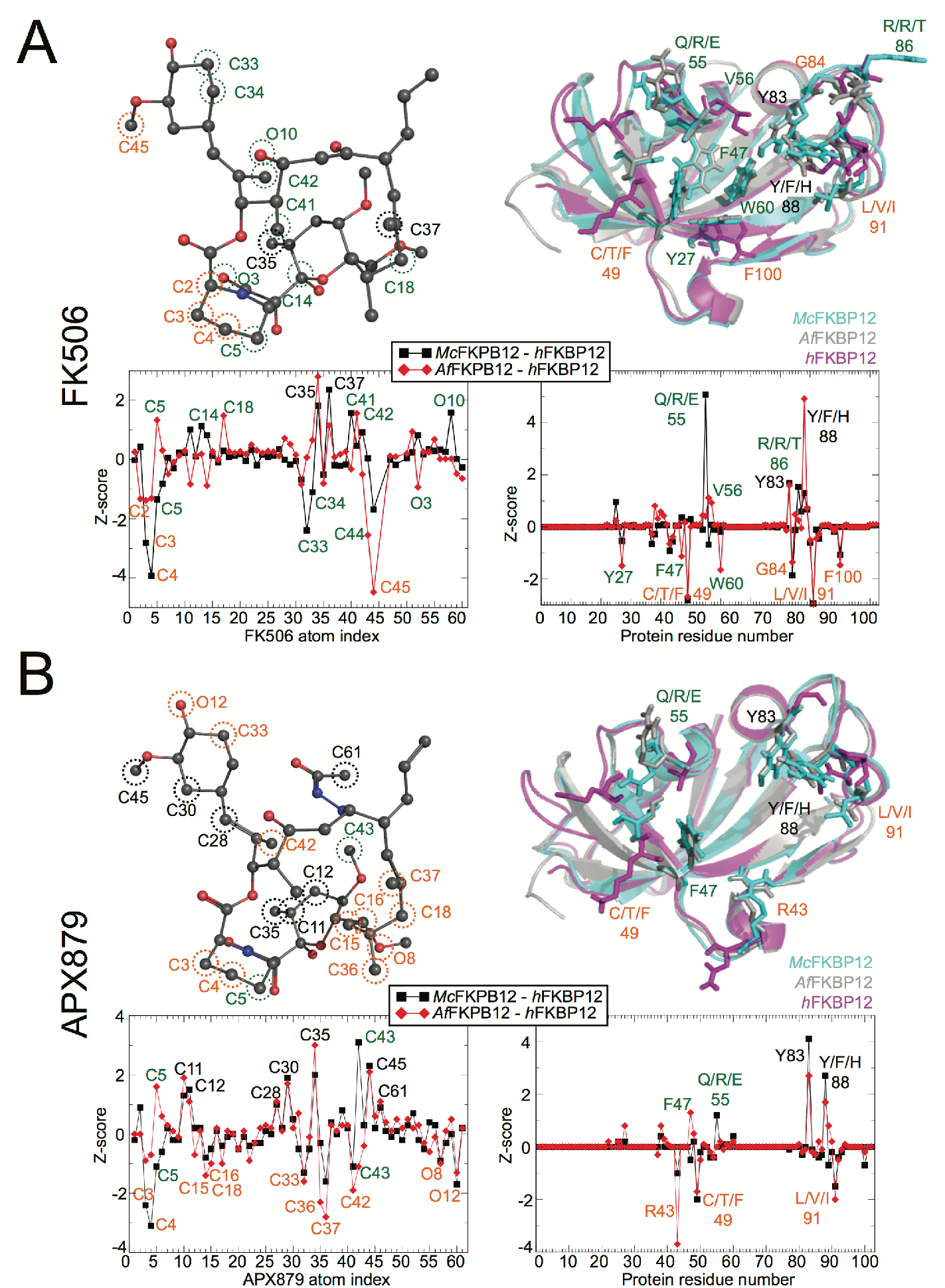
Ligand atoms and protein residue contacts observed in the MD simulations. Analysis of the significance of the observed contacts of *h*FKBP12, *Af*FKBP12, and *Mc*FKBP12 bound to (A) FK506 or (B) APX879. <u>Z-score graphs</u>: black squares represent Z-scores of *Mc*FKBP12 contacts minus *h*FKBP12 contacts, while red diamonds represent *Af*FKBP12 contacts minus *h*FKBP12 contacts. More significant contacts for the fungal proteins (Δ Z-score >1) are labeled in black while contacts more significant for the human protein (Δ Z-score <1) are labeled in orange. Significant contact differences between *Af*FKBP12 and *Mc*FKBP12 are labeled green. <u>Left side</u>: Stick and sphere representation of (A) FK506 and (B) APX879 colored accordingly to the atom type. Atoms making more significant contacts to *h*FKBP12 (orange labels), the fungal proteins (black labels), and differences between the fungal *Af*FKBP12 and *Mc*FKBP12 (green labels) are circled using the same coloring scheme. <u>Right side</u>: Residues of *h*FKBP12 (magenta), *Af*FKBP12 (gray), and *Mc*FKBP12 (cyan) making more significant interactions to the ligand (orange labels for *h*FKBP12, black labels for fungal proteins and green labels between the fungal *Af*FKBP12 and *Mc*FKBP12) are shown in stick format on the protein structures.

Overall, these observations indicate that the fungal FKBP12 proteins rely generally on the same amino acids (most notably Tyr83 and Phe88) to interact with FK506/APX879 while *h*FKBP12 relies significantly on the 40’s loop and Ile91. The C22 acetohydrazine moiety in APX879 increases the number of significant and differentiating contacts between the human and fungal FKBP12 proteins suggesting that APX879 might be a better starting scaffold than FK506 for modifications to further increase selectivity for the fungal proteins and amplify the difference in the balance between the antifungal and immunosuppressive activities.

## Discussion

Targeting calcineurin is a promising approach towards the development of novel antifungals due to its central role in diverse cellular processes, including antifungal drug resistance and pathogenesis of the major human fungal pathogens(5, 38). Although FK506 efficiently targets calcineurin, currently available formulations are immunosuppressive and, therefore, cannot be used as antifungal therapeutics. In depth, atomistic level understanding of calcineurin inhibition through FK506-FKBP12 binding has the potential to reveal unique features differentiating the human and fungal proteins that could be exploited to enhance fungal selectivity. The studies herein, for the first time, identified distinct differences in selective binding determinants between the fungal and human calcineurin-inhibitor-FKBP12 complexes at atomic resolution that inform future inhibitor design.

Here we report the first crystal structure of *Mc*FKBP12 bound to FK506 revealing structural similarity with other previously crystallized mammalian and fungal FKBP12 proteins (**Fig. 2** and **Sup. Table 1**)(21, 25, 26). Interestingly, *Mc*FKBP12 is not functionally equivalent to *Af*FKBP12 in its inhibitory interaction with *A. fumigatus* calcineurin, primarily due to the critical charge and side-chain size requirements of residue Phe_*Af*_/Tyr_*Mc*_88. MD simulations suggest that *Mc*FKBP12 and *Mc*FKBP12-Y88F interact similarly with FK506, but that the Tyr88Phe mutation alters the hydrogen bonding patterns found in the 50’s and 80’s loop, central for FK506 binding. This mutation does not affect the ability of FKBP12 to bind FK506 or, subsequently, to bind calcineurin but rather alters the formation of a “productive” inhibitory protein-ligand-protein interface (**Fig. 3** and **S1-3**).

Crystal structures of FKBP12 proteins bound to APX879, a less immunosuppressive FK506 analog with an acetohydrazine moiety on FK506-C22, revealed similar interactions as when bound to FK506 (**Fig. 4** and **Table S1**) but, strikingly, also revealed a high degree of flexibility and rearrangement of the acetohydrazine moiety in order to prevent clashes with the FKBP12 residue 88. This mobility might well be restricted by the formation of the ternary complex with calcineurin. In solution, binding of APX879 revealed a 40-fold decrease in affinity for *h*FKBP12 and *Mc*FKBP12 compared to FK506 while *Af*FKBP12 showed a 100-fold reduction in affinity (**Table 1** and **Fig. S4**). Consistent with APX879 being an FK506 analog, the protein responses captured by the NMR chemical shift perturbations were similar when bound to either ligand but notably different when comparing the human and fungal FKBP12 proteins. Interestingly, when fully bound by APX879, both *h*FKBP12 and *Mc*FKBP12 showed several residues in the 40’s, 50’s and 80’s loops experiencing peak doubling not observed when bound to FK506 or in *Af*FKBP12 (**Fig. 5** and **Fig. S5**). This suggests an increased conformational flexibility of regions of *h*FKBP12 and *Mc*FKBP12 central to the formation of the inhibitory ligand-protein interface, not captured solely by the crystal structures.

Areas of the FKBP12 proteins experiencing the greatest conformational difference between the crystal structures and the MD simulations contained a significant number of crystal contacts and are implicated in the formation of the interface for calcineurin binding (**Figs. S6-8**, and **Table S2**). The 80’s loop in all complexes and the 50’s loop in the fungal FKBP12 proteins, showed a high RMSF compared to the crystal structures. Though partially captured in the crystal structures, the MD simulations enabled the exploration of a broader and more detailed scope of conformational flexibility (**Fig. S7**). Altogether, our MD simulation results give credence to the rationale of using conformational ensembles rather than a single X-ray-characterized structure as a search model for rational structure-based ligand design(39-42). A previous report utilizing 14 targets as a test system has benchmarked that using <100 conformers to represent protein conformational flexibility versus a single conformer can increase the accuracy of the protein-ligand interaction predictions by more than 20%(43).

Comparison of the FKBP12-ligand interactions captured by the three structural biology methods used here (*i*.*e*. H-bonds in crystal structures, chemical shift perturbation by NMR, and H-bonds in MD simulations) revealed the interaction between the FKBP12 Ile57 and the ligands atom O2 is maintained in all complexes independent of the methodology (**Table 2** and **Table S3**). Crystal structures and MD simulations show a higher degree of agreement between themselves than with the NMR titrations. This might, in part, be due to the NMR data presented here being limited to the protein backbone chemical shifts (^1^H/^15^N-HSQC), where impact on the side chains is indirectly inferred, and also in part due to the MD simulations initially relying on the crystal structures. These comparisons also allowed the identification of the 50’s loop (residues 50, 52, 55-57, and 60) as being important in all complexes as captured at different levels by these methodologies. Interestingly, in contrast to crystal structures, both the MD simulation and the NMR titration studies determined that the interactions between the fungal FKBP12 proteins and the ligands are exceedingly similar to one another and notably different when compared to the human FKBP12 protein.

**Table 2.**
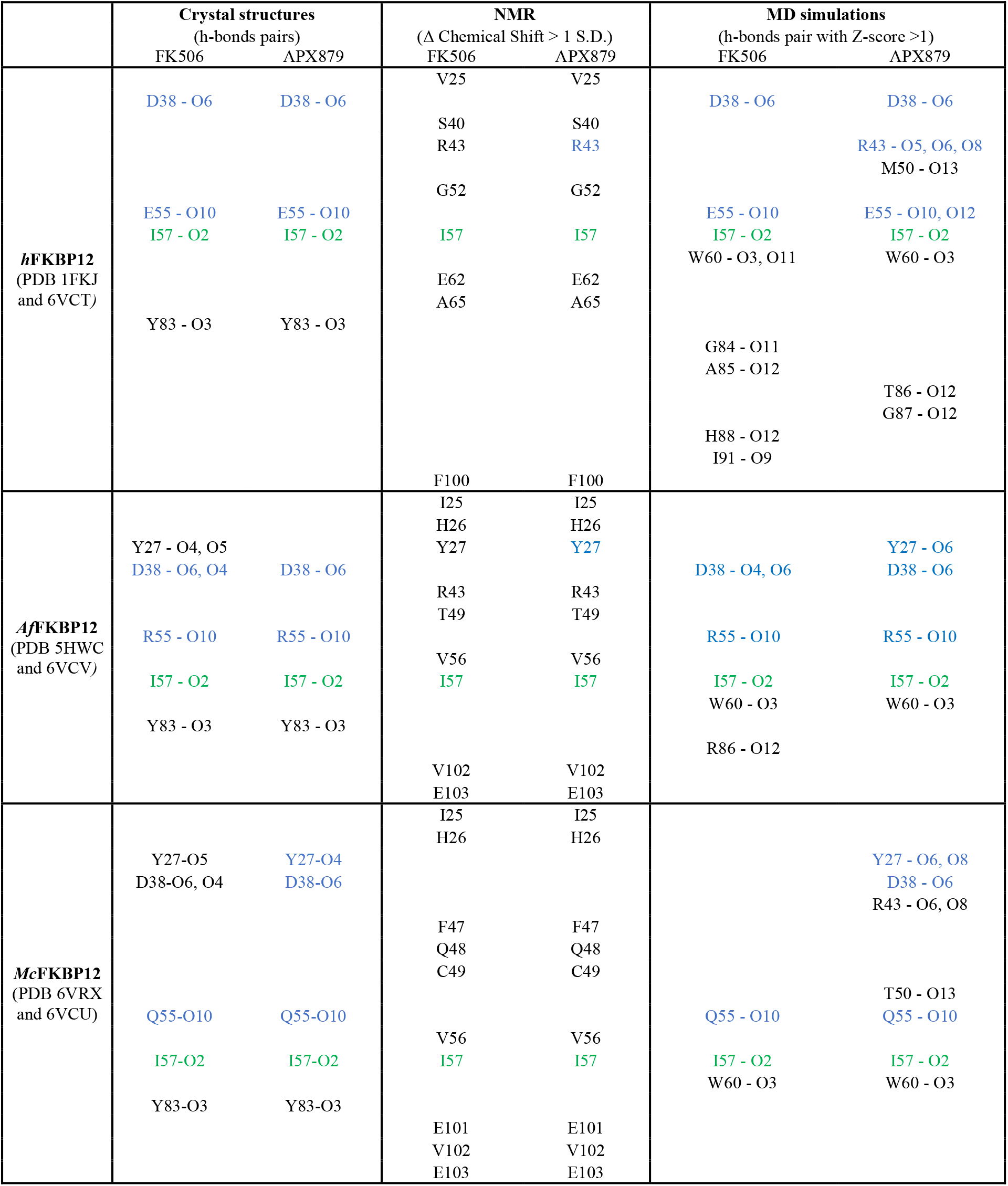
Summary of results from biophysical characterization of FK506 and APX879 complexes. Residues in blue are observed by two methods while in green are observed by the three methods for a same protein-ligand complex. For MD simulation H-bonds Z-scores calculation see **Table S3**.

Additionally, the MD simulations allowed for comparison of the significance of individual FKBP12-ligand interactions, and identification of regions of the ligands where modifications may contribute to enhanced fungal selectivity (**Fig. 6** and **Fig. S9**). Similar free energy scores (ΔG°) for the binding of FK506/APX879 to the FKBP12 proteins were obtained from both the MD simulations and ITC experiments; further justifying their use in rational structure-based design (FK506: MD ≈ -11.3 kcal/mol, ITC ≈ -11.5 kcal/mol; APX879: MD ≈ -10.4 kcal/mol, ITC ≈ -9 kcal/mol). The 80’s loop residues Tyr83 and His_*h*_/Phe_*Af*_/Tyr_*Mc*_88 are key for binding to the fungal FKBP12 proteins while residues Phe49 and Ile91 are more significant for FK506/APX879 binding to *h*FKBP12 (**Fig. 6, Fig. S10**, and **Table S2**). *h*FKBP12 relies more significantly on the pipecolate ring (C2–C5) and C31-O-methylation through atom C45 (cyclohexylidene ring) for FK506 binding while the C35 and C37 methylation sites in the vicinity of the pyranose ring are more significant for binding to the fungal FKBP12 proteins. These FK506 regions could therefore be targeted in an attempt to rationally enhance fungal FKBP12 specific binding; however, these regions are not structurally clustered and involve a small number of atoms.

The APX879 acetohydrazine moiety alters the interactions with the FKBP12 proteins in comparison to FK506 as captured by lower ITC binding affinities, NMR binding assays, and lower MD free-energy values. Z-scores analysis revealed that atoms of the cyclohexylidene (O12, C33, and C42) and pipecolate (C3-C4) rings are significantly implicated for *h*FKBP12 binding, as observed for FK506, but interestingly, the C15-C19 region of APX879 is more significantly involved in binding to *h*FKBP12 (C15-C18, C36, C37, and O8). In contrast, the pyranose (C11, C12, and C35) and part of the cyclohexylidene (C28, C30, and C45) rings show increased importance for interactions with the fungal FKBP12 proteins. In summary, although APX879 makes a greater number of structurally clustered interactions with *h*FKBP12 than with the fungal counterparts, the human and fungal FKBP12 proteins do not rely on the same atoms of APX879 for their interaction. This suggests that APX879 may serve as a more beneficial scaffold for future modifications aimed at enhancing fungal selectivity than FK506.

Our findings corroborate medicinal chemistry studies performed using FK506; identifying the pipecolate ring as important for the immunosuppressive activity(44). Our results are also concordant with a recent report on FK506 analogs modified on the pipecolate and cyclohexylidene (C31) rings showing reduced immunosuppressive activity while, in some cases, maintaining antifungal activity(45). In addition to studies currently underway exploring the effects of selective extensions to the FK506 scaffold centered around C21 and C22 (APX879) (22), the observed reduced immunosuppressive activity around C31 (45), and the selective extensions produced biosynthetically around C9 (26, 27), the data presented here strongly suggests selective extensions or removal of branching atoms centered around C15, C16, C18, C36, and C37. This region is chemically accessible through the creation of the C18-hydroxy analog to which various functional groups can be added (ketones, esters, carbamates, etc). C16 modifications can be achieved through the use of a C18-keto analog.

Further structural, biophysical, and NMR-based dynamics investigations of the calcineurin-APX879-FKBP12 complexes, in combination with the data presented here, will greatly contribute to our understanding of the differential conformational flexibility in the human and fungal complexes, and guide the rational development of fungal-selective and non-immunosuppressive FK506 analogs.

## Methods

### DNA constructs, protein expression and purification

Protein expression for crystallization, isothermal titration calorimetry (ITC) and NMR were performed as previously reported(31). Briefly, *M. circinelloides, A. fumigatus* and human FKBP12 DNA constructs in the pET-15b plasmid (expression with an N-termini hexahistidine tag (His6-tag) cleavable by thrombin) and codon-optimized for *E. coli* expression, were purchased from GenScript (Piscataway, NJ). Expression was performed using *E. coli* BL21(DE3) cells. Cells were propagated at 37°C with agitation to an OD_600_ of 0.6 in modified M9 minimal medium containing 100 μg/mL ampicillin (Sigma Aldrich, St. Louis, MO), 1 g/L NH_4_Cl (or ^15^NH_4_Cl for NMR (Cambridge Isotopes, Tewksbury, MA)) and 55 g/L of sorbitol (*M. circinelloides* only). Protein expression was initiated by the addition of 1 mM Isopropyl β-d-1-thiogalactopyranoside (IPTG) and incubation for 16 h at 25°C with agitation. Cells were harvested by centrifugation at 4°C for 20 minutes at 6000×*g* and the pellet stored at −80°C until purification. The cell pellets were resuspended in 30 mL of 50 mM sodium phosphate, 500 mM NaCl, pH 8.0 buffer. Lysis was performed by three cycles of 30 seconds of sonication at a power of 12 watts with a 2-minute rest interval on ice or by French press followed by the addition of 1 mM of phenylmethylsulfonyl fluoride (PMSF). The lysate was clarified by centrifugation (4°C, 15 min at 20,000×*g*) and filtration using a 0.22 µm PES syringe filter. The chromatography was undertaken at 4°C using an ÄKTA FPLC (GE Healthcare). The clarified supernatant was loaded onto a 5 mL prepacked Ni-nitrilotriacetic acid (NTA) column. Protein was eluted using a 0 to 1 M stepwise gradient of imidazole. Fractions containing FKBP12 proteins were identified by SDS-PAGE gel electrophoresis and Coomassie Blue staining. Combined fractions were dialyzed into 50 mM sodium phosphate, 500 mM NaCl, pH 8.0 buffer to remove the imidazole (2 cycles in 1 L for 2 h followed by 1 cycle in 2 L for 16-18 h). The His6-tag was cleaved for 16 h at 4°C using 1U of thrombin (GE Healthcare) per 100 μg of total protein. The cleaved proteins were then run over the 5 mL prepacked Ni–NTA column to remove the cleaved His-tag and any uncleaved protein. For crystallography, in presence of the ligand, APX879 or FK506 were added in a 1 to 1.5 molar ratio of FKBP12:ligand using a solution stock of the ligand at 10 mg/mL in 100% DMSO. The total volume of DMSO added was limited to less than 5%. In all cases, protein solutions were concentrated to a volume of ∼2 mL using a 3000 MWCO Amicon concentrator followed by size-exclusion chromatography using a Sephacyrl S100HR XK26/60 FPLC column. Fractions containing protein were identified by SDS-PAGE gel electrophoresis and Coomassie Blue staining. Typical yields were 40 mg/L of > 98% pure protein.

### Crystallization and determination of the structures of M. circinelloides bound to FK506 and M. circinelloides, A. fumigatus and H. sapiens FKBP12 bound to APX879

After size exclusion purification, the proteins were concentrated to 10 mg/mL using a 3000 MWCO Amicon concentrator. For *Mc*FKBP12/FK506 and *h*FKBP12/APX879, crystals were grown at 22°C with a hanging drop vapor diffusion setting using a 1 to 1 ratio of the protein and reservoir solutions. *Mc*FKBP12/FK506 crystals were grown in 2100 mM DL malic acid pH 7.0. Needle shaped crystals of *h*FKBP12/APX879 were grown with 2.5 M ammonium sulfate, 0.1 M sodium acetate trihydrate pH 4.6 as the reservoir solution. For *Mc*FKBP12/APX879, the same hanging drop vapor diffusion setting was used with a protein to the reservoir solution ratio of 0.33-0.66. Crystals were grown with 1600 mM sodium citrate tribasic as the reservoir solution. For *Af*FKBP12/APX879, initial crystals were grown at 22°C using a hanging drop vapor diffusion setting and 10 mM MES pH 6.0, 200 mM zinc acetate and 15% reagent alcohol as the reservoir solution. Protein drops were prepared using a 1 to 1 ratio of the protein and reservoir solution. A single crystal was then harvested to prepare a seed stock using the Hampton Research (Aliso Viejo, CA) Seed Bead kit and the Classical method. Crystals were grown in 5 mM MES pH 6.0, 200 mM zinc acetate and 15% reagent alcohol using a 1 to 1 ratio of the protein and reservoir solution and streaking of the drop using the seed stock. All crystals were cryopreserved directly from the drop. Diffraction data were collected at the Advanced Photon Source using sector 22 BM and ID beamlines. The collected diffraction images were indexed, integrated, and scaled using HKL2000(46). Initial phases were calculated by molecular replacement using Phenix.PHASER(47, 48) and the PDBs 5HUA (*Mc*FKBP12/FK506 or APX879), 2PPN (*h*FKBP12/APX879) and 5HWB (*Af*FKBP12/APX879) as search models(21, 26). Iterative rounds of manual model building using Coot(49) and automatic refinement in PHENIX(47) were performed. Data collection and refinement statistics are summarized in **Table S1**. The refined structures have been deposited to the Protein Data Bank (https://www.rcsb.org) under the accession codes 6VRX, 6VCT, 6VCU and 6VCV.

### Active-site Volume Estimation

The binding pocket cavity volume was estimated using 3V: Voss Volume Voxelator(28). The estimation was made with a small sphere of 1.5 Å radius and a large sphere of 7 Å radius.

### Isothermal titration calorimetry (ITC) experiments

After the size exclusion purification, the proteins were concentrated to ∼2 mg/mL using a 3000 MWCO Amicon concentrator. Proteins were exhaustively dialyzed at 4°C in the ITC buffer (50 mM sodium phosphate, 50 mM sodium chloride pH 7.0). Experiments were performed using a VP-ITC instrument (Microcal Inc. Northampton, MA) at 25°C. All solutions were degassed under vacuum at 25°C for 15 minutes immediately before use. For FK506, because of its high affinity, it was used in the sample cell at a concentration of 2 µM with constant stirring at 307 rpm. The protein was loaded into the syringe at a concentration of 25 µM and titrated in by a first injection of 2 µL followed by 22 injections of 6 µL. Following each injection, the cell was equilibrated for 3 minutes. For APX879, because of its low solubility restricting the usable concentration and its lower affinity, it was used in the syringe at a concentration of 150 µM. The proteins were loaded in the cell at a concentration of 10 µM with constant stirring at 307 rpm (25°C). APX879 was titrated in by a first injection of 2 µL followed by 29 injections of 8 µL and equilibrated for 3 minutes. The enthalpy of binding (ΔH, kcal/mol) was determined by integration of the injection peaks minus the controls for the heat of dilution (equivalent experiments without the ligand and without the protein). The MicroCal Origin software (OriginLab Corp., Northampton, MA) was used for a variety of binding models.

### Construction of the A. fumigatus strain expressing McFKBP12 and in vitro susceptibility assays

The *Mcfkbp12* expression construct was codon optimized for expression in *A. fumigatus* and synthesized by GenScript and cloned into the pUC57 vector at KpnI-BamHI sites to generate the pUC57-McFkbp12 vector. The codon optimized *Mcfkbp12* gene along with the 800 bp *Affkbp12* promoter from the pUC57-McFkbp12 vector was then cloned into the pUCGH(50) vector at KpnI-BamHI sites to generate the pUCGH-McFkbp12promo-McFkbp12 vector. To facilitate homologous recombination a 1000 bp *Affkbp12* terminator was then cloned at the SbfI-HindIII on pUCGH-McFkbp12promo-McFkbp12 vector to generate the final pUCGH-McFkbp12promo-McFkbp12-McFkbp12term vector. The pUCGH-McFkbp12promo-McFkbp12-McFkbp12term vector was linearized by digestion with KpnI and the 7148 bp linearized fragment was transformed into the *A. fumigatus akuB*^KU80^ strain. Transformants were selected with hygromycin B (150 µg/ml). The *Mcfkbp12-Y88F* mutant construct was generated by site-directed mutagenesis PCR using the primers pUCGH-2033-F (GCGTTGGCCGATTCATTA) and McFkbp12-Y88F-R (AAGTCCAGGGAAGCCGCGCTC) to obtain a 1290 bp PCR fragment, and McFkbp12-Y88H-F (GAGCGCGGCTTCCCTGGACTT) and Hyg-R (GCCCATGAACTGGCTCTTAA) to obtain a 1267 bp PCR fragment. Fusion PCR was then performed with pUCGH-2033-F and Hyg-R using a 1:1 mixture of the two PCR fragments (1290 bp and 1267 bp) to obtain the final 2536 bp PCR fragment harboring the Y88F mutation. This 2536 bp fragment was digested with NotI-KpnI and cloned into pUCGH at NotI-KpnI sites to obtain the pUCGH-McFkbp12promo-McFkbp12-Y88F vector. In the next step the *Affkbp12* terminator was inserted at SbfI-HindIII as described for the pUCGH-McFkbp12promo-McFkbp12 vector followed by linearization and transformation into the *A. fumigatus akuB*^KU80^ strain. All the constructs were sequenced to confirm accuracy before using for transformations. The transformants were verified for homologous integration by PCR, and accuracy of the *Mcfkbp12* and *Mcfkbp12-Y88F* sequences was verified by sequencing and visualized by fluorescent microscopy. *E. coli* DH5α competent cells were used for subcloning experiments. *A. fumigatus* wild-type strain *akuB*^*KU80*^ was used for growth and recombinant strain generation experiments. *A. fumigatus* wild-type or recombinant strains were cultured on glucose minimal media (GMM) or RPMI liquid media at 37°C for 24 or 48 h time periods. In certain experiments, GMM agar or RPMI liquid media were supplemented with FK506 (0.01-4 µg/mL) or APX879 (0.01-4 µg/mL). All growth experiments were repeated as technical triplicates, each also as biological triplicates.

### Fluorescence microscopy

Conidia (10^4^) from the recombinant strains of *A. fumigatus* were inoculated into 5 ml GMM medium and poured over a sterile coverslip (22×60 mm; No.1) placed in a sterile dish (60×15 mm). Cultures grown for 18-20 h at 37°C were observed by fluorescence microscopy using an Axioskop 2 plus microscope (Zeiss) equipped with AxioVision 4.6 imaging software. FK506 (100 ng/mL) was added to the cultures in order to visualize the *in vivo* binding of FKBP12 to the calcineurin complex at the septum.

### Molecular dynamic simulations

MD simulations were performed to provide a better representation of the protein’s conformational flexibility and to more accurately characterize the proteins’ solution structure bound to APX879 and FK506(19, 21). Crystal structures were used as the starting conformations: *Mc*FKBP12 bound to FK506 (PDB 6VRX) and APX879 (PDB 6VCT), *h*FKBP12 bound to FK506 (PDB 1FKF) and APX879 (PDB 6VCU), and *Af*FKBP12 bound to FK506 (PDB 5HWC – P90G mutant) and APX879 (PDB 6VCV). For the *Mc*FKBP12(Y88F) mutant bound to FK506, the wild-type crystal structure was used and mutated *in silico* accordingly. FK506 and APX879 small molecule parameter and topology files were downloaded and created utilizing Automated Topology Builder (ATB) and repository(51, 52). All molecular dynamic (MD) simulations were performed with the GROMACS 5.0.1 software package utilizing 6 CPU cores and one NVIDIA Tesla K80 GPU(53). The single starting conformations used for all of the MD simulations were resulting X-ray characterized crystal structures described here or otherwise noted above. MD simulations were performed with the GROMOS54a7 force field and the flexible simple point-charge water model. The initial structures were immersed in a periodic water box with a dodecahedron shape that extended 1 nm beyond the protein in any dimension and neutralized with counterions. Energy minimization was accomplished through use of the steepest descent algorithm with a final maximum force below 100 kJ/mol/min (0.01 nm step size, cutoff of 1.2 nm for neighbor list, Coulomb interactions, and Van der Waals interactions). After energy minimization, the system was subjected to equilibration at 300K and normal pressure for 1 ns. All bonds were constrained with the LINCS algorithm (cutoff of 1.2 nm neighbor list, Coulomb interactions, and Van der Waals interactions). After temperature stabilization, pressure stabilization was obtained by utilizing the v-rescale thermostat to hold the temperature at 300K and the Berendsen barostat was used to bring the system to 1 bar pressure. Production MD calculations (500 ns) were performed under the same conditions, except that the position restraints were removed, and the simulation was run for 500 ns (cutoff of 1.1, 0.9, and 0.9 nm for neighbor list, Coulomb interactions, and Van der Waals interactions). These MD simulations were repeated 6 times, with the exception of the *Mc*FKBP12(Y88F)-FK506 which was performed once. Cα-RMSD, SASA, Rg, and COM all confirmed stability and accuracy of the MD simulations by stabilizing after a 100 ns equilibration period (in most cases) allowing for the analysis of the last 400 ns of the simulations. Only three of the 36 calculations (6 complex repeated 6 times) were rejected from the analysis due to unstable Cα-RMSD, Rg, and/or COM (**Fig. S6**).

GROMACS built-in and homemade scripts were used to analyze the MD simulation results and averaged over the 6 simulations. All images were produced using PyMOL(53). Atom indices for FK506 and APX879 are provided in **Tables S4** and **S5**.

### NMR

[^15^N]-labeled samples were concentrated to 0.4–0.7 mM and buffer exchanged into 20 mM sodium phosphate, 100 mM NaCl, 0.02% NaN_3_ and 5% D_2_O, pH 6.0 using a 3000 MWCO Amicon concentrator. All NMR experiments were performed at 25°C, as calibrated with a standard methanol sample. Previously reported NMR backbone resonance assignments were used (BMRB Codes 27732, 27733, 27734, 27737, 27738, 27739)(31). All NMR experiments were performed on a Bruker Avance III spectrometer at 16.4T (700 MHz) equipped with a 4 nucleus QXI probe and pulsed-field Z-gradient. NMR data were processed using NMRPipe(54) and analyzed using Sparky(55, 56) and NMRViewJ version 8.0(57). Chemical shifts were referenced to an external 2,2-dimethyl-2-silapentane-5-sulfonate (DSS) sample.

## Acknowledgement

This work was supported by grants from the NIH/NIAID (R01 AI112595-04; P01 AI104533-05). Compound APX879 was supplied by Amplyx Pharmaceuticals, Inc. Use of the Advanced Photon Source was supported by the U. S. Department of Energy, Office of Science, Office of Basic Energy Sciences, under Contract No. W-31-109-Eng-38. Data were collected at Southeast Regional Collaborative Access Team (SER-CAT) 22-ID and 22-BM beamline at the Advanced Photon Source, Argonne National Laboratory. SER-CAT is supported by its member institutions (http://www.ser-cat.org/members.html), and equipment grants (S10_RR25528 and S10_RR028976) from the National Institutes of Health. The authors thank Duke’s Research Computing staff for the use of Duke Computing Cluster and its support. The authors also thank Nathan Nicely and Priyamvada Acharya for the use of the equipment for protein crystallization and Michael Hoy, Zanetta Chang, Soo Chan Lee, and Anna Averette for support and discussions. Use of the Duke NMR Spectroscopy Center instrumentation is gratefully acknowledged. S.M.G. was the recipient of a Natural Sciences and Engineering Research Council of Canada (NSERC) postdoctoral fellowship.

## Author Contribution

S.M.G. designed, and performed protein expression and X-ray crystallography, and isothermal titration calorimetry (ITC) assays; analyzed and interpreted data and wrote the manuscript. B.G.B. designed, performed, and analyzed the MD simulations, and contributed to writing. P.R.J. supervised, designed, and performed genetic, biochemical, and microscopy experiments and contributed to writing. D.C.C. performed antifungal susceptibility experiments. R.A.V. and S.M.G. designed, performed, and interpreted NMR experiments. J.H., W.J.S., and L.D.S designed and supervised the overall study. All authors were involved in editing the manuscript.

## Competing interests

The authors declare no competing interests.

## Notes

### Competing Interest Statement

The authors have declared no competing interest.

## References

1. F. Bongomin, S. Gago, R. Oladele, D. Denning, Global and multi-national prevalence of fungal diseases—Estimate precision. Journal of Fungi 3, 57 (2017).

2. A. H. Kachroo et al., Systematic humanization of yeast genes reveals conserved functions and genetic modularity. Science 348, 921–925 (2015).

3. P. T. Marcyk et al., Fungal-Selective Resorcylate Aminopyrazole Hsp90 Inhibitors: Optimization of Whole-Cell Anticryptococcal Activity and Insights into the Structural Origins of Cryptococcal Selectivity. Journal of Medicinal Chemistry 64, 1139–1169 (2021).

4. L. Whitesell et al., Structural basis for species-selective targeting of Hsp90 in a pathogenic fungus. Nature Communications 10 (2019).

5. P. R. Juvvadi, S. C. Lee, J. Heitman, W. J. Steinbach, Calcineurin in fungal virulence and drug resistance: Prospects for harnessing targeted inhibition of calcineurin for an antifungal therapeutic approach. Virulence 8, 186–197 (2017).

6. H. S. Park, S. C. Lee, M. E. Cardenas, J. Heitman, Calcium-calmodulin-calcineurin signaling: A globally conserved virulence cascade in eukaryotic microbial pathogens. Cell Host Microbe 26, 453–462 (2019).

7. C. B. Klee, T. H. Crouch, M. H. Krinks, Calcineurin: a calcium- and calmodulin-binding protein of the nervous system. Proceedings of the National Academy of Sciences of the United States of America 76, 6270–6273 (1979).

8. C. S. Hemenway, J. Heitman, Calcineurin. Cell Biochemistry and Biophysics 30, 115–151 (1999).

9. N. A. Clipstone, G. R. Crabtree, Identification of calcineurin as a key signalling enzyme in T-lymphocyte activation. Nature 357, 695–697 (1992).

10. S. J. O’Keefe, J. i. Tamura, R. L. Kincaid, M. J. Tocci, E.A. O’Neill, FK-506- and CsA-sensitive activation of the interleukin-2 promoter by calcineurin. Nature 357, 692–694 (1992).

11. E. W. L. Chow et al., Elucidation of the calcineurin-Crz1 stress response transcriptional network in the human fungal pathogen Cryptococcus neoformans. PLoS Genetics 13, e1006667 (2017).

12. R. A. Cramer, Jr. et al., Calcineurin target CrzA regulates conidial germination, hyphal growth, and pathogenesis of Aspergillus fumigatus. Eukaryot Cell 7, 1085–1097 (2008).

13. A. H. Andreotti, Native state proline isomerization: An intrinsic molecular switch. Biochemistry 42, 9515–9524 (2003).

14. G. Fischer, T. Aumüller, “Regulation of peptide bond cis/trans isomerization by enzyme catalysis and its implication in physiological processes” in Reviews of Physiology, Biochemistry and Pharmacology. (2003), 10.1007/s10254-003-0011-3 chap. Chapter 3, pp. 105–150.

15. J. Fanghanel, G. Fischer, Insights into the catalytic mechanism of peptidyl prolyl cis/trans isomerases. Frontiers in Bioscience 9, 3453–3478 (2004).

16. B. Aghdasi et al., FKBP12, the 12-kDa FK506-binding protein, is a physiologic regulator of the cell cycle. Proceedings of the National Academy of Sciences 98, 2425–2430 (2001).

17. R. K. Harrison, R. L. Stein, Substrate specificities of the peptidyl prolyl cis-trans isomerase activities of cyclophilin and FK-506 binding protein: evidence for the existence of a family of distinct enzymes. Biochemistry 29, 3813–3816 (2002).

18. S. Schreiber, Chemistry and biology of the immunophilins and their immunosuppressive ligands. Science 251, 283–287 (1991).

19. G. Van Duyne, R. Standaert, P. Karplus, S. Schreiber, J. Clardy, Atomic structure of FKBP-FK506, an immunophilin-immunosuppressant complex. Science 252, 839–842 (1991).

20. S. Michnick, M. Rosen, T. Wandless, M. Karplus, S. Schreiber, Solution structure of FKBP, a rotamase enzyme and receptor for FK506 and rapamycin. Science 252, 836–839 (1991).

21. N. K. Tonthat et al., Structures of Pathogenic Fungal FKBP12s Reveal Possible Self-Catalysis Function. mBio 7, e00492–00416 (2016).

22. P. R. Juvvadi et al., Harnessing calcineurin-FK506-FKBP12 crystal structures from invasive fungal pathogens to develop antifungal agents. Nature Communications 10, 4275 (2019).

23. U. Binder, E. Maurer, C. Lass-Flörl, Mucormycosis – from the pathogens to the disease. Clinical Microbiology and Infection 20, 60–66 (2014).

24. J. R. Kohler, A. Casadevall, J. Perfect, The spectrum of fungi that infects humans. Cold Spring Harbor Perspectives in Medicine 5, a019273–a019273 (2014).

25. K. P. Wilson et al., Comparative X-ray structures of the major binding protein for the immunosuppressant FK506 (tacrolimus) in unliganded form and in complex with FK506 and rapamycin. Acta Crystallographica Section D Biological Crystallography 51, 511–521 (1995).

26. S. Szep, S. Park, E. T. Boder, G. D. Van Duyne, J. G. Saven, Structural coupling between FKBP12 and buried water. Proteins: Structure, Function, and Bioinformatics 74, 603–611 (2009).

27. K. Falloon et al., Characterization of the FKBP12-encoding genes in Aspergillus fumigatus. PLoS One 10, e0137869 (2015).

28. N. R. Voss, M. Gerstein, 3V: cavity, channel and cleft volume calculator and extractor. Nucleic Acids Res 38, W555–562 (2010).

29. Y. Shen, A. Bax, Prediction of Xaa-Pro peptide bond conformation from sequence and chemical shifts. Journal of Biomolecular NMR 46, 199–204 (2009).

30. Y. C. Lee et al., NMR conformational analysis of cis and trans proline isomers in the neutrophil chemoattractant, N-acetyl-proline-glycine-proline. Biopolymers 58, 548–561 (2001).

31. S. M. C. Gobeil, B. G. Bobay, L. D. Spicer, R. A. Venters, 15N, 13C and 1H resonance assignments of FKBP12 proteins from the pathogenic fungi Mucor circinelloides and Aspergillus fumigatus. Biomolecular NMR Assignments 13, 207–212 (2019).

32. J. P. Griffith et al., X-ray structure of calcineurin inhibited by the immunophilin-immunosuppressant FKBP12-FK506 complex. Cell 82, 507–522 (1995).

33. B. E. Bierer et al., Two distinct signal transmission pathways in T lymphocytes are inhibited by complexes formed between an immunophilin and either FK506 or rapamycin. Proceedings of the National Academy of Sciences 87, 9231–9235 (1990).

34. P. R. Connelly et al., Enthalpy of hydrogen bond formation in a protein-ligand binding reaction. Proceedings of the National Academy of Sciences 91, 1964–1968 (1994).

35. P. R. Connelly, J. A. Thomson, Heat capacity changes and hydrophobic interactions in the binding of FK506 and rapamycin to the FK506 binding protein. Proceedings of the National Academy of Sciences 89, 4781–4785 (1992).

36. G. Solomentsev, C. Diehl, M. Akke, Conformational entropy of FK506 binding to FKBP12 determined by nuclear magnetic resonance relaxation and molecular dynamics simulations. Biochemistry 57, 1451–1461 (2018).

37. M. A. Wear, A. Patterson, M. D. Walkinshaw, A kinetically trapped intermediate of FK506 binding protein forms in vitro: Chaperone machinery dominates protein folding in vivo. Protein Expression and Purification 51, 80–95 (2007).

38. W. J. Steinbach, J. L. Reedy, R. A. Cramer, Jr., J. R. Perfect, J. Heitman, Harnessing calcineurin as a novel anti-infective agent against invasive fungal infections. Nat Rev Microbiol 5, 418–430 (2007).

39. N. Floquet et al., Normal mode analysis as a prerequisite for drug design: Application to matrix metalloproteinases inhibitors. FEBS Letters 580, 5130–5136 (2006).

40. D. A. Antunes, D. Devaurs, L. E. Kavraki, Understanding the challenges of protein flexibility in drug design. Expert Opinion on Drug Discovery 10, 1301–1313 (2015).

41. S. E. Allen, N. V. Dokholyan, A. A. Bowers, Dynamic docking of conformationally constrained macrocycles: methods and applications. ACS Chemical Biology 11, 10–24 (2015).

42. C. Pallara, M. Rueda, R. Abagyan, J. Fernández-Recio, Conformational heterogeneity of unbound proteins enhances recognition in protein–protein encounters. Journal of Chemical Theory and Computation 12, 3236–3249 (2016).

43. M. Rueda, G. Bottegoni, R. Abagyan, Consistent improvement of cross-docking results using binding site ensembles generated with elastic network normal modes. Journal of Chemical Information and Modeling 49, 716–725 (2009).

44. M. T. Goulet, K. M. Rupprecht, P. J. Sinclair, M. J. Wyvratt, W. H. Parsons, The medicinal chemistry of FK-506. Perspectives in Drug Discovery and Design 2, 145–162 (1994).

45. Y. Lee et al., In vitro and in vivo assessment of FK506 analogs as novel antifungal drug candidates. Antimicrobial Agents and Chemotherapy 62, e01627–01618 (2018).

46. Z. Otwinowski, W. Minor, “Processing of X-ray diffraction data collected in oscillation mode” in Macromolecular Crystallography Part A. (1997), 10.1016/s0076-6879(97)76066-x, pp. 307–326.

47. P. D. Adams et al., PHENIX: a comprehensive Python-based system for macromolecular structure solution. Acta Crystallographica Section D Biological Crystallography 66, 213–221 (2010).

48. A. J. McCoy et al., Phaser crystallographic software. Journal of Applied Crystallography 40, 658–674 (2007).

49. P. Emsley, B. Lohkamp, W. G. Scott, K. Cowtan, Features and development of Coot. Acta Crystallographica Section D Biological Crystallography 66, 486–501 (2010).

50. P. R. Juvvadi et al., Calcineurin localizes to the hyphal septum in Aspergillus fumigatus: implications for septum formation and conidiophore development. Eukaryot Cell 7, 1606–1610 (2008).

51. A. K. Malde et al., An automated force field topology builder (ATB) and repository: Version 1.0. J Chem Theory Comput 7, 4026–4037 (2011).

52. M. Stroet et al., Automated topology builder version 3.0: Prediction of solvation free enthalpies in water and hexane. J Chem Theory Comput 14, 5834–5845 (2018).

53. M. J. Abraham et al., GROMACS: High performance molecular simulations through multi-level parallelism from laptops to supercomputers. SoftwareX 1-2, 19–25 (2015).

54. F. Delaglio et al., NMRPipe: a multidimensional spectral processing system based on UNIX pipes. J Biomol NMR 6, 277–293 (1995).

55. T. D. Goddard, D. G. Kneller, SPARKY 3. University of California, San Francisco, CA (2008).

56. W. Lee, M. Tonelli, J. L. Markley, NMRFAM-SPARKY: enhanced software for biomolecular NMR spectroscopy. Bioinformatics 31, 1325–1327 (2015).

57. B. A. Johnson, R. A. Blevins, NMR View: A computer program for the visualization and analysis of NMR data. J Biomol NMR 4, 603–614 (1994).

