## Supporting Information for "Leveraging Fungal Calcineurin-Inhibitor Structures, Biophysics and Dynamics to Design Selective and Non-Immunosuppressive FK506 Analogs"

**Table S1. Data collection and refinement statistics for the FK506-*Mc*FKBP12 and human, *Af* and *Mc* FKBP12 proteins bound APX879 crystal structures.**

| | *Mc*FKBP12-FK506 PDB 6VRX | *h*FKBP12-APX879 PDB 6VCU | *Af*FKBP12- APX879 PDB 6VCV | *Mc*FKBP12- APX879 PDB 6VCT |
|---|---|---|---|---|
| Data collection | | | | |
| Space group | $P3_221$ | $P3_2$ | $P1$ | $C222_1$ |
| Unit-cell dimensions | | | | |
| a, b, c (Å) | 104.9, 104.9, 111.6 | 53.6, 53.6, 126.9 | 35.6, 39.6, 40.8 | 58.5, 75.5, 46.5 |
| α, β, ϒ (º) | 90.0, 90.0, 120.0 | 90.0, 90.0, 120.0 | 76.8, 89.9, 85.7 | 90.0, 90.0, 90.0 |
| Resolution (Å) | 37.20-2.54 | 31.26-1.69 | 27.12-1.60 | 32.81-1.94 |
| CC(1/2) | 99.6 (80.6) | 98.5 (94.0) | 98.6 (94.0) | 99.6 (99.4) |
| CC* | 99.9 (94.5) | 99.6 (98.4) | 99.7 (98.4) | 99.9 (99.9) |
| Rpim | 4.3 | 5.3 | 7.3 | 2.1 |
| Overall R-merge (%) | 9.0 | 12.2 | 9.8 | 7.6 |
| I/σ(I) | 27.1 (2.05) | 30.5 (5.08) | 33.9 (14.74) | 56.9 (13.73) |
| Completeness (%) | 99.6 (96.9) | 99.9 (98.8) | 96.2 (96.0) | 99.7 (99.0) |
| Redundancy | 9.3 (10.2) | 6.5 (6.2) | 2.3 (2.3) | 14.5 (14.6) |
| Refinement | | | | |
| No. reflection/unique | 221,719/23,722 | 293,678/45,479 | 63,353/27,429 | 114,812/7,938 |
| R-work/R-free (%) | 19.7/24.4 | 15.6/19.6 | 17.6/21.4 | 15.4/20.6 |
| Molprobity | | | | |
| Ramachandran favored | 94.06 | 97.16 | 98.62 | 97.14 |
| Ramachandran outlier | 0.24 | 0.00 | 0.00 | 0.00 |
| Rotamer outliers | 3.83 | 0.00 | 0.00 | 0.00 |
| Clash score | 5 | 3 | 2 | 1 |
| No. of water | 30 | 591 | 332 | 120 |
| RMSD | | | | |
| Bond lengths (Å) | 0.007 | 0.007 | 0.007 | 0.006 |
| Bond angles (°) | 1.15 | 1.19 | 1.35 | 1.07 |

**Table S2. Crystal contacts between symmetry related molecules in crystal structures (Excel document).**

**Supplementary Table 2. Crystal contacts for each of the FKBP12 crystal structures under study.**

| Residue number | Residue | 1 | 2 | 3 | 4 | 5 | 6 | 7 | 8 | 9 | 10 | 11 | 12 | 13 | 14 | 15 | 16 | 17 | 18 | 19 | 20 | 21 | 22 | 23 | 24 | 25 | 26 | 27 | 28 | 29 | 30 | 31 | 32 | 33 | 34 | 35 | 36 | 37 | 38 | 39 | 40 |
|---|---|---|---|---|---|---|---|---|---|---|---|---|---|---|---|---|---|---|---|---|---|---|---|---|---|---|---|---|---|---|---|---|---|---|---|---|---|---|---|---|---|
| HsFKB12/FK506 (PDB_1FKJ) | - | | | | | | | | | | | | | | | | | | | | | | | | | | | | | | | | | | | | | | | | |
| AfFKB12/FK506 (PDB_5HWC) | - | | | | | | | | | | | | | | | | | | | | | | | | | | | | | | | | | | | | | | | | |
| McFKB12/FK506 | Chain A | | | | | | | | | | | | | | | | | | | | | | | | | | | | | | | | | | | | | | | | |
| McFKB12/FK506 | Chain B | | | | | | | | | | | | | | | | | | | | | | | | | | | | | | | | | | | | | | | | |
| McFKB12/FK506 | Chain C | | | | | | | | | | | | | | | | | | | | | | | | | | | | | | | | | | | | | | | | |
| McFKB12/FK506 | Chain D | | | | | | | | | | | | | | | | | | | | | | | | | | | | | | | | | | | | | | | | |
| HsFKB12/APX | Chain A | | | | | | | | | | | | | | | | | | | | | | | | | | | | | | | | | | | | | | | | |
| HsFKB12/APX | Chain B | | | | | | | | | | | | | | | | | | | | | | | | | | | | | | | | | | | | | | | | |
| HsFKB12/APX | Chain C | | | | | | | | | | | | | | | | | | | | | | | | | | | | | | | | | | | | | | | | |
| HsFKB12/APX | Chain D | | | | | | | | | | | | | | | | | | | | | | | | | | | | | | | | | | | | | | | | |
| AfFKB12/APX879 | Chain A | | | | | | | | | | | | | | | | | | | | | | | | | | | | | | | | | | | | | | | | |
| AfFKB12/APX879 | Chain B | | | | | | | | | | | | | | | | | | | | | | | | | | | | | | | | | | | | | | | | |
| McFKB12/APX879 | - | | | | | | | | | | | | | | | | | | | | | | | | | | | | | | | | | | | | | | | | |

| Residue number | | 41 | 42 | 43 | 44 | 45 | 46 | 47 | 48 | 49 | 50 | 51 | 52 | 53 | 54 | 55 | 56 | 57 | 58 | 59 | 60 | 61 | 62 | 63 | 64 | 65 | 66 | 67 | 68 | 69 | 70 | 71 | 72 | 73 | 74 | 75 | 76 | 77 | 78 | 79 | 80 |
|---|---|---|---|---|---|---|---|---|---|---|---|---|---|---|---|---|---|---|---|---|---|---|---|---|---|---|---|---|---|---|---|---|---|---|---|---|---|---|---|---|---|
| HsFKB12/FK506 (PDB_1FKJ) | - | | | | | | | | | | | | | | | | | | | | | | | | | | | | | | | | | | | | | | | | |
| AfFKB12/FK506 (PDB_5HWC) | - | | | | | | | | | | | | | | | | | | | | | | | | | | | | | | | | | | | | | | | | |
| McFKB12/FK506 | Chain A | | | | | | | | | | | | | | | | | | | | | | | | | | | | | | | | | | | | | | | | |
| McFKB12/FK506 | Chain B | | | | | | | | | | | | | | | | | | | | | | | | | | | | | | | | | | | | | | | | |
| McFKB12/FK506 | Chain C | | | | | | | | | | | | | | | | | | | | | | | | | | | | | | | | | | | | | | | | |
| McFKB12/FK506 | Chain D | | | | | | | | | | | | | | | | | | | | | | | | | | | | | | | | | | | | | | | | |
| HsFKB12/APX | Chain A | | | | | | | | | | | | | | | | | | | | | | | | | | | | | | | | | | | | | | | | |
| HsFKB12/APX | Chain B | | | | | | | | | | | | | | | | | | | | | | | | | | | | | | | | | | | | | | | | |
| HsFKB12/APX | Chain C | | | | | | | | | | | | | | | | | | | | | | | | | | | | | | | | | | | | | | | | |
| HsFKB12/APX | Chain D | | | | | | | | | | | | | | | | | | | | | | | | | | | | | | | | | | | | | | | | |
| AfFKB12/APX879 | Chain A | | | | | | | | | | | | | | | | | | | | | | | | | | | | | | | | | | | | | | | | |
| AfFKB12/APX879 | Chain B | | | | | | | | | | | | | | | | | | | | | | | | | | | | | | | | | | | | | | | | |
| McFKB12/APX879 | - | | | | | | | | | | | | | | | | | | | | | | | | | | | | | | | | | | | | | | | | |

| Residue number | | 81 | 82 | 83 | 84 | 85 | 86 | 87 | 88 | 89 | 90 | 91 | 92 | 93 | 94 | 95 | 96 | 97 | 98 | 99 | 100 | 101 | 102 | 103 | 104 | 105 | 106 | 107 | 108 | 109 | 110 | 111 | 112 |
|---|---|---|---|---|---|---|---|---|---|---|---|---|---|---|---|---|---|---|---|---|---|---|---|---|---|---|---|---|---|---|---|---|---|
| HsFKB12/FK506 (PDB_1FKJ) | - | | | | | | | | | | | | | | | | | | | | | | | | | | | | | | | | |
| AfFKB12/FK506 (PDB_5HWC) | - | | | | | | | | | | | | | | | | | | | | | | | | | | | | | | | | |
| McFKB12/FK506 | Chain A | | | | | | | | | | | | | | | | | | | | | | | | | | | | | | | | |
| McFKB12/FK506 | Chain B | | | | | | | | | | | | | | | | | | | | | | | | | | | | | | | | |
| McFKB12/FK506 | Chain C | | | | | | | | | | | | | | | | | | | | | | | | | | | | | | | | |
| McFKB12/FK506 | Chain D | | | | | | | | | | | | | | | | | | | | | | | | | | | | | | | | |
| HsFKB12/APX | Chain A | | | | | | | | | | | | | | | | | | | | | | | | | | | | | | | | |
| HsFKB12/APX | Chain B | | | | | | | | | | | | | | | | | | | | | | | | | | | | | | | | |
| HsFKB12/APX | Chain C | | | | | | | | | | | | | | | | | | | | | | | | | | | | | | | | |
| HsFKB12/APX | Chain D | | | | | | | | | | | | | | | | | | | | | | | | | | | | | | | | |
| AfFKB12/APX879 | Chain A | | | | | | | | | | | | | | | | | | | | | | | | | | | | | | | | |
| AfFKB12/APX879 | Chain B | | | | | | | | | | | | | | | | | | | | | | | | | | | | | | | | |
| McFKB12/APX879 | - | | | | | | | | | | | | | | | | | | | | | | | | | | | | | | | | |

**Selection criteria: the residues marked in gray are**
**within 4 angstrom distance from a crystallographic or pseudo-crystallographic symmetry-related protein molecule**

**Table S3. Hydrogen bonds observed during last 400ns of MD simulations between each protein-ligand complex (Excel document).**

Hydrogen bonds observed during last 400ns of MD simulations between *h* FKBP12 and FK506

| Residue # | residue atom | FK506 atom | # of times hbond observed | frequency of hbond's existence | Z-Score | Overall residue frequency of hbonds |
|---|---|---|---|---|---|---|
| 27 | OH | O4 | 66 | 0.330% | 0 | 1.310% |
| 27 | OH | O5 | 195 | 0.975% | 0 | |
| 27 | OH | O6 | 1 | 0.005% | 0 | |
| 29 | N | O6 | 1 | 0.005% | 0 | 0.005% |
| 37 | O | O6 | 32 | 0.160% | 0 | 0.160% |
| 38 | N | O4 | 11 | 0.055% | 0 | 21.800% |
| 38 | N | O6 | 500 | 2.499% | 0 | |
| 38 | OD1 | O4 | 14 | 0.070% | 0 | |
| 38 | OD1 | O5 | 123 | 0.615% | 0 | |
| 38 | OD1 | O6 | 358 | 1.790% | 0 | |
| 38 | OD1 | O7 | 4 | 0.020% | 0 | |
| 38 | OD2 | O4 | 14 | 0.070% | 0 | |
| 38 | OD2 | O5 | 151 | 0.755% | 0 | |
| 38 | OD2 | O6 | 392 | 1.960% | 0 | |
| 38 | OD2 | O7 | 1 | 0.005% | 0 | |
| 38 | O | O4 | 739 | 3.694% | 0 | |
| 38 | O | O5 | 41 | 0.205% | 0 | |
| 38 | O | O6 | 2013 | 10.063% | 1 | |
| 43 | NH1 | O5 | 1 | 0.005% | 0 | 1.445% |
| 43 | NH1 | O6 | 49 | 0.245% | 0 | |
| 43 | NH1 | O7 | 4 | 0.020% | 0 | |
| 43 | NH1 | O8 | 87 | 0.435% | 0 | |
| 43 | NH2 | O4 | 76 | 0.380% | 0 | |
| 43 | NH2 | O5 | 56 | 0.280% | 0 | |
| 43 | NH2 | O6 | 1 | 0.005% | 0 | |
| 43 | NH2 | O7 | 1 | 0.005% | 0 | |
| 43 | NH2 | O8 | 14 | 0.070% | 0 | |
| 44 | ND2 | O6 | 1 | 0.005% | 0 | 0.005% |
| 53 | NZ | O10 | 5 | 0.025% | 0 | 0.025% |
| 54 | NE2 | O10 | 1 | 0.005% | 0 | 0.350% |
| 54 | NE2 | O11 | 5 | 0.025% | 0 | |
| 54 | NE2 | O12 | 9 | 0.045% | 0 | |
| 54 | OE1 | O11 | 1 | 0.005% | 0 | |
| 54 | OE1 | O12 | 4 | 0.020% | 0 | |
| 54 | O | O10 | 1 | 0.005% | 0 | |
| 54 | O | O11 | 33 | 0.165% | 0 | |
| 54 | O | O12 | 16 | 0.080% | 0 | |
| 55 | N | O10 | 3 | 0.015% | 0 | |
| 55 | OE1 | O10 | 27 | 0.135% | 0 | |
| 55 | OE2 | O10 | 28 | 0.140% | 0 | |

Hydrogen bonds observed during last 400ns of MD simulations between *h* FKBP12 and FK506

| Residue # | residue atom | FK506 atom | # of times hbond observed | frequency of hbond's existence | Z-Score | Overall residue frequency of hbonds |
|---|---|---|---|---|---|---|
| 55 | OE2 | O12 | 1 | 0.005% | 0 | 34.636% |
| 55 | O | O10 | 6811 | 34.047% | 5 | |
| 55 | O | O11 | 32 | 0.160% | 0 | |
| 55 | O | O12 | 13 | 0.065% | 0 | |
| 55 | O | O2 | 14 | 0.070% | 0 | |
| 56 | N | O10 | 3 | 0.015% | 0 | |
| 56 | O | O11 | 94 | 0.470% | 0 | 0.510% |
| 56 | O | O12 | 5 | 0.025% | 0 | |
| 57 | N | O2 | 13660 | 68.283% | 9 | 68.283% |
| 58 | NH1 | O11 | 19 | 0.095% | 0 | 0.105% |
| 58 | NH2 | O11 | 2 | 0.010% | 0 | |
| 60 | NE1 | O11 | 2131 | 10.652% | 1 | |
| 60 | NE1 | O12 | 17 | 0.085% | 0 | |
| 60 | NE1 | O2 | 5 | 0.025% | 0 | 30.542% |
| 60 | NE1 | O3 | 3940 | 19.695% | 3 | |
| 60 | NE1 | O4 | 15 | 0.075% | 0 | |
| 60 | N | O12 | 2 | 0.010% | 0 | |
| 79 | O | O12 | 124 | 0.620% | 0 | 0.620% |
| 82 | O | O11 | 16 | 0.080% | 0 | 0.230% |
| 82 | O | O12 | 30 | 0.150% | 0 | |
| 83 | N | O11 | 1 | 0.005% | 0 | |
| 83 | N | O12 | 14 | 0.070% | 0 | |
| 83 | OH | O10 | 6 | 0.030% | 0 | |
| 83 | OH | O11 | 74 | 0.370% | 0 | 2.579% |
| 83 | OH | O12 | 306 | 1.530% | 0 | |
| 83 | OH | O3 | 3 | 0.015% | 0 | |
| 83 | O | O11 | 9 | 0.045% | 0 | |
| 83 | O | O12 | 103 | 0.515% | 0 | |
| 84 | N | O11 | 1658 | 8.288% | 1 | |
| 84 | N | O12 | 234 | 1.170% | 0 | 10.972% |
| 84 | O | O11 | 146 | 0.730% | 0 | |
| 84 | O | O12 | 157 | 0.785% | 0 | |
| 85 | N | O11 | 699 | 3.494% | 0 | |
| 85 | N | O12 | 297 | 1.485% | 0 | 12.787% |
| 85 | O | O11 | 93 | 0.465% | 0 | |
| 85 | O | O12 | 1469 | 7.343% | 1 | |
| 86 | N | O11 | 438 | 2.189% | 0 | |
| 86 | N | O12 | 309 | 1.545% | 0 | |
| 86 | OG1 | O11 | 567 | 2.834% | 0 | |
| 86 | OG1 | O12 | 885 | 4.424% | 0 | 12.617% |

Hydrogen bonds observed during last 400ns of MD simulations between *h* FKBP12 and FK506

| Residue # | residue atom | FK506 atom | # of times hbond observed | frequency of hbond's existence | Z-Score | Overall residue frequency of hbonds |
|---|---|---|---|---|---|---|
| 86 | O | O10 | 1 | 0.005% | 0 | |
| 86 | O | O11 | 24 | 0.120% | 0 | |
| 86 | O | O12 | 300 | 1.500% | 0 | |
| 87 | N | O11 | 89 | 0.445% | 0 | |
| 87 | N | O12 | 283 | 1.415% | 0 | |
| 87 | O | O10 | 1 | 0.005% | 0 | 3.849% |
| 87 | O | O11 | 29 | 0.145% | 0 | |
| 87 | O | O12 | 368 | 1.840% | 0 | |
| 88 | ND1 | O10 | 7 | 0.035% | 0 | |
| 88 | ND1 | O12 | 54 | 0.270% | 0 | |
| 88 | ND1 | O9 | 1 | 0.005% | 0 | |
| 88 | NE2 | O10 | 254 | 1.270% | 0 | |
| 88 | NE2 | O11 | 50 | 0.250% | 0 | |
| 88 | NE2 | O12 | 47 | 0.235% | 0 | |
| 88 | NE2 | O9 | 71 | 0.355% | 0 | |
| 88 | N | O10 | 2 | 0.010% | 0 | 11.972% |
| 88 | N | O11 | 59 | 0.295% | 0 | |
| 88 | N | O12 | 466 | 2.329% | 0 | |
| 88 | N | O9 | 2 | 0.010% | 0 | |
| 88 | O | O10 | 40 | 0.200% | 0 | |
| 88 | O | O11 | 62 | 0.310% | 0 | |
| 88 | O | O12 | 1258 | 6.288% | 1 | |
| 88 | O | O9 | 22 | 0.110% | 0 | |
| 89 | N | O11 | 3 | 0.015% | 0 | |
| 89 | N | O12 | 6 | 0.030% | 0 | |
| 89 | O | O10 | 122 | 0.610% | 0 | |
| 89 | O | O11 | 13 | 0.065% | 0 | 1.140% |
| 89 | O | O12 | 31 | 0.155% | 0 | |
| 89 | O | O7 | 3 | 0.015% | 0 | |
| 89 | O | O9 | 50 | 0.250% | 0 | |
| 90 | N | O11 | 19 | 0.095% | 0 | |
| 90 | N | O12 | 125 | 0.625% | 0 | |
| 90 | N | O7 | 4 | 0.020% | 0 | |
| 90 | N | O9 | 32 | 0.160% | 0 | 1.020% |
| 90 | O | O11 | 11 | 0.055% | 0 | |
| 90 | O | O12 | 9 | 0.045% | 0 | |
| 90 | O | O7 | 3 | 0.015% | 0 | |
| 90 | O | O9 | 1 | 0.005% | 0 | |
| 91 | N | O11 | 15 | 0.075% | 0 | |
| 91 | N | O12 | 1 | 0.005% | 0 | |
| 91 | N | O7 | 282 | 1.410% | 0 | |

Hydrogen bonds observed during last 400ns of MD simulations between *h* FKBP12 and FK506

| Residue # | residue atom | FK506 atom | # of times hbond observed | frequency of hbond's existence | Z-Score | Overall residue frequency of hbonds |
|---|---|---|---|---|---|---|
| 91 | N | O9 | 1178 | 5.889% | 1 | 7.478% |
| 91 | O | O12 | 11 | 0.055% | 0 | |
| 91 | O | O3 | 1 | 0.005% | 0 | |
| 91 | O | O7 | 7 | 0.035% | 0 | |
| 91 | O | O9 | 1 | 0.005% | 0 | |
| 93 | O | O11 | 1 | 0.005% | 0 | 0.005% |
| 95 | NE2 | O11 | 4 | 0.020% | 0 | 0.020% |

Hydrogen bonds observed during last 400ns of MD simulations between *h* FKBP12 and APX879

| Residue # | residue atom | APX879 atom | # of times hbond observed | frequency of hbond's existence | Z-Score | Overall residue frequency of hbonds |
|---|---|---|---|---|---|---|
| 27 | OH | O4 | 14 | 0.070% | 0 | |
| 27 | OH | O5 | 5 | 0.025% | 0 | |
| 27 | OH | O6 | 161 | 0.805% | 0 | 3.574% |
| 27 | OH | O8 | 530 | 2.649% | 0 | |
| 27 | OH | O13 | 5 | 0.025% | 0 | |
| 28 | O | O4 | 3 | 0.015% | 0 | 0.015% |
| 37 | O | O6 | 3 | 0.015% | 0 | 0.015% |
| 38 | N | O4 | 1 | 0.005% | 0 | |
| 38 | N | O6 | 19 | 0.095% | 0 | |
| 38 | N | O8 | 3 | 0.015% | 0 | |
| 38 | OD1 | O4 | 113 | 0.565% | 0 | |
| 38 | OD1 | O6 | 3129 | 15.641% | 5 | |
| 38 | OD1 | O8 | 29 | 0.145% | 0 | |
| 38 | OD2 | O4 | 129 | 0.645% | 0 | 39.745% |
| 38 | OD2 | O5 | 3 | 0.015% | 0 | |
| 38 | OD2 | O6 | 3356 | 16.776% | 5 | |
| 38 | OD2 | O8 | 29 | 0.145% | 0 | |
| 38 | O | O4 | 232 | 1.160% | 0 | |
| 38 | O | O5 | 1 | 0.005% | 0 | |
| 38 | O | O6 | 907 | 4.534% | 1 | |
| 39 | OG | O6 | 1 | 0.005% | 0 | 0.005% |
| 43 | NE | O6 | 19 | 0.095% | 0 | |
| 43 | NE | O8 | 479 | 2.394% | 0 | |
| 43 | NE | O13 | 10 | 0.050% | 0 | |
| 43 | NH1 | O4 | 523 | 2.614% | 0 | |
| 43 | NH1 | O5 | 746 | 3.729% | 1 | |
| 43 | NH1 | O6 | 868 | 4.339% | 1 | |
| 43 | NH1 | O7 | 3 | 0.015% | 0 | |
| 43 | NH1 | O8 | 2323 | 11.612% | 3 | |
| 43 | NH1 | O10 | 1 | 0.005% | 0 | 55.281% |
| 43 | NH1 | O13 | 24 | 0.120% | 0 | |
| 43 | NH2 | N54 | 1 | 0.005% | 0 | |
| 43 | NH2 | O4 | 22 | 0.110% | 0 | |
| 43 | NH2 | O5 | 53 | 0.265% | 0 | |
| 43 | NH2 | O6 | 3781 | 18.900% | 6 | |
| 43 | NH2 | O7 | 21 | 0.105% | 0 | |
| 43 | NH2 | O8 | 2176 | 10.877% | 3 | |
| 43 | NH2 | O13 | 9 | 0.045% | 0 | |
| 48 | O | O10 | 24 | 0.120% | 0 | |

Hydrogen bonds observed during last 400ns of MD simulations between *h* FKBP12 and APX879

| Residue # | residue atom | APX879 atom | # of times hbond observed | frequency of hbond's existence | Z-Score | Overall residue frequency of hbonds |
|---|---|---|---|---|---|---|
| 48 | O | O11 | 2 | 0.010% | 0 | 1.185% |
| 48 | O | O12 | 101 | 0.505% | 0 | |
| 48 | O | O13 | 110 | 0.550% | 0 | |
| 50 | N | O10 | 354 | 1.770% | 0 | 11.537% |
| 50 | N | O11 | 4 | 0.020% | 0 | |
| 50 | N | O12 | 269 | 1.345% | 0 | |
| 50 | N | O13 | 1663 | 8.313% | 2 | |
| 50 | O | O10 | 5 | 0.025% | 0 | |
| 50 | O | O12 | 1 | 0.005% | 0 | |
| 50 | O | O13 | 12 | 0.060% | 0 | |
| 52 | O | N55 | 1 | 0.005% | 0 | 0.005% |
| 53 | NZ | O10 | 1 | 0.005% | 0 | 1.510% |
| 53 | NZ | O11 | 4 | 0.020% | 0 | |
| 53 | NZ | O12 | 15 | 0.075% | 0 | |
| 53 | NZ | O13 | 137 | 0.685% | 0 | |
| 53 | O | N55 | 144 | 0.720% | 0 | |
| 53 | O | O10 | 1 | 0.005% | 0 | |
| 54 | NE2 | N55 | 4 | 0.020% | 0 | 1.365% |
| 54 | NE2 | O10 | 14 | 0.070% | 0 | |
| 54 | NE2 | O11 | 1 | 0.005% | 0 | |
| 54 | NE2 | O12 | 3 | 0.015% | 0 | |
| 54 | NE2 | O13 | 39 | 0.195% | 0 | |
| 54 | OE1 | N54 | 3 | 0.015% | 0 | |
| 54 | OE1 | N55 | 161 | 0.805% | 0 | |
| 54 | OE1 | O10 | 3 | 0.015% | 0 | |
| 54 | OE1 | O12 | 1 | 0.005% | 0 | |
| 54 | OE1 | O13 | 13 | 0.065% | 0 | |
| 54 | O | N55 | 1 | 0.005% | 0 | |
| 54 | O | O10 | 4 | 0.020% | 0 | |
| 54 | O | O11 | 18 | 0.090% | 0 | |
| 54 | O | O12 | 8 | 0.040% | 0 | |
| 55 | OE1 | N55 | 120 | 0.600% | 0 | 26.458% |
| 55 | OE1 | O10 | 102 | 0.510% | 0 | |
| 55 | OE1 | O11 | 1 | 0.005% | 0 | |
| 55 | OE1 | O12 | 574 | 2.869% | 1 | |
| 55 | OE1 | O13 | 6 | 0.030% | 0 | |
| 55 | OE2 | N55 | 97 | 0.485% | 0 | |
| 55 | OE2 | O10 | 81 | 0.405% | 0 | |
| 55 | OE2 | O12 | 579 | 2.894% | 1 | |

Hydrogen bonds observed during last 400ns of MD simulations between *h* FKBP12 and APX879

| Residue # | residue atom | APX879 atom | # of times hbond observed | frequency of hbond's existence | Z-Score | Overall residue frequency of hbonds |
|---|---|---|---|---|---|---|
| 55 | OE2 | O13 | 4 | 0.020% | 0 | |
| 55 | O | N55 | 197 | 0.985% | 0 | |
| 55 | O | O2 | 61 | 0.305% | 0 | |
| 55 | O | O10 | 3387 | 16.931% | 5 | |
| 55 | O | O11 | 78 | 0.390% | 0 | |
| 55 | O | O12 | 6 | 0.030% | 0 | |
| 56 | N | O10 | 61 | 0.305% | 0 | |
| 56 | O | O11 | 10 | 0.050% | 0 | 0.360% |
| 56 | O | O12 | 1 | 0.005% | 0 | |
| 57 | N | O2 | 2567 | 12.832% | 4 | 13.112% |
| 57 | N | O10 | 56 | 0.280% | 0 | |
| 58 | NH1 | O11 | 7 | 0.035% | 0 | |
| 58 | NH1 | O12 | 3 | 0.015% | 0 | 0.060% |
| 58 | NH2 | O11 | 2 | 0.010% | 0 | |
| 60 | NE1 | N45 | 1 | 0.005% | 0 | |
| 60 | NE1 | O2 | 4 | 0.020% | 0 | |
| 60 | NE1 | O3 | 963 | 4.814% | 1 | 6.758% |
| 60 | NE1 | O4 | 383 | 1.915% | 0 | |
| 60 | NE1 | O12 | 1 | 0.005% | 0 | |
| 81 | OH | O2 | 26 | 0.130% | 0 | |
| 81 | OH | O3 | 30 | 0.150% | 0 | |
| 81 | OH | O11 | 1 | 0.005% | 0 | 0.655% |
| 81 | OH | O12 | 1 | 0.005% | 0 | |
| 81 | O | O11 | 7 | 0.035% | 0 | |
| 81 | O | O12 | 66 | 0.330% | 0 | |
| 82 | O | O3 | 1 | 0.005% | 0 | |
| 82 | O | O7 | 1 | 0.005% | 0 | 0.165% |
| 82 | O | O11 | 1 | 0.005% | 0 | |
| 82 | O | O12 | 30 | 0.150% | 0 | |
| 83 | OH | O3 | 14 | 0.070% | 0 | |
| 83 | OH | O11 | 9 | 0.045% | 0 | |
| 83 | OH | O12 | 108 | 0.540% | 0 | 1.065% |
| 83 | O | O11 | 19 | 0.095% | 0 | |
| 83 | O | O12 | 63 | 0.315% | 0 | |
| 84 | N | O11 | 72 | 0.360% | 0 | |
| 84 | N | O12 | 49 | 0.245% | 0 | 1.170% |
| 84 | O | O11 | 34 | 0.170% | 0 | |
| 84 | O | O12 | 79 | 0.395% | 0 | |
| 85 | N | O11 | 140 | 0.700% | 0 | |

Hydrogen bonds observed during last 400ns of MD simulations between *h* FKBP12 and APX879

| Residue # | residue atom | APX879 atom | # of times hbond observed | frequency of hbond's existence | Z-Score | Overall residue frequency of hbonds |
|---|---|---|---|---|---|---|
| 85 | N | O12 | 225 | 1.125% | 0 | 2.949% |
| 85 | O | O7 | 4 | 0.020% | 0 | |
| 85 | O | O11 | 123 | 0.615% | 0 | |
| 85 | O | O12 | 98 | 0.490% | 0 | |
| 86 | N | O11 | 38 | 0.190% | 0 | 9.323% |
| 86 | N | O12 | 421 | 2.104% | 0 | |
| 86 | OG1 | O10 | 25 | 0.125% | 0 | |
| 86 | OG1 | O11 | 186 | 0.930% | 0 | |
| 86 | OG1 | O12 | 1016 | 5.079% | 1 | |
| 86 | O | N55 | 14 | 0.070% | 0 | |
| 86 | O | O7 | 4 | 0.020% | 0 | |
| 86 | O | O10 | 25 | 0.125% | 0 | |
| 86 | O | O11 | 7 | 0.035% | 0 | |
| 86 | O | O12 | 123 | 0.615% | 0 | |
| 86 | O | O13 | 6 | 0.030% | 0 | |
| 87 | N | O7 | 53 | 0.265% | 0 | 9.933% |
| 87 | N | O11 | 35 | 0.175% | 0 | |
| 87 | N | O12 | 1493 | 7.463% | 2 | |
| 87 | O | O10 | 1 | 0.005% | 0 | |
| 87 | O | O11 | 67 | 0.335% | 0 | |
| 87 | O | O12 | 338 | 1.690% | 0 | |
| 88 | ND1 | N55 | 1 | 0.005% | 0 | 6.353% |
| 88 | ND1 | O10 | 3 | 0.015% | 0 | |
| 88 | ND1 | O12 | 32 | 0.160% | 0 | |
| 88 | NE2 | O7 | 5 | 0.025% | 0 | |
| 88 | NE2 | O10 | 7 | 0.035% | 0 | |
| 88 | NE2 | O11 | 6 | 0.030% | 0 | |
| 88 | NE2 | O12 | 63 | 0.315% | 0 | |
| 88 | NE2 | O13 | 3 | 0.015% | 0 | |
| 88 | N | O7 | 114 | 0.570% | 0 | |
| 88 | N | O10 | 37 | 0.185% | 0 | |
| 88 | N | O11 | 54 | 0.270% | 0 | |
| 88 | N | O12 | 229 | 1.145% | 0 | |
| 88 | N | O13 | 2 | 0.010% | 0 | |
| 88 | O | O11 | 341 | 1.705% | 0 | |
| 88 | O | O12 | 374 | 1.870% | 0 | |
| 89 | N | O11 | 11 | 0.055% | 0 | 1.130% |
| 89 | N | O12 | 1 | 0.005% | 0 | |
| 89 | O | O7 | 11 | 0.055% | 0 | |
| 89 | O | O11 | 8 | 0.040% | 0 | |

Hydrogen bonds observed during last 400ns of MD simulations between *h* FKBP12 and APX879

| Residue # | residue atom | APX879 atom | # of times hbond observed | frequency of hbond's existence | Z-Score | Overall residue frequency of hbonds |
|---|---|---|---|---|---|---|
| 89 | O | O12 | 195 | 0.975% | 0 | |
| 90 | N | O3 | 1 | 0.005% | 0 | 0.945% |
| 90 | N | O11 | 64 | 0.320% | 0 | |
| 90 | N | O12 | 95 | 0.475% | 0 | |
| 90 | O | O6 | 2 | 0.010% | 0 | |
| 90 | O | O7 | 20 | 0.100% | 0 | |
| 90 | O | O12 | 7 | 0.035% | 0 | |
| 91 | N | O7 | 77 | 0.385% | 0 | 0.885% |
| 91 | N | O8 | 3 | 0.015% | 0 | |
| 91 | N | O11 | 3 | 0.015% | 0 | |
| 91 | N | O12 | 9 | 0.045% | 0 | |
| 91 | O | O6 | 32 | 0.160% | 0 | |
| 91 | O | O7 | 39 | 0.195% | 0 | |
| 91 | O | O8 | 5 | 0.025% | 0 | |
| 91 | O | O12 | 9 | 0.045% | 0 | |

Hydrogen bonds observed during last 400ns of MD simulations between *Af*FKBP12 and FK506

| Residue # | residue atom | FK506 atom | # of times hbond observed | frequency of hbond's existence | Z-SCORE | Overall residue frequency of hbonds |
|---|---|---|---|---|---|---|
| 27 | OH | O4 | 110 | 0.458% | 0 | |
| 27 | OH | O5 | 712 | 2.966% | 0 | 3.453% |
| 27 | OH | O6 | 7 | 0.029% | 0 | |
| 28 | O | O4 | 2 | 0.008% | 0 | 0.008% |
| 37 | O | O6 | 34 | 0.142% | 0 | 0.142% |
| 38 | N | O4 | 210 | 0.875% | 0 | |
| 38 | N | O6 | 5138 | 21.403% | 3 | |
| 38 | OD1 | O4 | 29 | 0.121% | 0 | |
| 38 | OD1 | O5 | 49 | 0.204% | 0 | |
| 38 | OD1 | O6 | 933 | 3.887% | 0 | |
| 38 | OD1 | O7 | 1 | 0.004% | 0 | |
| 38 | OD2 | O4 | 33 | 0.137% | 0 | 47.347% |
| 38 | OD2 | O5 | 53 | 0.221% | 0 | |
| 38 | OD2 | O6 | 828 | 3.449% | 0 | |
| 38 | OD2 | O7 | 2 | 0.008% | 0 | |
| 38 | O | O4 | 1400 | 5.832% | 1 | |
| 38 | O | O5 | 317 | 1.321% | 0 | |
| 38 | O | O6 | 2373 | 9.885% | 1 | |
| 43 | NE | O7 | 2 | 0.008% | 0 | |
| 43 | NH1 | O4 | 267 | 1.112% | 0 | |
| 43 | NH1 | O5 | 102 | 0.425% | 0 | |
| 43 | NH1 | O6 | 4 | 0.017% | 0 | |
| 43 | NH1 | O7 | 25 | 0.104% | 0 | |
| 43 | NH1 | O8 | 13 | 0.054% | 0 | 2.391% |
| 43 | NH2 | O4 | 13 | 0.054% | 0 | |
| 43 | NH2 | O5 | 9 | 0.037% | 0 | |
| 43 | NH2 | O6 | 110 | 0.458% | 0 | |
| 43 | NH2 | O7 | 22 | 0.092% | 0 | |
| 43 | NH2 | O8 | 7 | 0.029% | 0 | |
| 49 | OG1 | O2 | 3 | 0.012% | 0 | 0.012% |
| 54 | N | O11 | 43 | 0.179% | 0 | |
| 54 | O | O10 | 17 | 0.071% | 0 | 2.970% |
| 54 | O | O11 | 54 | 0.225% | 0 | |
| 54 | O | O12 | 599 | 2.495% | 0 | |
| 55 | NE | O10 | 25 | 0.104% | 0 | |
| 55 | NE | O11 | 24 | 0.100% | 0 | |
| 55 | NE | O12 | 35 | 0.146% | 0 | |
| 55 | NH1 | O10 | 68 | 0.283% | 0 | |
| 55 | NH1 | O11 | 91 | 0.379% | 0 | |
| 55 | NH1 | O12 | 93 | 0.387% | 0 | |
| 55 | NH1 | O9 | 2 | 0.008% | 0 | |

Hydrogen bonds observed during last 400ns of MD simulations between *Af*FKBP12 and FK506

| Residue # | residue atom | FK506 atom | # of times hbond observed | frequency of hbond's existence | Z-SCORE | Overall residue frequency of hbonds |
|---|---|---|---|---|---|---|
| 55 | NH2 | O10 | 15 | 0.062% | 0 | 34.866% |
| 55 | NH2 | O11 | 30 | 0.125% | 0 | |
| 55 | NH2 | O12 | 36 | 0.150% | 0 | |
| 55 | NH2 | O9 | 3 | 0.012% | 0 | |
| 55 | O | O10 | 7787 | 32.438% | 4 | |
| 55 | O | O11 | 78 | 0.325% | 0 | |
| 55 | O | O12 | 64 | 0.267% | 0 | |
| 55 | O | O2 | 19 | 0.079% | 0 | |
| 56 | N | O10 | 5 | 0.021% | 0 | 0.062% |
| 56 | N | O11 | 1 | 0.004% | 0 | |
| 56 | O | O11 | 6 | 0.025% | 0 | |
| 56 | O | O12 | 3 | 0.012% | 0 | |
| 57 | N | O2 | 17050 | 71.024% | 10 | 71.024% |
| 58 | NZ | O11 | 2 | 0.008% | 0 | 0.008% |
| 60 | NE1 | O3 | 1733 | 7.219% | 1 | 7.244% |
| 60 | NE1 | O4 | 6 | 0.025% | 0 | |
| 82 | O | O11 | 8 | 0.033% | 0 | 0.100% |
| 82 | O | O12 | 16 | 0.067% | 0 | |
| 83 | OH | O1 | 24 | 0.100% | 0 | 5.345% |
| 83 | OH | O11 | 20 | 0.083% | 0 | |
| 83 | OH | O12 | 95 | 0.396% | 0 | |
| 83 | OH | O2 | 13 | 0.054% | 0 | |
| 83 | OH | O3 | 883 | 3.678% | 0 | |
| 83 | OH | O6 | 17 | 0.071% | 0 | |
| 83 | O | O11 | 30 | 0.125% | 0 | |
| 83 | O | O12 | 201 | 0.837% | 0 | |
| 84 | N | O11 | 947 | 3.945% | 0 | 5.361% |
| 84 | N | O12 | 149 | 0.621% | 0 | |
| 84 | O | O11 | 100 | 0.417% | 0 | |
| 84 | O | O12 | 91 | 0.379% | 0 | |
| 85 | N | O11 | 458 | 1.908% | 0 | 8.319% |
| 85 | N | O12 | 501 | 2.087% | 0 | |
| 85 | O | O10 | 9 | 0.037% | 0 | |
| 85 | O | O11 | 140 | 0.583% | 0 | |
| 85 | O | O12 | 889 | 3.703% | 0 | |
| 86 | NE | O11 | 138 | 0.575% | 0 | |
| 86 | NE | O12 | 73 | 0.304% | 0 | |
| 86 | NE | O7 | 67 | 0.279% | 0 | |
| 86 | NH1 | O11 | 114 | 0.475% | 0 | |
| 86 | NH1 | O12 | 117 | 0.487% | 0 | |
| 86 | NH1 | O6 | 5 | 0.021% | 0 | |

Hydrogen bonds observed during last 400ns of MD simulations between *Af* FKBP12 and FK506

| Residue # | residue atom | FK506 atom | # of times hbond observed | frequency of hbond's existence | Z-SCORE | Overall residue frequency of hbonds |
|---|---|---|---|---|---|---|
| 86 | NH1 | O7 | 46 | 0.192% | 0 | |
| 86 | NH1 | O8 | 2 | 0.008% | 0 | |
| 86 | NH1 | O9 | 2 | 0.008% | 0 | |
| 86 | NH2 | O10 | 1 | 0.004% | 0 | |
| 86 | NH2 | O11 | 115 | 0.479% | 0 | 10.027% |
| 86 | NH2 | O12 | 34 | 0.142% | 0 | |
| 86 | NH2 | O6 | 11 | 0.046% | 0 | |
| 86 | NH2 | O7 | 34 | 0.142% | 0 | |
| 86 | NH2 | O9 | 1 | 0.004% | 0 | |
| 86 | N | O11 | 60 | 0.250% | 0 | |
| 86 | N | O12 | 158 | 0.658% | 0 | |
| 86 | O | O10 | 91 | 0.379% | 0 | |
| 86 | O | O11 | 58 | 0.242% | 0 | |
| 86 | O | O12 | 1274 | 5.307% | 1 | |
| 86 | O | O7 | 6 | 0.025% | 0 | |
| 87 | N | O11 | 79 | 0.329% | 0 | |
| 87 | N | O12 | 250 | 1.041% | 0 | |
| 87 | N | O7 | 1 | 0.004% | 0 | 3.695% |
| 87 | O | O11 | 41 | 0.171% | 0 | |
| 87 | O | O12 | 516 | 2.149% | 0 | |
| 88 | N | O11 | 283 | 1.179% | 0 | |
| 88 | N | O12 | 106 | 0.442% | 0 | |
| 88 | N | O9 | 150 | 0.625% | 0 | |
| 88 | O | O10 | 156 | 0.650% | 0 | 7.082% |
| 88 | O | O11 | 121 | 0.504% | 0 | |
| 88 | O | O12 | 877 | 3.653% | 0 | |
| 88 | O | O9 | 7 | 0.029% | 0 | |
| 89 | N | O12 | 4 | 0.017% | 0 | |
| 89 | N | O9 | 3 | 0.012% | 0 | |
| 89 | O | O10 | 70 | 0.292% | 0 | |
| 89 | O | O11 | 66 | 0.275% | 0 | |
| 89 | O | O12 | 75 | 0.312% | 0 | 1.616% |
| 89 | O | O6 | 11 | 0.046% | 0 | |
| 89 | O | O7 | 13 | 0.054% | 0 | |
| 89 | O | O9 | 146 | 0.608% | 0 | |
| 90 | N | O11 | 93 | 0.387% | 0 | |
| 90 | N | O12 | 14 | 0.058% | 0 | |
| 90 | N | O6 | 1 | 0.004% | 0 | |
| 90 | N | O7 | 1 | 0.004% | 0 | |
| 90 | N | O9 | 12 | 0.050% | 0 | |
| 90 | O | O11 | 18 | 0.075% | 0 | 0.683% |

Hydrogen bonds observed during last 400ns of MD simulations between *Af* FKBP12 and FK506

| Residue # | residue atom | FK506 atom | # of times hbond observed | frequency of hbond's existence | Z-SCORE | Overall residue frequency of hbonds |
|---|---|---|---|---|---|---|
| 90 | O | O12 | 10 | 0.042% | 0 | |
| 90 | O | O3 | 1 | 0.004% | 0 | |
| 90 | O | O6 | 12 | 0.050% | 0 | |
| 90 | O | O7 | 1 | 0.004% | 0 | |
| 90 | O | O9 | 1 | 0.004% | 0 | |
| 91 | N | O11 | 41 | 0.171% | 0 | |
| 91 | N | O3 | 4 | 0.017% | 0 | |
| 91 | N | O6 | 3 | 0.012% | 0 | |
| 91 | N | O7 | 49 | 0.204% | 0 | |
| 91 | N | O9 | 67 | 0.279% | 0 | 0.921% |
| 91 | O | O11 | 2 | 0.008% | 0 | |
| 91 | O | O12 | 2 | 0.008% | 0 | |
| 91 | O | O6 | 51 | 0.212% | 0 | |
| 91 | O | O7 | 2 | 0.008% | 0 | |
| 92 | N | O11 | 4 | 0.017% | 0 | |
| 92 | N | O6 | 1 | 0.004% | 0 | 0.025% |
| 92 | O | O11 | 1 | 0.004% | 0 | |

Hydrogen bonds observed during last 400ns of MD simulations between *Af*FKBP12 and APX879

| Residue # | residue atom | APX879 atom | # of times hbond observed | frequency of hbond's existence | Z-SCORE | Overall residue frequency of hbonds |
|---|---|---|---|---|---|---|
| 27 | OH | O4 | 95 | 0.475% | 0 | |
| 27 | OH | O5 | 6 | 0.030% | 0 | |
| 27 | OH | O6 | 1987 | 9.933% | 2 | 13.837% |
| 27 | OH | O8 | 679 | 3.394% | 0 | |
| 27 | O | O4 | 1 | 0.005% | 0 | |
| 37 | O | O4 | 1 | 0.005% | 0 | 0.005% |
| 38 | N | O4 | 1 | 0.005% | 0 | |
| 38 | N | O6 | 18 | 0.090% | 0 | |
| 38 | OD1 | O4 | 189 | 0.945% | 0 | |
| 38 | OD1 | O6 | 6167 | 30.827% | 6 | |
| 38 | OD1 | O7 | 1 | 0.005% | 0 | |
| 38 | OD2 | O4 | 172 | 0.860% | 0 | 72.287% |
| 38 | OD2 | O6 | 5759 | 28.788% | 6 | |
| 38 | OD2 | O7 | 2 | 0.010% | 0 | |
| 38 | O | O4 | 34 | 0.170% | 0 | |
| 38 | O | O6 | 2115 | 10.572% | 2 | |
| 38 | O | O8 | 3 | 0.015% | 0 | |
| 39 | N | O6 | 6 | 0.030% | 0 | |
| 39 | OG | O6 | 2 | 0.010% | 0 | |
| 39 | OG | O7 | 1 | 0.005% | 0 | 0.210% |
| 39 | OG | O8 | 4 | 0.020% | 0 | |
| 39 | O | O6 | 26 | 0.130% | 0 | |
| 39 | O | O8 | 3 | 0.015% | 0 | |
| 40 | OG | O4 | 3 | 0.015% | 0 | 0.060% |
| 40 | OG | O6 | 9 | 0.045% | 0 | |
| 43 | NE | O6 | 9 | 0.045% | 0 | |
| 43 | NE | O8 | 3 | 0.015% | 0 | |
| 43 | NH1 | O5 | 5 | 0.025% | 0 | |
| 43 | NH1 | O6 | 50 | 0.250% | 0 | |
| 43 | NH1 | O8 | 67 | 0.335% | 0 | |
| 43 | NH1 | O13 | 6 | 0.030% | 0 | 1.795% |
| 43 | NH2 | O5 | 5 | 0.025% | 0 | |
| 43 | NH2 | O6 | 138 | 0.690% | 0 | |
| 43 | NH2 | O7 | 3 | 0.015% | 0 | |
| 43 | NH2 | O8 | 66 | 0.330% | 0 | |
| 43 | NH2 | O13 | 7 | 0.035% | 0 | |
| 48 | O | N54 | 1 | 0.005% | 0 | |
| 48 | O | N55 | 12 | 0.060% | 0 | |
| 48 | O | O10 | 1 | 0.005% | 0 | 0.260% |
| 48 | O | O12 | 6 | 0.030% | 0 | |

Hydrogen bonds observed during last 400ns of MD simulations between *Af*FKBP12 and APX879

| Residue # | residue atom | APX879 atom | # of times hbond observed | frequency of hbond's existence | Z-SCORE | Overall residue frequency of hbonds |
|---|---|---|---|---|---|---|
| 48 | O | O13 | 32 | 0.160% | 0 | |
| 49 | OG1 | N54 | 6 | 0.030% | 0 | |
| 49 | OG1 | N55 | 194 | 0.970% | 0 | |
| 49 | OG1 | O2 | 5 | 0.025% | 0 | 2.129% |
| 49 | OG1 | O10 | 172 | 0.860% | 0 | |
| 49 | OG1 | O13 | 49 | 0.245% | 0 | |
| 50 | NE2 | N55 | 1 | 0.005% | 0 | |
| 50 | NE2 | O10 | 14 | 0.070% | 0 | |
| 50 | NE2 | O13 | 10 | 0.050% | 0 | |
| 50 | N | O10 | 47 | 0.235% | 0 | |
| 50 | N | O12 | 12 | 0.060% | 0 | 4.369% |
| 50 | N | O13 | 723 | 3.614% | 0 | |
| 50 | OE1 | N55 | 47 | 0.235% | 0 | |
| 50 | OE1 | O10 | 13 | 0.065% | 0 | |
| 50 | OE1 | O13 | 3 | 0.015% | 0 | |
| 50 | O | O10 | 4 | 0.020% | 0 | |
| 54 | O | O10 | 45 | 0.225% | 0 | |
| 54 | O | O11 | 46 | 0.230% | 0 | 0.975% |
| 54 | O | O12 | 104 | 0.520% | 0 | |
| 55 | NE | N54 | 1 | 0.005% | 0 | |
| 55 | NE | N55 | 1 | 0.005% | 0 | |
| 55 | NE | O10 | 11 | 0.055% | 0 | |
| 55 | NE | O11 | 14 | 0.070% | 0 | |
| 55 | NE | O12 | 2 | 0.010% | 0 | |
| 55 | NE | O13 | 44 | 0.220% | 0 | |
| 55 | NH1 | N54 | 4 | 0.020% | 0 | |
| 55 | NH1 | N55 | 2 | 0.010% | 0 | |
| 55 | NH1 | O10 | 5 | 0.025% | 0 | |
| 55 | NH1 | O11 | 12 | 0.060% | 0 | |
| 55 | NH1 | O12 | 23 | 0.115% | 0 | |
| 55 | NH1 | O13 | 149 | 0.745% | 0 | 30.647% |
| 55 | NH2 | N54 | 1 | 0.005% | 0 | |
| 55 | NH2 | N55 | 1 | 0.005% | 0 | |
| 55 | NH2 | O10 | 2 | 0.010% | 0 | |
| 55 | NH2 | O11 | 5 | 0.025% | 0 | |
| 55 | NH2 | O12 | 18 | 0.090% | 0 | |
| 55 | NH2 | O13 | 61 | 0.305% | 0 | |
| 55 | N | O10 | 22 | 0.110% | 0 | |
| 55 | O | N55 | 209 | 1.045% | 0 | |
| 55 | O | O10 | 5513 | 27.558% | 5 | |

Hydrogen bonds observed during last 400ns of MD simulations between *Af*FKBP12 and APX879

| Residue # | residue atom | APX879 atom | # of times hbond observed | frequency of hbond's existence | Z-SCORE | Overall residue frequency of hbonds |
|---|---|---|---|---|---|---|
| 55 | O | O11 | 24 | 0.120% | 0 | |
| 55 | O | O12 | 7 | 0.035% | 0 | |
| 56 | N | O10 | 128 | 0.640% | 0 | 0.685% |
| 56 | O | O11 | 9 | 0.045% | 0 | |
| 57 | N | O2 | 5508 | 27.533% | 5 | 27.563% |
| 57 | N | O10 | 6 | 0.030% | 0 | |
| 60 | NE1 | O2 | 1 | 0.005% | 0 | |
| 60 | NE1 | O3 | 921 | 4.604% | 1 | 4.754% |
| 60 | NE1 | O4 | 29 | 0.145% | 0 | |
| 82 | O | O11 | 1 | 0.005% | 0 | 0.020% |
| 82 | O | O12 | 3 | 0.015% | 0 | |
| 83 | OH | N55 | 7 | 0.035% | 0 | |
| 83 | OH | O2 | 7 | 0.035% | 0 | |
| 83 | OH | O3 | 343 | 1.715% | 0 | |
| 83 | OH | O7 | 21 | 0.105% | 0 | |
| 83 | OH | O10 | 22 | 0.110% | 0 | 3.059% |
| 83 | OH | O11 | 99 | 0.495% | 0 | |
| 83 | OH | O12 | 105 | 0.525% | 0 | |
| 83 | OH | O13 | 3 | 0.015% | 0 | |
| 83 | O | O11 | 2 | 0.010% | 0 | |
| 83 | O | O12 | 3 | 0.015% | 0 | |
| 84 | N | O11 | 2 | 0.010% | 0 | |
| 84 | N | O12 | 4 | 0.020% | 0 | 0.130% |
| 84 | O | O12 | 20 | 0.100% | 0 | |
| 85 | N | O12 | 1 | 0.005% | 0 | |
| 85 | O | O11 | 12 | 0.060% | 0 | 0.835% |
| 85 | O | O12 | 154 | 0.770% | 0 | |
| 86 | NE | O10 | 3 | 0.015% | 0 | |
| 86 | NE | O11 | 79 | 0.395% | 0 | |
| 86 | NE | O12 | 226 | 1.130% | 0 | |
| 86 | NH1 | O7 | 1 | 0.005% | 0 | |
| 86 | NH1 | O10 | 12 | 0.060% | 0 | |
| 86 | NH1 | O11 | 216 | 1.080% | 0 | |
| 86 | NH1 | O12 | 726 | 3.629% | 0 | |
| 86 | NH2 | O10 | 2 | 0.010% | 0 | 8.358% |
| 86 | NH2 | O11 | 125 | 0.625% | 0 | |
| 86 | NH2 | O12 | 236 | 1.180% | 0 | |
| 86 | N | O12 | 3 | 0.015% | 0 | |
| 86 | O | O2 | 2 | 0.010% | 0 | |
| 86 | O | O7 | 2 | 0.010% | 0 | |

Hydrogen bonds observed during last 400ns of MD simulations between *Af* FKBP12 and APX879

| Residue # | residue atom | APX879 atom | # of times hbond observed | frequency of hbond's existence | Z-SCORE | Overall residue frequency of hbonds |
|---|---|---|---|---|---|---|
| 86 | O | O11 | 10 | 0.050% | 0 | |
| 86 | O | O12 | 29 | 0.145% | 0 | |
| 87 | N | O11 | 148 | 0.740% | 0 | 4.134% |
| 87 | N | O12 | 276 | 1.380% | 0 | |
| 87 | O | O11 | 154 | 0.770% | 0 | |
| 87 | O | O12 | 249 | 1.245% | 0 | |
| 88 | N | O3 | 10 | 0.050% | 0 | 0.055% |
| 88 | N | O12 | 1 | 0.005% | 0 | |
| 89 | N | O12 | 1 | 0.005% | 0 | 0.550% |
| 89 | O | O3 | 8 | 0.040% | 0 | |
| 89 | O | O6 | 66 | 0.330% | 0 | |
| 89 | O | O7 | 26 | 0.130% | 0 | |
| 89 | O | O8 | 6 | 0.030% | 0 | |
| 89 | O | O11 | 2 | 0.010% | 0 | |
| 89 | O | O12 | 1 | 0.005% | 0 | |
| 90 | O | O11 | 1 | 0.005% | 0 | 0.005% |
| 91 | O | O3 | 8 | 0.040% | 0 | 0.040% |
| 92 | N | O3 | 1 | 0.005% | 0 | 0.005% |
| 111 | NH1 | O13 | 3 | 0.015% | 0 | 0.045% |
| 111 | NH2 | O13 | 6 | 0.030% | 0 | |

Hydrogen bonds observed during last 400ns of MD simulations between *Mc* FKBP12 and FK506

| Residue # | residue atom | FK506 atom | # of times hbond observed | frequency of hbond's existence | Z-SCORE | Overall residue frequency of hbonds |
|---|---|---|---|---|---|---|
| 27 | OH | O4 | 8 | 0.040% | 0 | |
| 27 | OH | O5 | 277 | 1.385% | 0 | 1.590% |
| 27 | OH | O6 | 33 | 0.165% | 0 | |
| 29 | N | O4 | 1 | 0.005% | 0 | 0.020% |
| 29 | N | O6 | 3 | 0.015% | 0 | |
| 37 | O | O6 | 20 | 0.100% | 0 | 0.100% |
| 38 | N | O4 | 2 | 0.010% | 0 | |
| 38 | N | O6 | 155 | 0.775% | 0 | |
| 38 | N | O7 | 23 | 0.115% | 0 | |
| 38 | OD1 | O4 | 17 | 0.085% | 0 | |
| 38 | OD1 | O5 | 175 | 0.875% | 0 | |
| 38 | OD1 | O6 | 371 | 1.855% | 0 | |
| 38 | OD1 | O7 | 1 | 0.005% | 0 | 8.953% |
| 38 | OD2 | O4 | 14 | 0.070% | 0 | |
| 38 | OD2 | O5 | 203 | 1.015% | 0 | |
| 38 | OD2 | O6 | 342 | 1.710% | 0 | |
| 38 | OD2 | O7 | 1 | 0.005% | 0 | |
| 38 | O | O4 | 48 | 0.240% | 0 | |
| 38 | O | O5 | 10 | 0.050% | 0 | |
| 38 | O | O6 | 429 | 2.144% | 0 | |
| 39 | OG | O6 | 4 | 0.020% | 0 | 0.020% |
| 43 | NE | O6 | 7 | 0.035% | 0 | |
| 43 | NH1 | O7 | 30 | 0.150% | 0 | |
| 43 | NH1 | O8 | 8 | 0.040% | 0 | 1.355% |
| 43 | NH2 | O6 | 4 | 0.020% | 0 | |
| 43 | NH2 | O7 | 160 | 0.800% | 0 | |
| 43 | NH2 | O8 | 62 | 0.310% | 0 | |
| 50 | OG1 | O10 | 6 | 0.030% | 0 | 0.035% |
| 50 | OG1 | O2 | 1 | 0.005% | 0 | |
| 53 | O | O10 | 4 | 0.020% | 0 | 0.020% |
| 54 | N | O11 | 3 | 0.015% | 0 | |
| 54 | O | O11 | 5 | 0.025% | 0 | 0.440% |
| 54 | O | O12 | 80 | 0.400% | 0 | |
| 55 | NE2 | O10 | 196 | 0.980% | 0 | |
| 55 | NE2 | O11 | 48 | 0.240% | 0 | |
| 55 | NE2 | O12 | 32 | 0.160% | 0 | |
| 55 | NE2 | O2 | 1 | 0.005% | 0 | |
| 55 | N | O10 | 25 | 0.125% | 0 | |
| 55 | OE1 | O10 | 315 | 1.575% | 0 | |

Hydrogen bonds observed during last 400ns of MD simulations between *Mc* FKBP12 and FK506

| Residue # | residue atom | FK506 atom | # of times hbond observed | frequency of hbond's existence | Z-SCORE | Overall residue frequency of hbonds |
|---|---|---|---|---|---|---|
| 55 | OE1 | O11 | 7 | 0.035% | 0 | 35.276% |
| 55 | OE1 | O12 | 133 | 0.665% | 0 | |
| 55 | OE1 | O9 | 2 | 0.010% | 0 | |
| 55 | O | O1 | 1 | 0.005% | 0 | |
| 55 | O | O10 | 5978 | 29.883% | 4 | |
| 55 | O | O11 | 6 | 0.030% | 0 | |
| 55 | O | O12 | 2 | 0.010% | 0 | |
| 55 | O | O2 | 311 | 1.555% | 0 | |
| 56 | N | O10 | 15 | 0.075% | 0 | 0.120% |
| 56 | O | O11 | 7 | 0.035% | 0 | |
| 56 | O | O12 | 2 | 0.010% | 0 | |
| 57 | N | O2 | 13379 | 66.878% | 10 | 66.893% |
| 57 | N | O3 | 3 | 0.015% | 0 | |
| 58 | NZ | O11 | 2 | 0.010% | 0 | 0.010% |
| 60 | NE1 | O3 | 3049 | 15.241% | 2 | 15.491% |
| 60 | NE1 | O4 | 50 | 0.250% | 0 | |
| 82 | O | O11 | 1 | 0.005% | 0 | 0.040% |
| 82 | O | O12 | 4 | 0.020% | 0 | |
| 82 | O | O3 | 3 | 0.015% | 0 | |
| 83 | OH | O11 | 216 | 1.080% | 0 | 2.914% |
| 83 | OH | O12 | 307 | 1.535% | 0 | |
| 83 | OH | O3 | 11 | 0.055% | 0 | |
| 83 | OH | O9 | 18 | 0.090% | 0 | |
| 83 | O | O11 | 7 | 0.035% | 0 | |
| 83 | O | O12 | 24 | 0.120% | 0 | |
| 84 | N | O11 | 696 | 3.479% | 0 | 5.224% |
| 84 | N | O12 | 159 | 0.795% | 0 | |
| 84 | O | O11 | 98 | 0.490% | 0 | |
| 84 | O | O12 | 92 | 0.460% | 0 | |
| 85 | N | O11 | 603 | 3.014% | 0 | 6.573% |
| 85 | N | O12 | 146 | 0.730% | 0 | |
| 85 | OE1 | O12 | 46 | 0.230% | 0 | |
| 85 | OE2 | O12 | 30 | 0.150% | 0 | |
| 85 | O | O11 | 41 | 0.205% | 0 | |
| 85 | O | O12 | 449 | 2.244% | 0 | |
| 86 | NE | O11 | 187 | 0.935% | 0 | |
| 86 | NE | O12 | 126 | 0.630% | 0 | |
| 86 | NH1 | O10 | 2 | 0.010% | 0 | |
| 86 | NH1 | O11 | 334 | 1.670% | 0 | |

Hydrogen bonds observed during last 400ns of MD simulations between *Mc* FKBP12 and FK506

| Residue # | residue atom | FK506 atom | # of times hbond observed | frequency of hbond's existence | Z-SCORE | Overall residue frequency of hbonds |
|---|---|---|---|---|---|---|
| 86 | NH1 | O12 | 277 | 1.385% | 0 | |
| 86 | NH2 | O10 | 3 | 0.015% | 0 | |
| 86 | NH2 | O11 | 132 | 0.660% | 0 | 14.806% |
| 86 | NH2 | O12 | 72 | 0.360% | 0 | |
| 86 | N | O11 | 489 | 2.444% | 0 | |
| 86 | N | O12 | 891 | 4.454% | 0 | |
| 86 | O | O10 | 4 | 0.020% | 0 | |
| 86 | O | O11 | 67 | 0.335% | 0 | |
| 86 | O | O12 | 378 | 1.890% | 0 | |
| 87 | N | O11 | 117 | 0.585% | 0 | |
| 87 | N | O12 | 242 | 1.210% | 0 | 4.189% |
| 87 | O | O11 | 164 | 0.820% | 0 | |
| 87 | O | O12 | 315 | 1.575% | 0 | |
| 88 | N | O10 | 4 | 0.020% | 0 | |
| 88 | N | O11 | 170 | 0.850% | 0 | |
| 88 | N | O12 | 372 | 1.860% | 0 | |
| 88 | N | O9 | 5 | 0.025% | 0 | |
| 88 | OH | O10 | 2 | 0.010% | 0 | |
| 88 | OH | O11 | 1 | 0.005% | 0 | |
| 88 | OH | O12 | 40 | 0.200% | 0 | 6.338% |
| 88 | OH | O3 | 39 | 0.195% | 0 | |
| 88 | OH | O6 | 1 | 0.005% | 0 | |
| 88 | OH | O7 | 12 | 0.060% | 0 | |
| 88 | OH | O9 | 5 | 0.025% | 0 | |
| 88 | O | O11 | 108 | 0.540% | 0 | |
| 88 | O | O12 | 509 | 2.544% | 0 | |
| 89 | N | O11 | 1 | 0.005% | 0 | |
| 89 | N | O12 | 1 | 0.005% | 0 | |
| 89 | O | O11 | 5 | 0.025% | 0 | 0.320% |
| 89 | O | O12 | 51 | 0.255% | 0 | |
| 89 | O | O6 | 4 | 0.020% | 0 | |
| 89 | O | O7 | 2 | 0.010% | 0 | |
| 90 | N | O11 | 56 | 0.280% | 0 | |
| 90 | N | O12 | 21 | 0.105% | 0 | |
| 90 | N | O7 | 2 | 0.010% | 0 | |
| 90 | O | O11 | 12 | 0.060% | 0 | 0.710% |
| 90 | O | O12 | 9 | 0.045% | 0 | |
| 90 | O | O6 | 38 | 0.190% | 0 | |
| 90 | O | O7 | 4 | 0.020% | 0 | |
| 91 | N | O11 | 6 | 0.030% | 0 | |

Hydrogen bonds observed during last 400ns of MD simulations between *Mc* FKBP12 and FK506

| Residue # | residue atom | FK506 atom | # of times hbond observed | frequency of hbond's existence | Z-SCORE | Overall residue frequency of hbonds |
|---|---|---|---|---|---|---|
| 91 | N | O12 | 1 | 0.005% | 0 | 0.220% |
| 91 | N | O6 | 18 | 0.090% | 0 | |
| 91 | N | O7 | 8 | 0.040% | 0 | |
| 91 | O | O12 | 1 | 0.005% | 0 | |
| 91 | O | O6 | 3 | 0.015% | 0 | |
| 91 | O | O7 | 7 | 0.035% | 0 | |
| 92 | N | O4 | 2 | 0.010% | 0 | 0.050% |
| 92 | N | O6 | 8 | 0.040% | 0 | |

Hydrogen bonds observed during last 400ns of MD simulations between *Mc* FKBP12 and APX879

| Residue # | residue atom | APX879 atom | # of times hbond observed | frequency of hbond's existence | Z-SCORE | Overall residue frequency of hbonds |
|---|---|---|---|---|---|---|
| 22 | NZ | O13 | 15 | 0.075% | 0 | 0.075% |
| 27 | OH | O4 | 15 | 0.075% | 0 | |
| 27 | OH | O5 | 12 | 0.060% | 0 | |
| 27 | OH | O6 | 1312 | 6.558% | 1 | 14.691% |
| 27 | OH | O8 | 1599 | 7.993% | 1 | |
| 27 | OH | O13 | 1 | 0.005% | 0 | |
| 37 | O | O4 | 1 | 0.005% | 0 | 0.165% |
| 37 | O | O6 | 32 | 0.160% | 0 | |
| 38 | N | O4 | 5 | 0.025% | 0 | |
| 38 | N | O6 | 109 | 0.545% | 0 | |
| 38 | N | O8 | 4 | 0.020% | 0 | |
| 38 | OD1 | O4 | 329 | 1.645% | 0 | |
| 38 | OD1 | O5 | 1 | 0.005% | 0 | |
| 38 | OD1 | O6 | 5650 | 28.243% | 6 | |
| 38 | OD1 | O7 | 6 | 0.030% | 0 | |
| 38 | OD1 | O8 | 21 | 0.105% | 0 | |
| 38 | OD2 | O4 | 360 | 1.800% | 0 | 68.008% |
| 38 | OD2 | O5 | 3 | 0.015% | 0 | |
| 38 | OD2 | O6 | 6057 | 30.277% | 6 | |
| 38 | OD2 | O7 | 4 | 0.020% | 0 | |
| 38 | OD2 | O8 | 24 | 0.120% | 0 | |
| 38 | O | O4 | 11 | 0.055% | 0 | |
| 38 | O | O5 | 17 | 0.085% | 0 | |
| 38 | O | O6 | 948 | 4.739% | 1 | |
| 38 | O | O8 | 56 | 0.280% | 0 | |
| 43 | NE | O6 | 99 | 0.495% | 0 | |
| 43 | NE | O8 | 226 | 1.130% | 0 | |
| 43 | NE | O13 | 32 | 0.160% | 0 | |
| 43 | NH1 | N54 | 1 | 0.005% | 0 | |
| 43 | NH1 | O4 | 44 | 0.220% | 0 | |
| 43 | NH1 | O5 | 8 | 0.040% | 0 | |
| 43 | NH1 | O6 | 113 | 0.565% | 0 | |
| 43 | NH1 | O7 | 8 | 0.040% | 0 | |
| 43 | NH1 | O8 | 616 | 3.079% | 0 | |
| 43 | NH1 | O10 | 7 | 0.035% | 0 | 33.152% |
| 43 | NH1 | O13 | 147 | 0.735% | 0 | |
| 43 | NH2 | N54 | 1 | 0.005% | 0 | |
| 43 | NH2 | N55 | 1 | 0.005% | 0 | |
| 43 | NH2 | O4 | 6 | 0.030% | 0 | |
| 43 | NH2 | O5 | 74 | 0.370% | 0 | |

Hydrogen bonds observed during last 400ns of MD simulations between *Mc* FKBP12 and APX879

| **Residue #** | **residue atom** | **APX879 atom** | **# of times hbond observed** | **frequency of hbond's existence** | **Z-SCORE** | **Overall residue frequency of hbonds** |
|---|---|---|---|---|---|---|
| 43 | NH2 | O6 | 3492 | 17.456% | 3 | |
| 43 | NH2 | O7 | 62 | 0.310% | 0 | |
| 43 | NH2 | O8 | 1540 | 7.698% | 1 | |
| 43 | NH2 | O10 | 30 | 0.150% | 0 | |
| 43 | NH2 | O13 | 125 | 0.625% | 0 | |
| 48 | NE2 | O13 | 2 | 0.010% | 0 | |
| 48 | N | O13 | 1 | 0.005% | 0 | |
| 48 | OE1 | N55 | 4 | 0.020% | 0 | 0.945% |
| 48 | O | N55 | 55 | 0.275% | 0 | |
| 48 | O | O10 | 28 | 0.140% | 0 | |
| 48 | O | O13 | 99 | 0.495% | 0 | |
| 50 | N | O10 | 224 | 1.120% | 0 | |
| 50 | N | O13 | 1275 | 6.373% | 1 | |
| 50 | OG1 | N55 | 20 | 0.100% | 0 | |
| 50 | OG1 | O10 | 697 | 3.484% | 0 | 14.731% |
| 50 | OG1 | O13 | 704 | 3.519% | 0 | |
| 50 | O | O10 | 3 | 0.015% | 0 | |
| 50 | O | O13 | 24 | 0.120% | 0 | |
| 51 | N | O13 | 1 | 0.005% | 0 | |
| 51 | O | N55 | 2 | 0.010% | 0 | 0.020% |
| 51 | O | O13 | 1 | 0.005% | 0 | |
| 53 | O | O13 | 1 | 0.005% | 0 | 0.005% |
| 54 | N | O11 | 2 | 0.010% | 0 | |
| 54 | O | O10 | 13 | 0.065% | 0 | 1.305% |
| 54 | O | O11 | 121 | 0.605% | 0 | |
| 54 | O | O12 | 125 | 0.625% | 0 | |
| 55 | NE2 | N54 | 2 | 0.010% | 0 | |
| 55 | NE2 | N55 | 18 | 0.090% | 0 | |
| 55 | NE2 | O10 | 257 | 1.285% | 0 | |
| 55 | NE2 | O11 | 145 | 0.725% | 0 | |
| 55 | NE2 | O12 | 72 | 0.360% | 0 | |
| 55 | NE2 | O13 | 55 | 0.275% | 0 | |
| 55 | N | O10 | 80 | 0.400% | 0 | |
| 55 | N | O12 | 1 | 0.005% | 0 | |
| 55 | N | O13 | 1 | 0.005% | 0 | |
| 55 | OE1 | N54 | 1 | 0.005% | 0 | |
| 55 | OE1 | N55 | 688 | 3.439% | 0 | 43.769% |
| 55 | OE1 | O10 | 381 | 1.905% | 0 | |
| 55 | OE1 | O11 | 31 | 0.155% | 0 | |
| 55 | OE1 | O12 | 22 | 0.110% | 0 | |

Hydrogen bonds observed during last 400ns of MD simulations between *Mc* FKBP12 and APX879

| Residue # | residue atom | APX879 atom | # of times hbond observed | frequency of hbond's existence | Z-SCORE | Overall residue frequency of hbonds |
|---|---|---|---|---|---|---|
| 55 | OE1 | O13 | 11 | 0.055% | 0 | |
| 55 | O | N55 | 61 | 0.305% | 0 | |
| 55 | O | O1 | 1 | 0.005% | 0 | |
| 55 | O | O2 | 23 | 0.115% | 0 | |
| 55 | O | O10 | 6831 | 34.147% | 7 | |
| 55 | O | O11 | 62 | 0.310% | 0 | |
| 55 | O | O12 | 10 | 0.050% | 0 | |
| 55 | O | O13 | 3 | 0.015% | 0 | |
| 56 | N | O10 | 96 | 0.480% | 0 | 0.610% |
| 56 | N | O11 | 1 | 0.005% | 0 | |
| 56 | O | O11 | 24 | 0.120% | 0 | |
| 56 | O | O12 | 1 | 0.005% | 0 | |
| 57 | N | O2 | 6047 | 30.227% | 6 | 30.232% |
| 57 | N | O10 | 1 | 0.005% | 0 | |
| 58 | N | O11 | 5 | 0.025% | 0 | 0.210% |
| 58 | N | O12 | 2 | 0.010% | 0 | |
| 58 | NZ | O11 | 3 | 0.015% | 0 | |
| 58 | NZ | O12 | 32 | 0.160% | 0 | |
| 60 | NE1 | O2 | 14 | 0.070% | 0 | 6.188% |
| 60 | NE1 | O3 | 1194 | 5.969% | 1 | |
| 60 | NE1 | O4 | 30 | 0.150% | 0 | |
| 79 | NE2 | O12 | 1 | 0.005% | 0 | 0.005% |
| 81 | O | O12 | 1 | 0.005% | 0 | 0.005% |
| 82 | O | O3 | 199 | 0.995% | 0 | 1.215% |
| 82 | O | O4 | 1 | 0.005% | 0 | |
| 82 | O | O11 | 16 | 0.080% | 0 | |
| 82 | O | O12 | 27 | 0.135% | 0 | |
| 83 | OH | O2 | 2 | 0.010% | 0 | 5.379% |
| 83 | OH | O3 | 710 | 3.549% | 0 | |
| 83 | OH | O5 | 2 | 0.010% | 0 | |
| 83 | OH | O7 | 27 | 0.135% | 0 | |
| 83 | OH | O11 | 33 | 0.165% | 0 | |
| 83 | OH | O12 | 115 | 0.575% | 0 | |
| 83 | OH | O13 | 1 | 0.005% | 0 | |
| 83 | O | O11 | 57 | 0.285% | 0 | |
| 83 | O | O12 | 129 | 0.645% | 0 | |
| 84 | N | O11 | 240 | 1.200% | 0 | 3.424% |
| 84 | N | O12 | 176 | 0.880% | 0 | |
| 84 | O | O11 | 29 | 0.145% | 0 | |
| 84 | O | O12 | 240 | 1.200% | 0 | |

Hydrogen bonds observed during last 400ns of MD simulations between *Mc* FKBP12 and APX879

| Residue # | residue atom | APX879 atom | # of times hbond observed | frequency of hbond's existence | Z-SCORE | Overall residue frequency of hbonds |
|---|---|---|---|---|---|---|
| 85 | N | O11 | 26 | 0.130% | 0 | |
| 85 | N | O12 | 87 | 0.435% | 0 | |
| 85 | OE1 | O11 | 2 | 0.010% | 0 | |
| 85 | OE1 | O12 | 150 | 0.750% | 0 | 2.224% |
| 85 | OE2 | O11 | 1 | 0.005% | 0 | |
| 85 | OE2 | O12 | 163 | 0.815% | 0 | |
| 85 | O | O11 | 4 | 0.020% | 0 | |
| 85 | O | O12 | 12 | 0.060% | 0 | |
| 86 | NE | O11 | 42 | 0.210% | 0 | |
| 86 | NE | O12 | 120 | 0.600% | 0 | |
| 86 | NE | O13 | 2 | 0.010% | 0 | |
| 86 | NH1 | N54 | 1 | 0.005% | 0 | |
| 86 | NH1 | N55 | 1 | 0.005% | 0 | |
| 86 | NH1 | O11 | 86 | 0.430% | 0 | |
| 86 | NH1 | O12 | 213 | 1.065% | 0 | |
| 86 | NH1 | O13 | 10 | 0.050% | 0 | |
| 86 | NH2 | O11 | 52 | 0.260% | 0 | 6.983% |
| 86 | NH2 | O12 | 126 | 0.630% | 0 | |
| 86 | NH2 | O13 | 7 | 0.035% | 0 | |
| 86 | N | O11 | 38 | 0.190% | 0 | |
| 86 | N | O12 | 454 | 2.269% | 0 | |
| 86 | O | O10 | 4 | 0.020% | 0 | |
| 86 | O | O11 | 65 | 0.325% | 0 | |
| 86 | O | O12 | 175 | 0.875% | 0 | |
| 86 | O | O13 | 1 | 0.005% | 0 | |
| 87 | N | O11 | 110 | 0.550% | 0 | |
| 87 | N | O12 | 76 | 0.380% | 0 | |
| 87 | O | N55 | 6 | 0.030% | 0 | |
| 87 | O | O10 | 2 | 0.010% | 0 | 2.644% |
| 87 | O | O11 | 103 | 0.515% | 0 | |
| 87 | O | O12 | 231 | 1.155% | 0 | |
| 87 | O | O13 | 1 | 0.005% | 0 | |
| 88 | N | O10 | 505 | 2.524% | 0 | |
| 88 | N | O11 | 19 | 0.095% | 0 | |
| 88 | N | O12 | 144 | 0.720% | 0 | |
| 88 | N | O13 | 3 | 0.015% | 0 | |
| 88 | OH | N55 | 3 | 0.015% | 0 | |
| 88 | OH | O3 | 1 | 0.005% | 0 | |
| 88 | OH | O4 | 1 | 0.005% | 0 | |
| 88 | OH | O5 | 1 | 0.005% | 0 | |

Hydrogen bonds observed during last 400ns of MD simulations between *Mc* FKBP12 and APX879

| Residue # | residue atom | APX879 atom | # of times hbond observed | frequency of hbond's existence | Z-SCORE | Overall residue frequency of hbonds |
|---|---|---|---|---|---|---|
| 88 | OH | O6 | 11 | 0.055% | 0 | 6.453% |
| 88 | OH | O7 | 215 | 1.075% | 0 | |
| 88 | OH | O8 | 1 | 0.005% | 0 | |
| 88 | OH | O11 | 61 | 0.305% | 0 | |
| 88 | OH | O12 | 131 | 0.655% | 0 | |
| 88 | OH | O13 | 2 | 0.010% | 0 | |
| 88 | O | O7 | 2 | 0.010% | 0 | |
| 88 | O | O10 | 2 | 0.010% | 0 | |
| 88 | O | O11 | 42 | 0.210% | 0 | |
| 88 | O | O12 | 147 | 0.735% | 0 | |
| 89 | N | O11 | 1 | 0.005% | 0 | 1.190% |
| 89 | O | O7 | 7 | 0.035% | 0 | |
| 89 | O | O11 | 12 | 0.060% | 0 | |
| 89 | O | O12 | 218 | 1.090% | 0 | |
| 90 | N | O11 | 85 | 0.425% | 0 | 0.600% |
| 90 | N | O12 | 6 | 0.030% | 0 | |
| 90 | O | O7 | 4 | 0.020% | 0 | |
| 90 | O | O12 | 25 | 0.125% | 0 | |
| 91 | N | O7 | 79 | 0.395% | 0 | 1.105% |
| 91 | N | O8 | 33 | 0.165% | 0 | |
| 91 | N | O11 | 23 | 0.115% | 0 | |
| 91 | N | O12 | 21 | 0.105% | 0 | |
| 91 | O | O6 | 18 | 0.090% | 0 | |
| 91 | O | O7 | 32 | 0.160% | 0 | |
| 91 | O | O11 | 1 | 0.005% | 0 | |
| 91 | O | O12 | 14 | 0.070% | 0 | |
| 92 | N | O12 | 6 | 0.030% | 0 | 0.060% |
| 92 | O | O7 | 6 | 0.030% | 0 | |

**Table S4. Atom index of FK506**

| Index | Type | Atoms | Index | Type | Atoms |
|---|---|---|---|---|---|
| 1 | C | C1 | 32 | CH2 | C33/H331/H332 |
| 2 | CH | C2/H2 | 33 | CH2 | C34/H341/H342 |
| 3 | CH2 | C3/H31/H32 | 34 | CH3 | C35/H351/H352/H353 |
| 4 | CH2 | C4/H41/H42 | 35 | CH3 | C36/H361/H362/H363 |
| 5 | CH2 | C5/H51/H52 | 36 | CH3 | C37/H371/H372/H373 |
| 6 | CH2 | C6/H61/H62 | 37 | CH2 | C38/H381/H382 |
| 7 | C | C8 | 38 | CH | C39/H39 |
| 8 | C | C9 | 39 | CH2 | C40/H401/H402 |
| 9 | C | C10 | 40 | CH3 | C41/H411/H412/H413 |
| 10 | CH | C11/H11 | 41 | CH3 | C42/H421/H422/H423 |
| 11 | CH2 | C12/H121/H122 | 42 | CH3 | C43/H431/H432/H433 |
| 12 | CH | C13/H13 | 43 | CH3 | C44/H441/H442/H443 |
| 13 | CH | C14/H14 | 44 | CH3 | C45/H451/H452/H453 |
| 14 | CH | C15/H15 | 45 | ---- | ---- |
| 15 | CH2 | C16/H161/H162 | 46 | ---- | ---- |
| 16 | CH | C17/H17 | 47 | N | N7 |
| 17 | CH2 | C18/H181/H182 | 48 | O | O9 |
| 18 | C | C19 | 49 | ---- | ---- |
| 19 | CH | C20/H20 | 50 | O | O1 |
| 20 | CH | C21/H21 | 51 | O | O2 |
| *21 | C | C22 | 52 | O | O3 |
| 22 | CH2 | C23/H231/H232 | 53 | O | O4 |
| 23 | CH | C24/H24 | 54 | O | O5 |
| 24 | CH | C25/H25 | 55 | OH | O6/H6 |
| 25 | CH | C26/H26 | 56 | O | O7 |
| 26 | C | C27 | 57 | O | O8 |
| 27 | CH | C28/H28 | 58 | OH | O10/H10 |
| 28 | CH | C29/H29 | 59 | O | O11 |
| 29 | CH2 | C30/H301/H302 | 60 | OH | O12/H11 |
| 30 | CH | C31/H7 | 61 | ---- | ---- |
| 31 | CH | C32/H8 | | | |

*Site of modification

**Table S5. Atom index of APX879**

| Index | Type | Atoms | Index | Type | Atoms |
|---|---|---|---|---|---|
| 1 | C | C1 | 32 | CH2 | C33/H26/H53 |
| 2 | CH | C2/H2 | 33 | CH2 | C34/H27/H54 |
| 3 | CH2 | C3/H3/H44 | 34 | CH3 | C35/H28/H55/H66 |
| 4 | CH2 | C4/H4/H45 | 35 | CH3 | C36/H29/H56/H67 |
| 5 | CH2 | C5/H5/H46 | 36 | CH3 | C37/H30/H57/H68 |
| 6 | CH2 | C6/H6/H47 | 37 | CH2 | C38/H31/H58 |
| 7 | C | C8 | 38 | CH | C39/H32 |
| 8 | C | C9 | 39 | CH2 | C40/H33/H59 |
| 9 | C | C10 | 40 | CH3 | C41/H34/H60/H69 |
| 10 | CH | C11/H7 | 41 | CH3 | C42/H35/H61/H70 |
| 11 | CH2 | C12/H8/H48 | 42 | CH3 | C43/H36/H62/H71 |
| 12 | CH | C13/H9 | 43 | CH3 | C44/H37/H63/H73 |
| 13 | CH | C14/H10 | 44 | CH3 | C45/H38/H64/H73 |
| 14 | CH | C15/H11 | 45 | C | C60 |
| 15 | CH2 | C16/H12/H49 | 46 | CH3 | C61/H01/H43/H65 |
| 16 | CH | C17/H13 | 47 | N | N7 |
| 17 | CH2 | C18/H14/H50 | 48 | N | N54 |
| 18 | C | C19 | 49 | NH | N55/H40 |
| 19 | CH | C20/H5 | 50 | O | O1 |
| 20 | CH | C21/H16 | 51 | O | O2 |
| *21 | C | C22 | 52 | O | O3 |
| 22 | CH2 | C23/H17/H51 | 53 | O | O4 |
| 23 | CH | C24/H18 | 54 | O | O5 |
| 24 | CH | C25/H19 | 55 | OH | O6/H39 |
| 25 | CH | C26/H20 | 56 | O | O7 |
| 26 | C | C27 | 57 | O | O8 |
| 27 | CH | C28/H21 | 58 | OH | O10/H41 |
| 28 | CH | C29/H22 | 59 | O | O11 |
| 29 | CH2 | C30/H23/H52 | 60 | OH | O12/H42 |
| 30 | CH | C31/H24 | 61 | O | O13 |
| 31 | CH | C32/H25 | | | |

*Site of modification

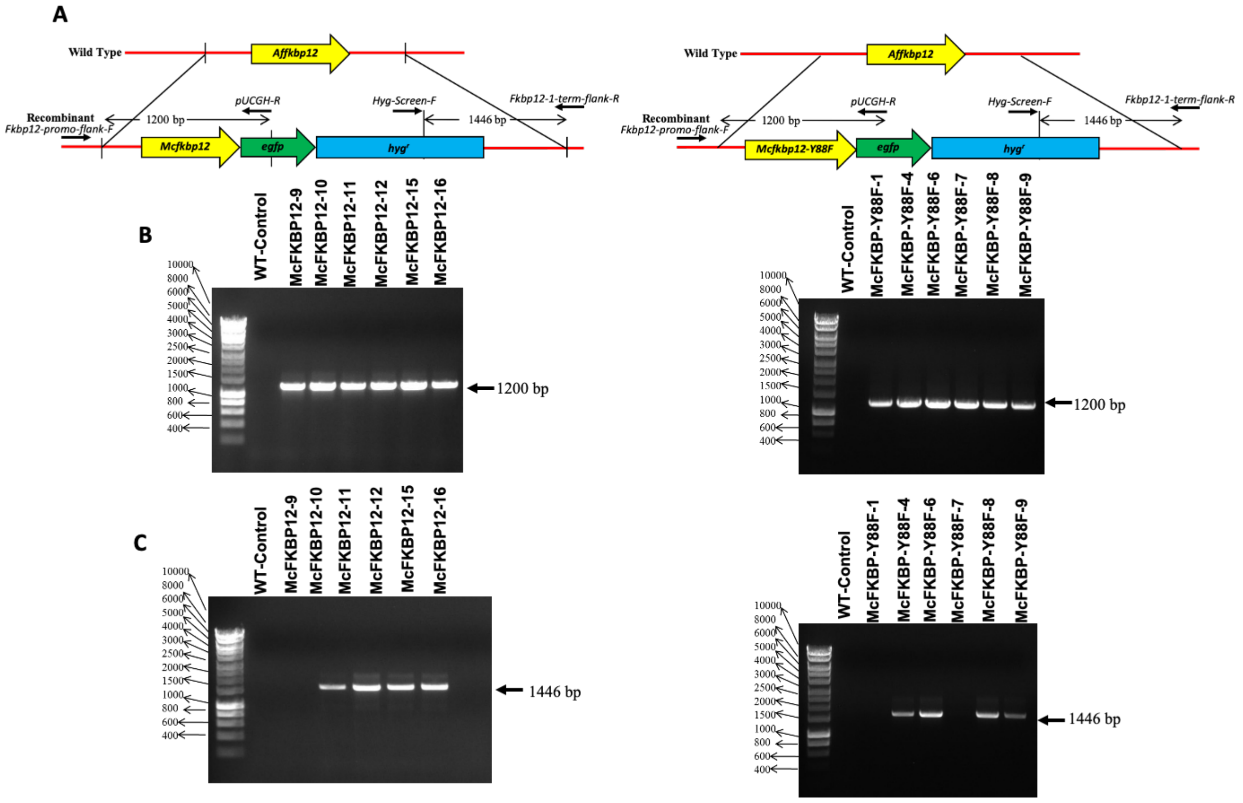


**Figure S1. Schematic of genomic locus of *Aspergillus fumigatus* expressing *Mucor circinelloides* FKBP12 and FKBP12-Y88F constructs.** (A) Schematic representation of the genomic locus of the wild-type *Affkbp12* and the recombinant strain expressing *Mcfkbp12* or *Mcfkbp12-Y88F*. The entire *A. fumigatus fkbp12* gene was replaced with the *Mcfkbp12* or *Mcfkbp12-Y88F* codon optimized DNA fused to the *gfp* sequence at its C-terminus using the hygromycin B resistance marker gene by homologous recombination. (B, C) PCR analysis for the verification of the proper integration of the *Mcfkbp12-gfp* and *Mcfkbp12-Y88F-gfp* constructs at the *Affkbp12* native locus. Primers Hyg-Screen-F and Fkbp12-term-flank-R (indicated by arrows) were used to amplify the 1446 bp PCR fragment from 6 recombinant strains. Primers Fkbp12-1-promo-flank-F and pUCGH-R (indicated by arrows) were used to amplify the 1200 bp PCR fragment from 6 recombinant strains. Strains 11, 12, 15 *Mcfkbp12-gfp* and Strains 4, 6, 9 *Mcfkbp12-Y88F-gfp* were selected for all the susceptibility assays.

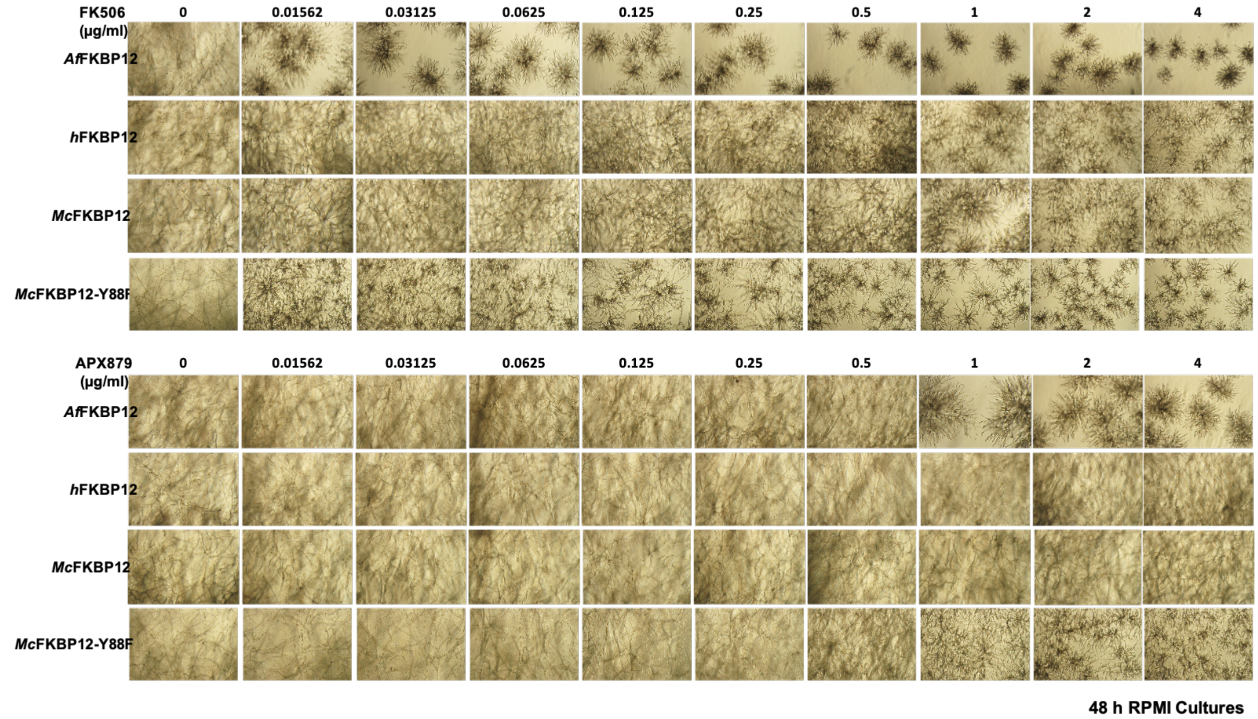


**Figure S2. Expression of human and *M. circinelloides* FKBP12 in *A. fumigatus*.** Strain expressing *h*FKBP12-GFP, *Mc*FKBP12-GFP, and *Mc*FKBP12-Y88F-GFP were cultured in RPMI liquid medium in the absence or presence of FK506 (top) and APX879 (bottom) (concentration ranging between 0 and 4 µg/mL) for 48 h. Note the resistance of *h*FKBP12 and *Mc*FKBP12 expressing strain to FK506 (0.015 µg/mL) and APX879 (1 µg/mL) in comparison to the WT strain.

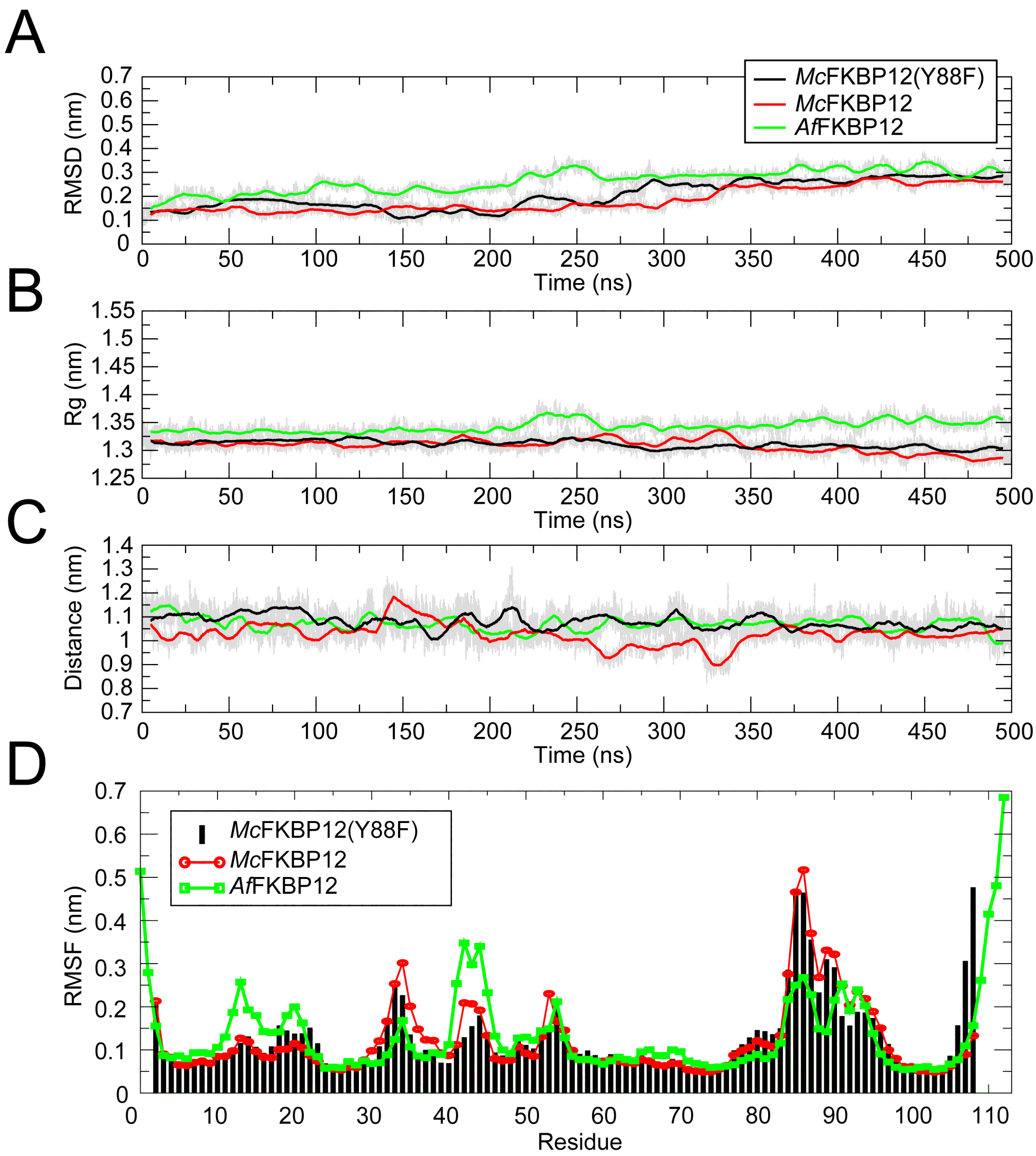


**Figure S3. Analysis of *Mc*FKBP12(Y88F)-FK506 MD simulation stability.** Measurement of (A) Cα-RMSD, (B) Radius of gyration (Rg), (C) Center of mass (COM – the 3D point of mass balance for each monomer) between the protein and FK506 and (D) RMSF values during the simulation on a per residue basis.

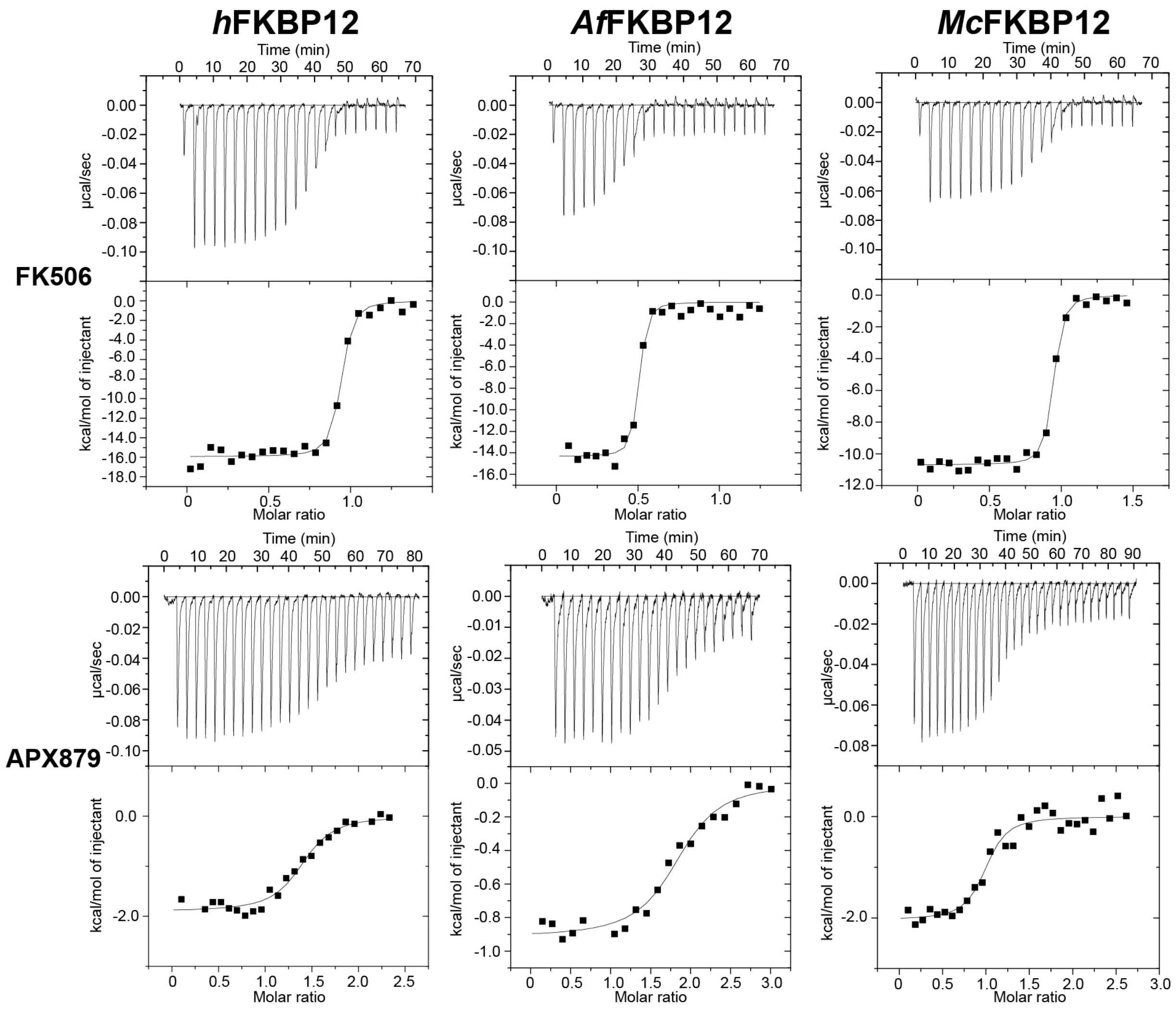


**Figure S4. The calorimetric titration of the human, *A. fumigatus,* and *M. circinelloides* FKBP12 proteins with FK506 and APX879.** Each peak of the heat pulse data (top part of each panel) represents the injection of 6µL of 25µM protein solution into 2µM of FK506 or 8µL of 150µM APX879 into 10µM protein solution performed at 25°C in 50 mM sodium phosphate, 50mM sodium chloride pH 7.0. The lower panel represents the integrated heat changes upon binding (kcal/mol) corrected for the protein and ligand heat of dilution. Fitting was performed using the MicroCal Origin software. See **Table 1** for the thermodynamic constants extracted from triplicate data.

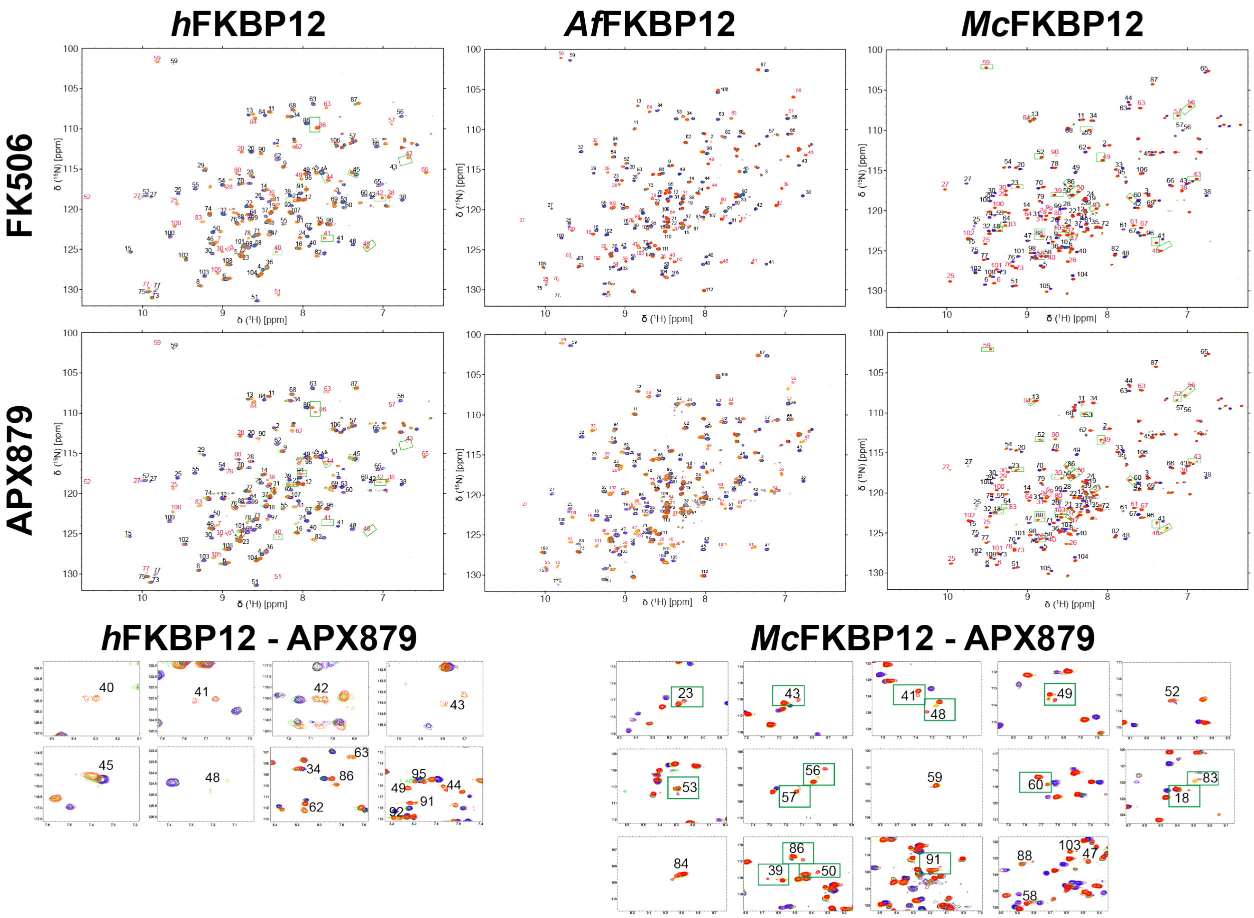


**Figure S5. NMR titrations of FK506 and APX879 in the human, *A. fumigatus* and *M. circinelloides* FKBP12 proteins.** Overlay of the $^{15}$N-HSQC titration points of FK506 and APX879 at 0:1 (black), 0.2:1 (purple), 0.4:1 (blue), 0.6:1 (green), 0.8:1 (yellow), 1.0:1 (orange) and 2.0:1 (red) molar equivalent of ligand to the human, *A. fumigatus* and *M. circinelloides* FKBP12 proteins. The peak doubling (indicated by green squares in the full spectrum) at the 1.0:1 and 2.0:1 titration points of APX879 are enlarged under the full spectrum.

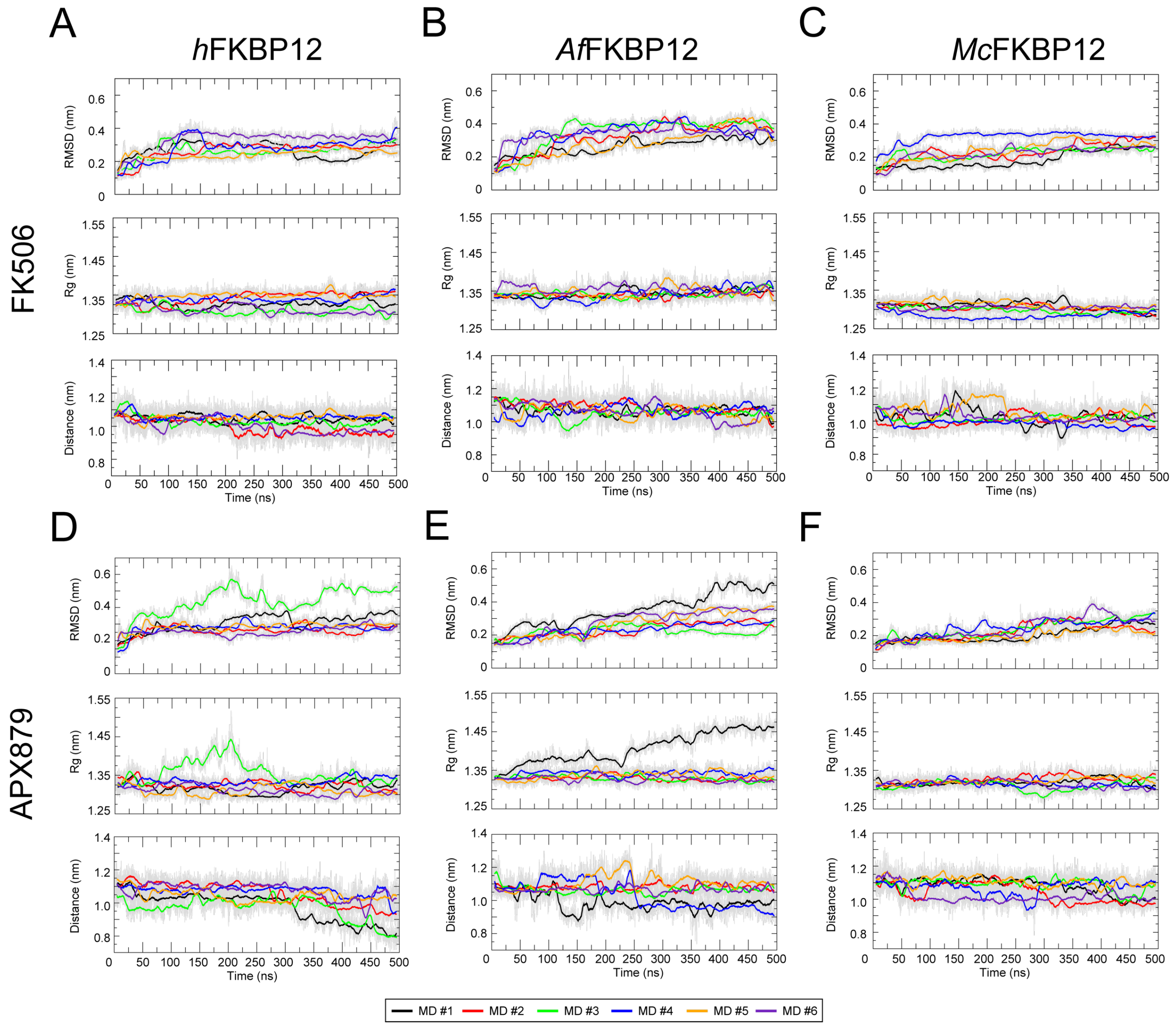


**Figure S6. Analysis of 500 ns MD simulation stability**. Panels A-C show plots of *h*FKBP12, *Af*FKBP12, and *Mc*FKBP12 bound to FK506. Panels D-F show plots of *h*FKBP12, *Af*FKBP12, and *Mc*FKBP12 bound to APX879. Top figure in each panel plots the Cα-RMSD for each MD simulation over the length of the simulation. Middle figure in each panel plots the radius of gyration (Rg) for each MD simulation over the length of the simulation. Bottom figure in each panel plots the center of mass (COM – the 3D point of mass balance for each monomer) between the protein and ligand for each MD simulation over the length of the simulation. MD sim #1 of *Af*FKBP12-APX879, #3 *h*FKBP12-APX879, and #4 *Mc*FKBP12-FK506 were removed from analysis due to unstable RMSD, Rg, and/or COM values observed during the length of the simulation.

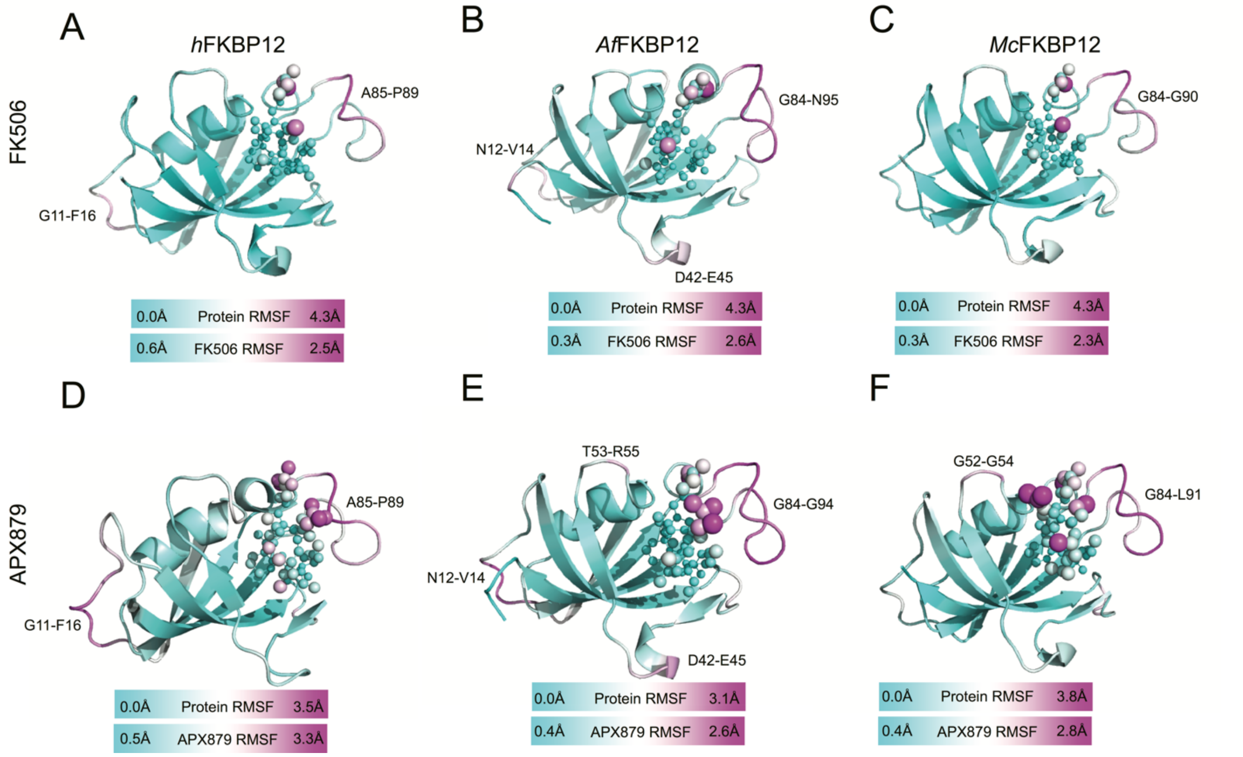


**Figure S7. RMSF comparison of the atomic positions between the X-ray crystal structures and MD simulations of human, *A. fumigatus* and *M. circinelloides* FKBP12 proteins bound to FK506 or APX879.** Cartoon representations of (A, D) *h*FKBP12, (B, E) *Af*FKBP12 and (C, F) *Mc*FKBP12 bound to (A - C) FK506 or (D - F) APX879. Residue root mean square fluctuations (RMSF) of atomic positions in the MD simulation are plotted on top of each structure and colored from cyan to white to magenta. FK506 and APX879 are shown in stick and sphere format with atomistic RSMF values with same coloring as FKBP12. Labels are provided for protein residues with the greatest difference between the X-ray crystal structures and the MD simulations.

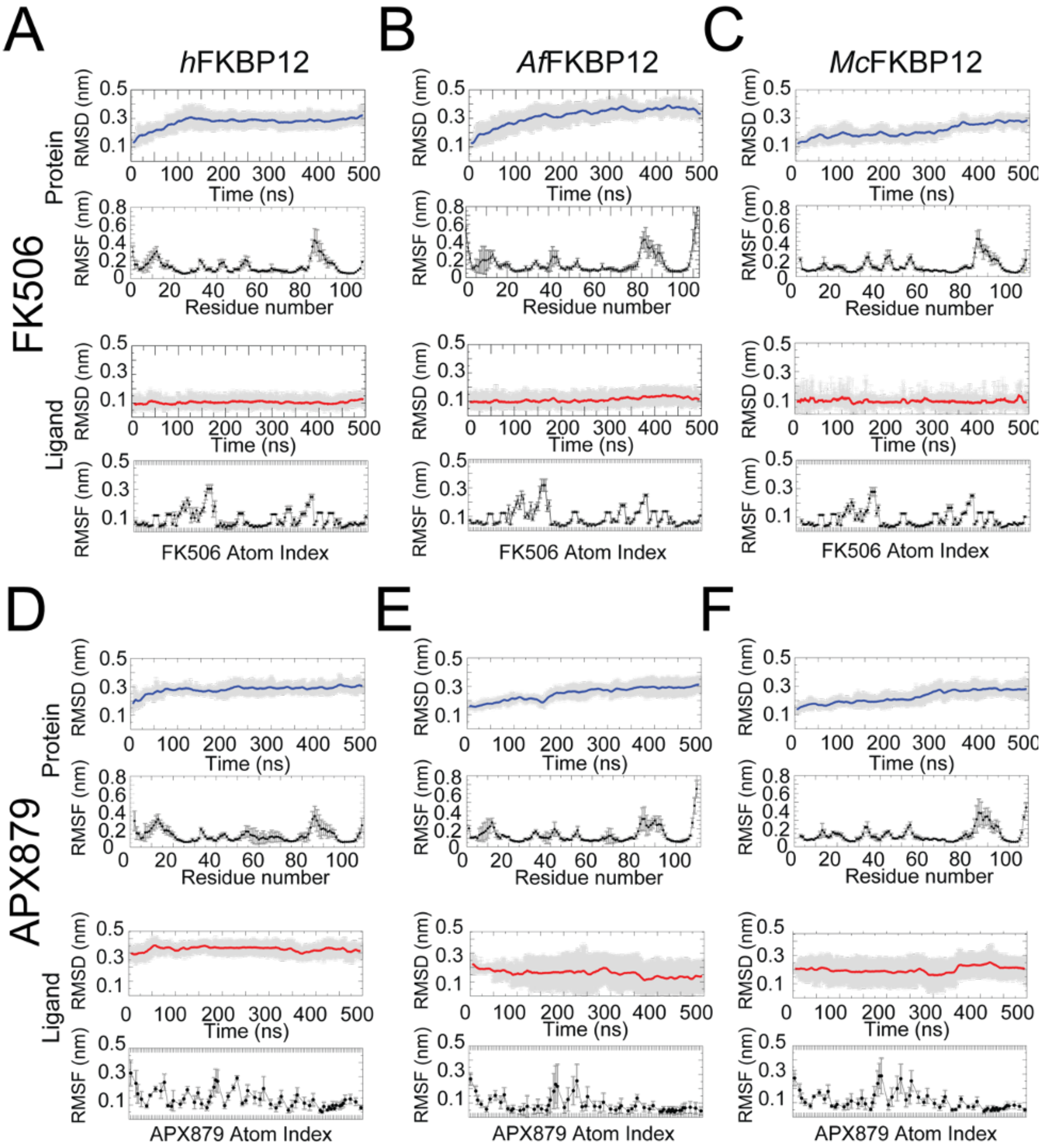


**Figure S8. RMSD and RMSF plots between the X-ray crystal characterized structures and the MD simulations.** Panels A-C show plots of *h*FKBP12, *Af*FKBP12, and *Mc*FKBP12 bound to FK506. Panels D-F show plots of *h*FKBP12, *Af*FKBP12, and *Mc*FKBP12 bound to APX879. Top two figures on each panel show the RMSD and RMSF of the FKBP12 proteins and the bottom two figures on each panel show the RMSD and RMSF of the ligand. These data were plotted on the protein and ligand structures as shown in **Fig. S7**.

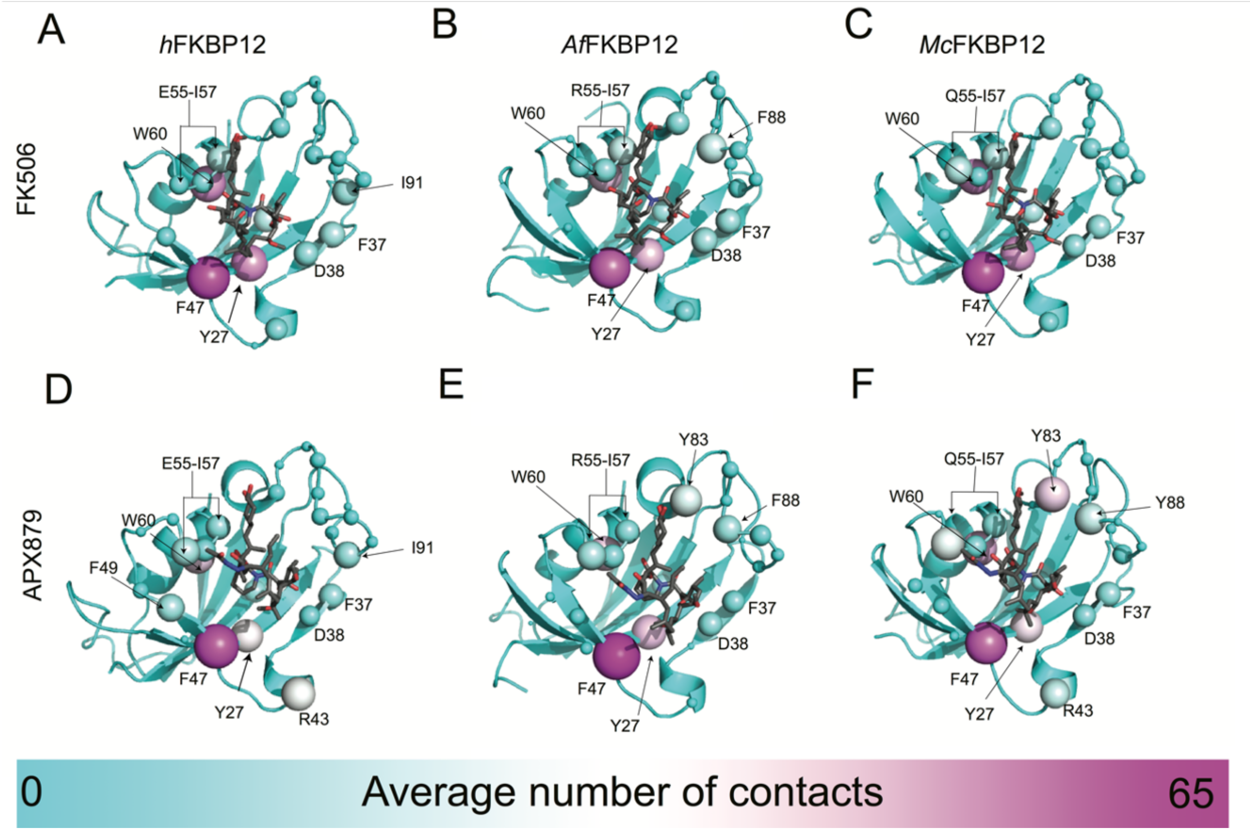


**Figure S9. MD simulation observed contacts between human, *A. fumigatus* and *M. circinelloides* FKBP12 proteins when bound to FK506 or APX879.** The average number of contacts (two non-hydrogen atoms with 4Å) that these ligands make to (A – D) *h*FKBP12, (B – E) *Af*FKBP12 and (C – F) *Mc*FKBP12 residues is plotted on top of each structure and those protein residues that have contacts to (A – C) FK506 or (D – F) APX879 are shown as spheres. The spheres size and color (small/cyan to medium/white to large/magenta) represent the number of average contacts observed. FK506 and APX879 are shown in stick format and colored according to its atom type.

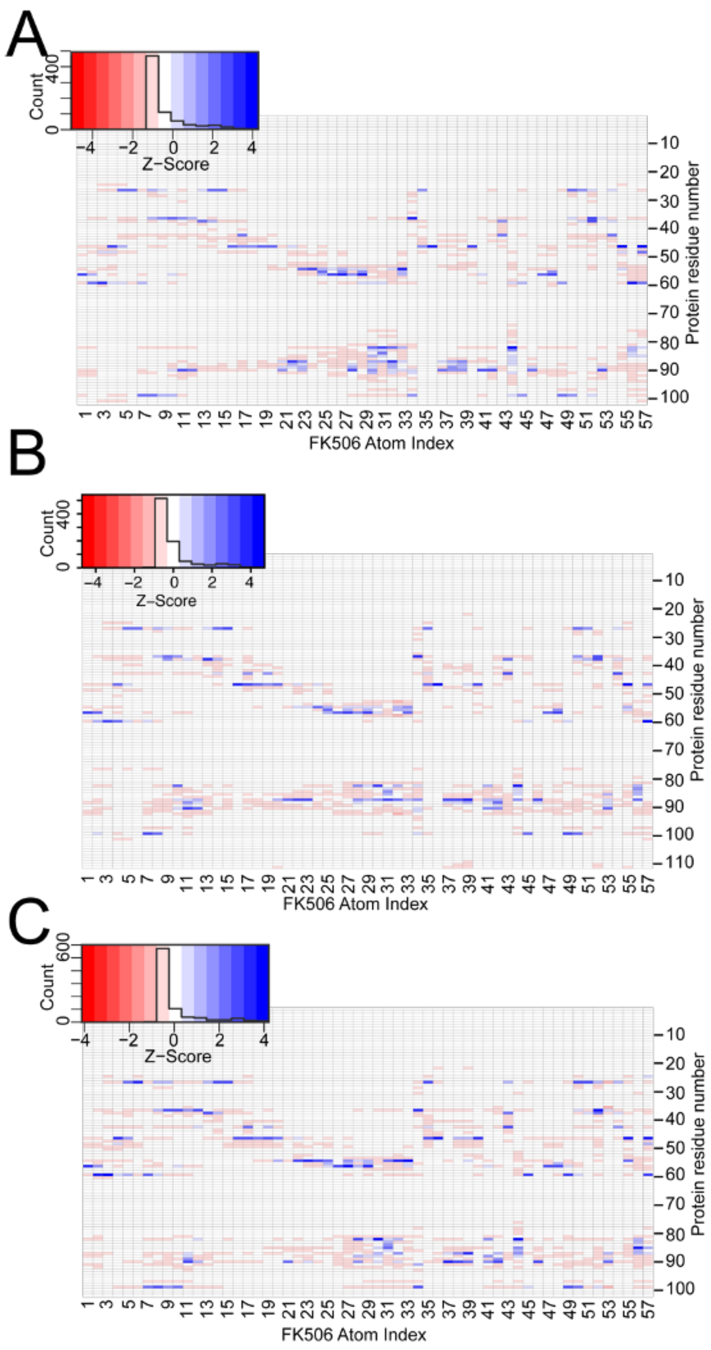
A
B
C
Count
Z-Score
FK506 Atom Index
Protein residue number

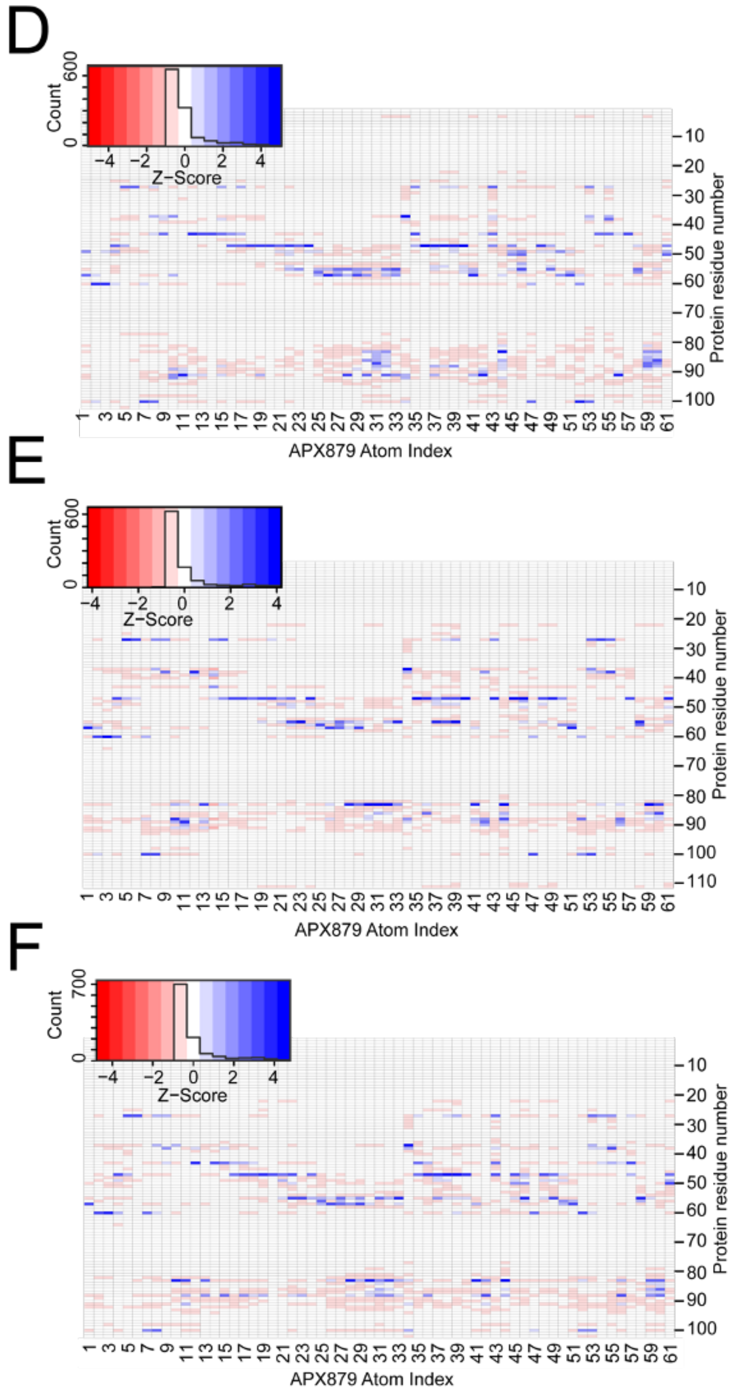


**Figure S10. Z-score matrix of protein and FK506/APX879 contacts.** Contact plots between (A and D) *h*FKBP12, (B and E) *Af*FKBP12, and (C and F) *Mc*FKBP12 and (A - C) FK506 or (D – F) APX879. The contact matrix shows the significance of the contacts observed during the simulations. These data for the protein residues were plotted on the protein structures as shown in **Figure 6 and S9**.
